# SURF: integrative analysis of a compendium of RNA-seq and CLIP-seq datasets highlights complex governing of alternative transcriptional regulation by RNA-binding proteins

**DOI:** 10.1101/2020.05.08.085316

**Authors:** Fan Chen, Sündüz Keleş

## Abstract

Advances in high-throughput profiling of RNA binding proteins (RBPs) have resulted in CLIP-seq datasets coupled with transcriptome profiling by RNA-seq. However, analysis methods that integrate both types of data are lacking. We describe SURF, Statistical Utility for RBP Functions, for integrative analysis of large collections of CLIP-seq and RNA-seq data. We demonstrate SURF’s ability to accurately detect differential alternative transcriptional regulation events and associate them to local protein-RNA interactions. We apply SURF to ENCODE RBP compendium and carry out downstream analysis with additional reference datasets. The results of this application are browsable at http://www.statlab.wisc.edu/shiny/surf/.

## 1 Background

RNA-binding proteins (RBPs) constitute a key class of regulators for post-transcriptional processes such as splicing, polyadenylation, transportation, translation, and degradation of RNA transcripts in eukaryotic organisms [1, 2]. They drive the diversity and complexity of transcriptome and proteome through gene expression regulation at the RNA level with alternative splicing (AS), alternative transcription initiation (ATI), and alternative polyadenylation (APA) [3, 4]. High-throughput sequencing-based assays that quantify protein-RNA interactions (e.g., ultraviolet cross-linking immunoprecipitation followed by high-throughput sequencing (CLIP-seq)) have recently matured and revealed a rich repertoire of RBP profiles in cell lines and tissues [5, 6, 7]. When coupled with analysis of RNA-seq experiments across multiple conditions to elucidate alternative transcriptional regulation (ATR) events of classes AS, ATI, and APA, these studies can suggest specific roles for RBPs in different types of events such as exon skipping (SE), alternative 3′ (A3SS) or 5′ (A5SS) splicing, intron retention (RI) within the AS class, alternative first exon (AFE) or alternative 5′UTR (A5U) within the ATI class, and intronic (IAP) or tandem (TAP) alternative polyadenylation within the APA class [8] (supplementary Fig. S1).

The ENCODE (ENCyclopedia of DNA elements) project [9, 10] recently generated eCLIP-seq datasets for 120 and 104 RBPs in human cells lines K562 and HepG2. These datasets were complemented with RNA-seq datasets of the same cell lines under conditions of both wild-type and short hairpin RNA (shRNA) knockdown of the individual RBPs to identify differentially regulated AS, ATI, and APA events (Fig. 1). The initial analysis by the ENCODE project revealed a large set of genomic elements interacting with RBPs and expanded the catalogue of biochemically active elements of the genome [11, 12]. In this paper, we develop a statistical framework that augments the existing analysis by specifically inferring global positioning principles of RBPs with respect to the above eight types of ATR events. Our method, named SURF for **S**tatistical **U**tility for **R**BP **F**unctions, consists of two analysis and a discovery modules. The first analysis module identifies differential ATR events within AS, ATI, and APA classes by comparing wild-type and shRNA knockdown RNA-seq data, with multiple replicates from each. The second module associates the detected differential ATR events to position-specific eCLIP-seq signals; thereby dissecting global positioning principles of protein-RNA interactions in different types of ATR events. The discovery module of SURF queries existing reference transcriptome databases, i.e., the genotype-tissue expression (GTEx) project [13] and the Cancer Genome Atlas (TCGA) program [14], to assess differential activity of ATR-specific transcript targets of RBPs between normal tissue and the relevant tumor samples.

**Fig. 1:**
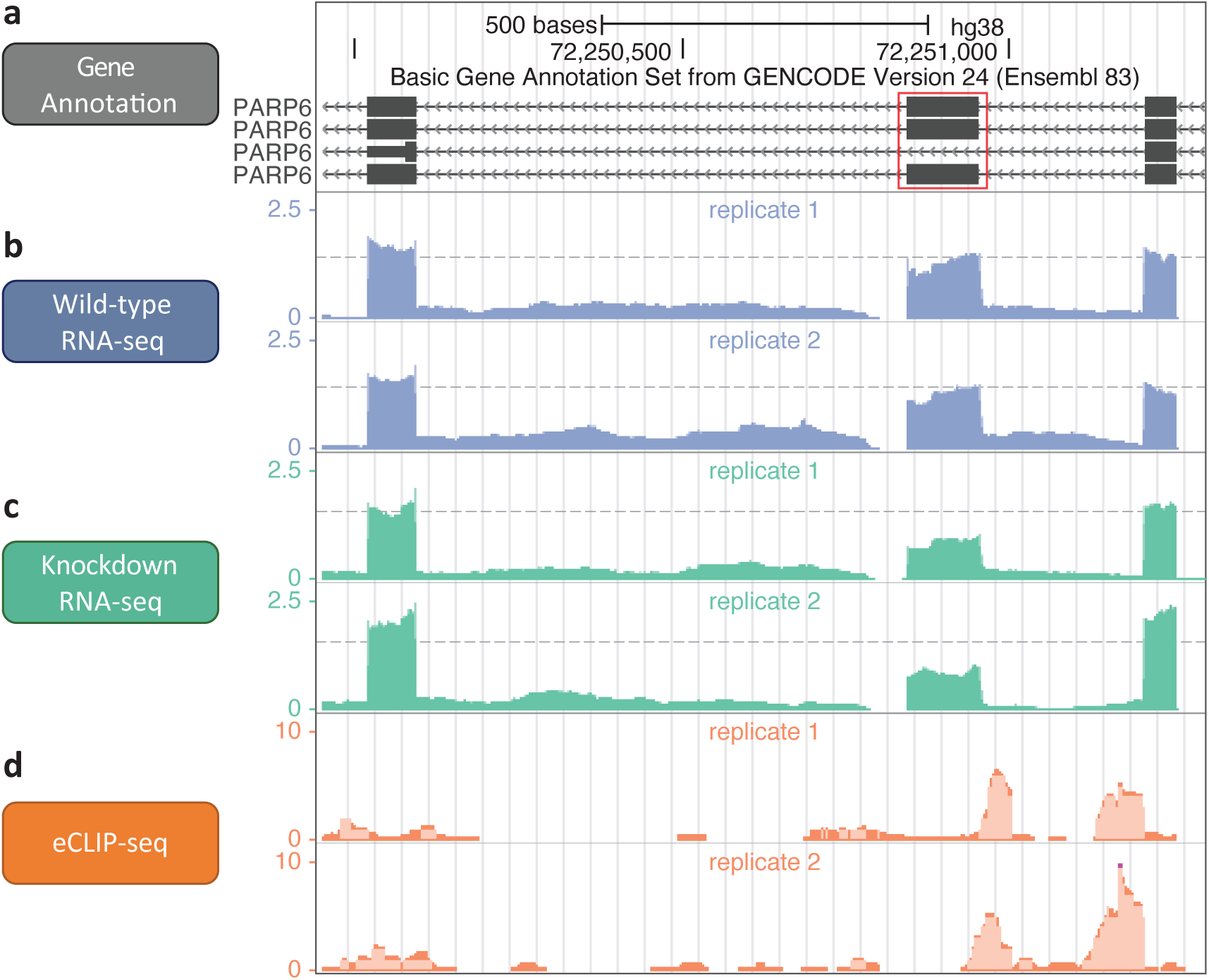
A skipped exon (SE) event in the PARP6 gene upon AQR knockdown. (a) The 18th exon (red box) of PARP6 gene model (GENCODE version 24), which resides in the minus strand. (b) The normalized read coverage in two replicates of wild-type RNA-seq. (c) The normalized read coverage in two replicates of shRNA AQR knockdown followed by RNA-seq. The 18th exon exhibits a decreased relative exon usage. (d) The normalized read coverage in two replicates of AQR eCLIP-seq, with two peaks at the upstream intron.

A number of computational tools exist for detecting differential alternative splicing from RNA-seq data across two conditions (e.g., MISO [15], MATS [16], DSGseq [17], MAJIQ [18], and rMATS [19]). These methods circumvent the challenges of direct transcriptome quantification imposed by the short read sequencing by evaluating differential ATR for each unit consisting of a cassette exon and two flanking exons. They directly test for differential ATR events of types SE, A3SS, A5SS, and RI, but not ATI and APA [20, 21, 22]. This is in contrast to methods such as Cufflinks [23], DiffSplice [24], SplicingCompass [25], and rSeqDiff [26], that operate at the isoform level. While methods that operate on isoforms test for combined effects of AS, ATI, and APA, the methods in the former category are generally more suited for investigating effects of RBPs on individual ATR events. A powerful alternative and a compromise between the two, as demonstrated by [27], is detecting alternative exon usage, irrespective of the type of ATR event the exon specifies (e.g., DEXSeq [28]). While usage-based method DEXSeq that focuses on “bins” [27] works well in conjunction with many different “unit” definitions (e.g., exons, sub-exonic bins, full length transcripts as “bins”), it does not delineate ATR event types and requires laborious inspection to classify bins into specific event types. The first module of SURF addresses this critical shortcoming by unifying differential ATR detection and generalizing DEXSeq to directly detect explicit types of AS, ATI, and APA events. Differential ATR detection by SURF exhibited superior performance to simple ways of translating DEXSeq usage inference into explicit types of AS, ATI, and APA events and rMATS, which is the only existing method that accommodates replicates within conditions and directly tests for differential ATR of AS events.

*De novo* sequence analysis of SURF-detected global positioning preferences of regions where eCLIP-seq signals associated with differential ATR event classes AS, ATI, and APA yielded sequence motif multi-modalities of several RBPs in regulating different ATR event types. Further evaluation of these regions revealed a significant enrichment of somatic mutations from both the International Cancer Genome Consortium (ICGC) [29, 30] and TCGA. In addition, analysis of SURF-identified target transcripts of RBPs for their transcriptome activity in normal GTEx versus adult acute myeloid leukemia (LAML) samples from TCGA detected a significant general shift in the activities of groups of transcripts affected by RBP-specific ATR events. These RBPs included both components of spliceosome such as AQR and SF3B4, as well as more specific RBPs such as FXR1 and PUM1. SURF is implemented in `R` [31] and is publicly available at https://github.com/keleslab/surf [32]. The results of the ENCODE data analysis are available through a shiny app at http://www.statlab.wisc.edu/shiny/surf/.

## 2 Results

### 2.1 SURF framework

SURF is designed for large-scale analysis of RBP CLIP-seq and RNA-seq data and includes analyses and discovery modules (Fig. 2). A key input to SURF is genome annotation from which SURF parses out all the eight types of ATR events, namely SE, RI, A3SS, A5SS, AFE, A5U, IAP, and TAP (Fig. 2a and supplementary Fig. S2). The parsing procedure utilizes the notion of *variable* exons (or sub-exons) [33] defined as parts of an exon that are in the gene model but are absent in one or more transcripts of the gene. Variable sub-exons of a given transcript can be clustered into mutually exclusive units, as described in Materials and Methods (supplementary Fig. S3), such that each is annotated with a unique ATR event type through a “decision tree” (supplementary Fig. S4). The analysis modules operate on these events, which we also interchangeably refer to as *event body*.

**Fig. 2:**
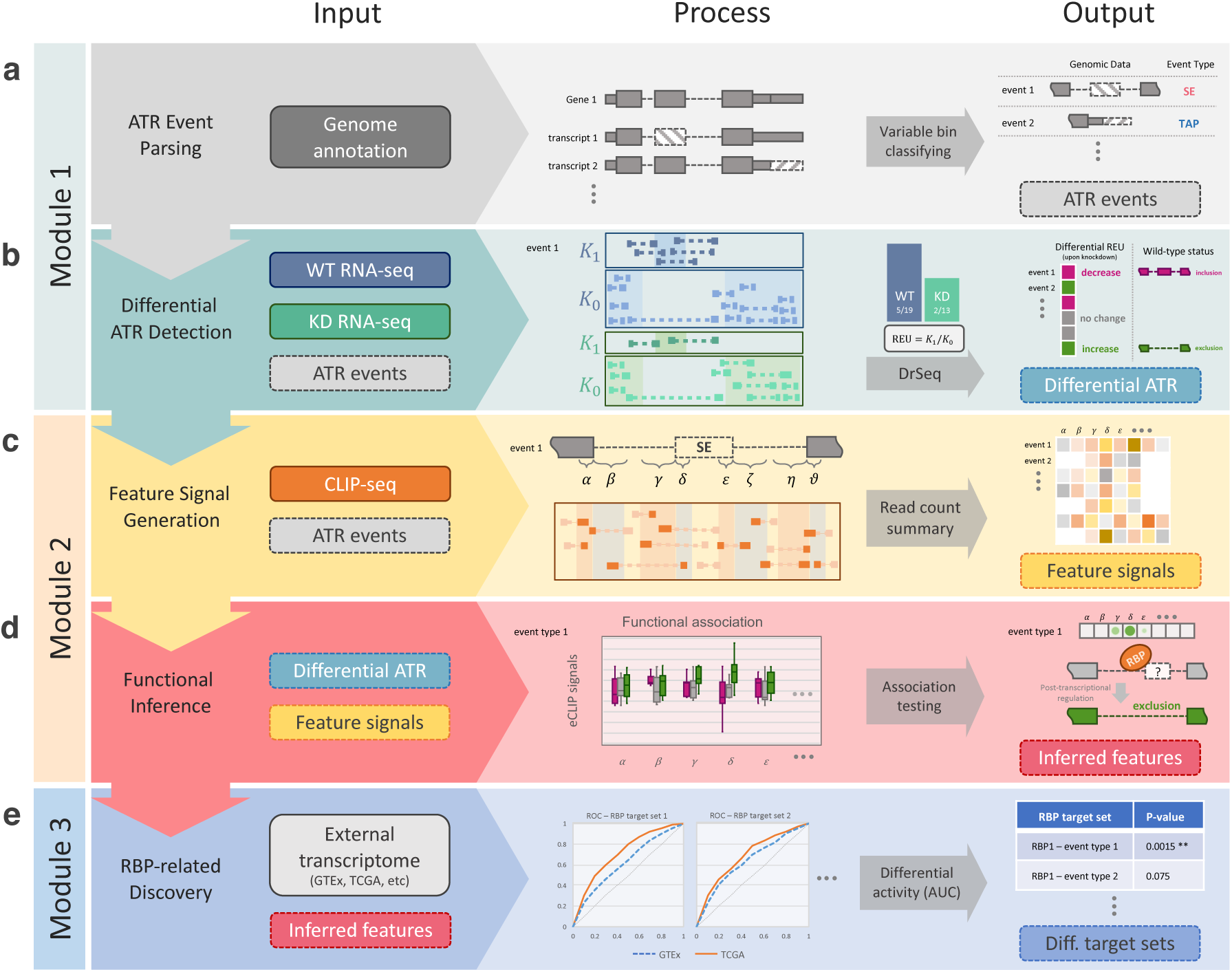
Schematic overview of SURF pipeline. (a) In the analysis module 1, the first step parses alternative transcriptional regulation (ATR) events from genome annotation. (b) SURF quantifies relative event/exon usage (REU) using RNA-seq data following RBP knockdown (with wild-type control) and performs differential REU analysis (DrSeq). As a result, it infers regulation effects of RBPs as phenotypical labels for each differential ATR event. (c) In the analysis module 2, SURF extracts location features for each ATR event and generates the feature signals using the complementary eCLIP-seq data. (d) Then, SURF models differential ATR labels by associating them with the feature signals and infers global positional preferences of RBPs. (e) In the discovery module, the inferred location features play a key role in downstream RBP-related discovery. The rank-based analysis of RBP transcript targets using external transcriptome datasets (e.g., TCGA and GTEx) discovers differential transcriptional activity through specific RBP and ATR event types.

#### 2.1.1 Analysis module 1: Detecting differential ATR events

We extended the DEXSeq framework to detect differential ATR events of all the eight types by “DrSeq” for **d**ifferential **r**egulation using RNA-**seq**. Specifically, DrSeq administers the relative exon usage (REU) to characterize any type of AS, ATI, and APA events by detecting differences between the experimental conditions, i.e., wild-type versus knockdown RNA-seq data (Fig. 2b). The REU is defined as the ratio of count of reads overlapping the event body, *K*_1_, over the count of those reads residing in the other exonic regions of the same gene, *K*_0_ and is computed as *K*_1_/*K*_0_ (Fig. 2b). For the *i*-th event in the *j*-th sample, *K*_*ijl*_ denotes either the read count on the event body (*l* = 1) or the read count of all the other exonic regions of the same gene (*l* = 0). Similar to the DEXSeq formulation [27] for exonic bins, we model *K*_*ijl*_ with a Negative Binomial (NB) distribution as

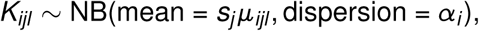

where *s*_*j*_ is the size factor of the *j*-th sample and adjusts for the sequencing depth, and *α_i_* is the dispersion parameter for event *i*. The mean parameter *µ_ijl_* is modeled by a log-link function as

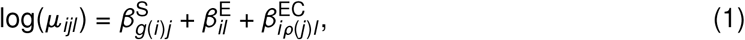

where *ρ*(*j*) is the experimental condition of sample *j* (i.e., wild-type or shRNA knockdown), and *g*(*i*) is the gene (group) from which *i*-th event originates. Here, 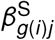 accounts for the main effect of each sample and provides a gene-specific log-scale overall read count. 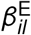 is non-zero only when *l* = 1 due to identifiability constraints; hence, it quantifies the logarithm of REU, *K*_*ij*1_/*K*_*ij*0_, for the *i*-th event across all samples regardless of experimental conditions. Finally, 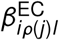 captures the effect of the experimental condition on the REU. This model is fit for each ATR event type by pooling across genes and the differential REU of each individual event is quantified by testing the REU × condition interaction term (*β*^EC^) in Equation (1) under the null hypothesis that the event has equal REU across the conditions. The overall false discovery rate (FDR) is controlled with the Benjamini-Hochberg (BH) procedure [34] by pooling over all ATR event types. SURF further quantifies the change in REU for each ATR event in the form of log2 fold change in REU. As a result, “increased REU upon RBP knockdown” indicates the exclusion of variable sub-exons under the wild-type condition. Similarly, a “decreased REU upon RBP knockdown” indicates that RBP tends to promote inclusion of variable sub-exons under the wild-type condition. We remark that this interpretation does not necessarily extend to core components of the spliceosome since these components are likely to interact with every spliced intron at some level regardless of the inclusion or exclusion outcome.

#### 2.1.2 Analysis module 2: Testing for positional associations

To infer global positioning preferences of protein-RNA interactions, SURF extracts event-specific *features* representing the potential locations of protein-RNA interactions, within and flanking the event body, as well as additional genomic regions near the constitutive exons (supplementary Fig. S5). Then, eCLIP-seq reads over these features are aggregated and normalized to adjust for the lengths of the features and the overall sequencing depth of the samples. We refer to these resulting quantities as *feature signals* (Fig. 2c). Next, SURF associates the feature signals to differential ATR events. Specifically, the ATR events are divided into three groups as “increase” (i.e., increased REU upon RBP knockdown), “decrease”, and “no change”, using the DrSeq results. Then, SURF evaluates the feature signals for their association with these labels of ATR events (Fig. 2d). By adapting a generalized linear model, SURF tests marginal association of each feature and outputs a list of genomic regions that define global positioning preferences of the RBP. This module is applied to each RBP-ATR event type combination separately.

#### 2.1.3 Discovery module: Assessing comparative transcriptional activity of RBP targets

This module leverages public transcriptome data from GTEx and TCGA to address whether transcripts harboring differential ATR events exhibit differential transcriptional activity between different clinical conditions, e.g., normal (GTEx) versus LAML (TCGA). The module first lifts over the SURF-inferred location features to transcript (or gene) level and forms the sets of transcript (or gene) targets of the RBP, while keeping them specific to ATR event types. For example, the transcript targets of SRSF1 through SE regulation (SRSF1-SE) are the set of transcripts with at least one differential ATR event of type SE regulated by SRSF1. We apply an area under the curve (AUC) approach similar to that in [35] to evaluate whether transcriptional activity of the transcript set significantly differs between the normal and tumor samples, for each combination of RBP and event type (Fig. 2e and supplementary Fig. S6). As a result, this module yields key RBPs with “increased” or “decreased” transcriptional activity in tumor samples through specific AS, ATI, and APA events.

Overall, SURF (i) leverages established appealing characteristics of DEXSeq [27] and generalizes it for detecting differential ATR events, (ii) systematically evaluates associations of differential ATR with protein-RNA interactions, and (iii) offers a unique way of further exploring large compendium of expression data with RBP and ATR event specific transcript sets.

### 2.2 SURF enables comprehensive detection of differential ATR events

We evaluated the differential ATR event detection module (DrSeq) of SURF by comparing it to rMATS [19]. The choice of rMATS among the plethora of available methods is reasoned by the fact that it accommodates biological replicates, which we have two per condition, and that it detects differential ATR for individual alternative splicing events as opposed to the differential exon usage in DEXSeq or the local splice variations in MAJIQ (supplementary Fig. S7). We additionally compared DrSeq with the following two simple strategies of mapping DEXSeq’s results to ATR event level. Specifically, we labelled an event as a differential ATR event when (i) at least one of its exonic bins (liberal strategy, DEXSeq-L) or (ii) at least half of its exonic bins (conservative strategy, DEXSeq-C) is/are identified as differentially used by DEXSeq. We evaluated the above strategies using human transcriptome-based simulated data as described in Materials and Methods. We quantified the performance of each method by its realized power and empirical FDR at gene level at target FDR levels of 0.01, 0.05, and 0.1.

The overall results, summarized in Fig. 3a and supplementary Fig. S8, delineated that all count-based methods (DrSeq and two variants of DEXSeq) achieve significantly higher TPR and lower FDR, compared to rMATS, despite detecting comparable number of events (supplementary Fig. S9). Lower power of rMATS is likely due to the fact that rMATS detects only simple splicing events (i.e., excludes ATI and API). For a limited number of simple ATR event types (e.g., SE), however, rMATS has comparable power with count-based methods (yet worse FDR, supplementary Fig. S10). Higher empirical FDR for rMATS (under various signal-to-noise levels, supplementary Fig. S11) can be largely attributed to the fact that rMATS only utilizes the reads nearby the event body and splice junctions, in contrast to the other count-based methods which account for all reads relevant to the genes (supplementary Fig. S12 and Tables S1 and S2). All count-based methods control the FDR similarly, while DrSeq achieves slightly higher power. The key difference between DrSeq and DEXSeq is mainly in their experimental units, which are ATR events and exonic bins, respectively.

**Fig. 3:**
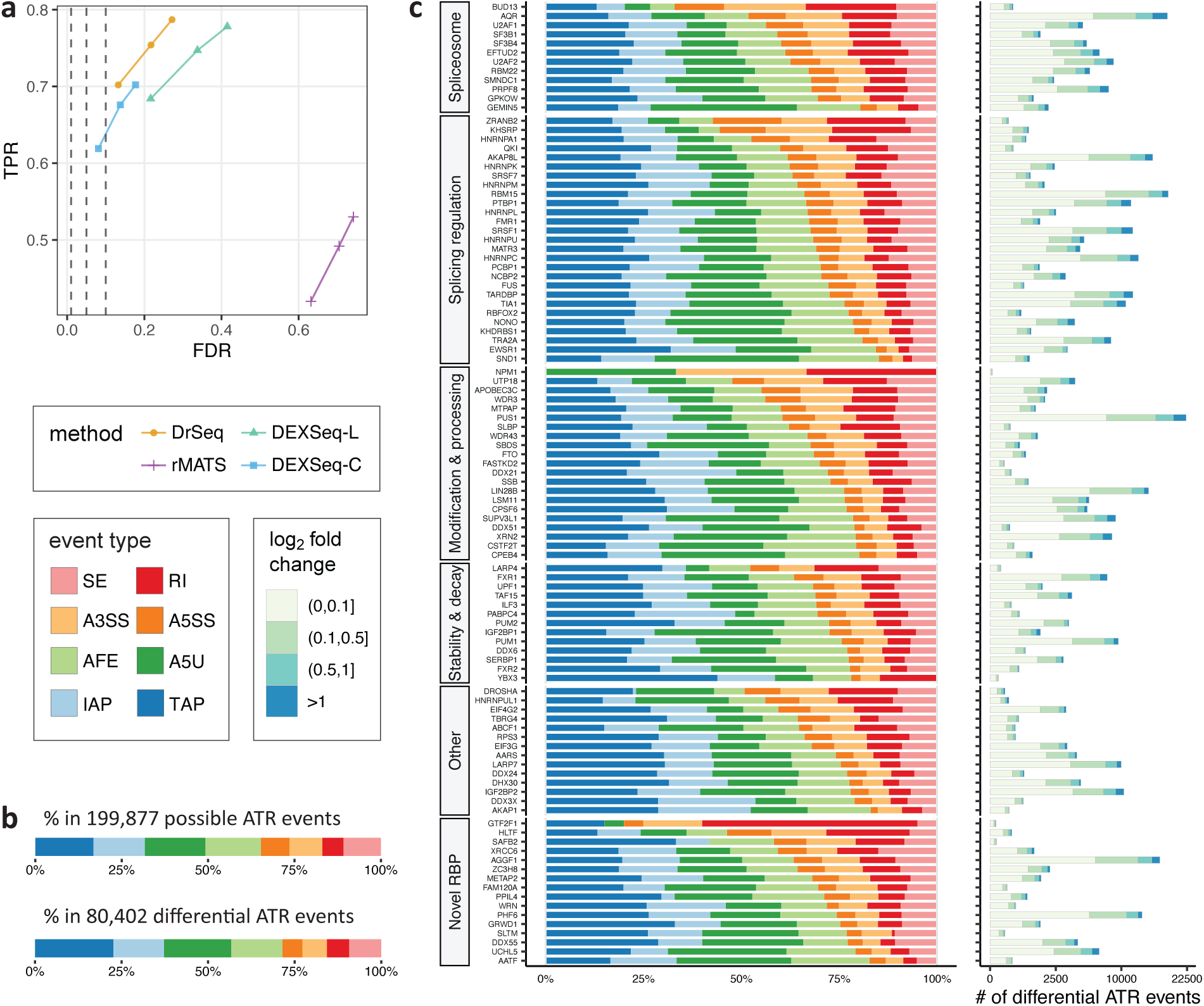
Detection of differential ATR events from RNA-seq data by DrSeq of SURF. (a) Comparison of the performances of DrSeq, rMATS, and two simple strategies for stitching DEXSeq inferences into ATR level (DEXSeq-L and DEXSeq-C), using simulated RNA-seq data. For each method, the three points display the true positive rate (TPR) and observed false discovery rate (FDR) at target FDR levels of 0.01, 0.05, and 0.1, respectively. (b) The distribution of eight event types in (upper panel) all 216,140 possible ATR events parsed from human genome annotation file (GENCODE version 24) and (lower panel) a total of 83,839 differential ATR events identified by DrSeq in at least one of the comparisons of 104 shRNA-seq experiments versus wild-type control (after duplication removal). (c) The distribution of differential ATR events identified by DrSeq in 104 RBP knockdown experiments across eight event types. The strip on the left-hand side of the plot groups RBPs by their previously reported primary functions [11]. For each RBP, the adjacent bar on the right-hand-side displays the total number of differential ATR events across all types.

Since a single ATR event typically spans multiple exonic bins, DrSeq combines data over exonic bins belonging to the same event and leads to better overall performance. In fact, the two “ad-hoc” approaches we designed either have satisfactory FDR control at the expense of power loss, or sustain a fairly large power while trading off FDR control. This analysis further supports that our extension of DEXSeq to ATR event level provides it with the competitive functionality as rMATS (i.e., event level resolution) in addition to its unique feature of encompassing differential analysis of ATI and APA event classes.

### 2.3 SURF reveals global positioning preferences of RBPs in AS, ATI, and APA events

We applied SURF to a collection of 104 paired RNA-seq and eCLIP-seq datasets from the v78 release of the ENCODE project. We extracted AS, ATI, and APA events from a filtered version of the current human genome annotation file (GENCODE [36] version 24) as described in Materials and Methods. The filtering procedure was motivated by the observation that isoform pre-filtering improves performance of count-based methods for analysis of differential transcript/exon usage [27] and resulted in 74,734 transcripts, 66,545 of which originated from protein coding genes. Collectively, these transcripts generated 199,877 AS, ATI, and APA events (upper panel in Fig. 3b). ATI and APA events constituted 65.31% of the total events with A5U at 17.50% and TAP at 16.86%, and SE, at 10.81%, was the most abundant category in AS.

Application of DrSeq to paired wild-type RNA-seq and shRNA-seq datasets of 104 RBPs in K562 cells identified 80,402 differential AS, ATI, and APA events (at FDR level of 0.05 and after duplication removal for analysis module 2). We observed that ATI and APA are the major sources of differential ATR (lower panel in Fig. 3b). In addition, we observed that a higher level of transcriptional complexity resulted from AS in the human genome than previously revealed by rMATS (supplementary Fig. S13) [11]. Overall, these results reflect both the direct (i.e., through occupancy of events by RBPs) and indirect effects of RBPs on transcriptome and instigate the previously reported findings for *D. Melanogaster* [37]. The observed complexity further accords with the results of a recent study on cross-tissue isoform differences [38]. Collectively, Fig. 3c shows the types and quantities of the differential ATR events. It highlights that (i) the impact of RBPs on ATR varies widely in magnitude (Fig. 3c), and (ii) many RBPs affect a wide variety of ATR events (directly or indirectly), indicating a potentially complex co-regulating network of RBPs (e.g., AQR and SF3B4 as depicted by the volcano plots in supplementary Figs. S14 and S15), and (iii) some RBPs exhibit event type specificity (e.g., CPSF6 exhibits evidence of TAP specificity, supplementary Fig. S16).

We next focused on 76 RBPs with eCLIP-seq data to study the direct effects of RBPs and their global positioning preferences for different types of ATR events. We limited our exposition to RBPs with at least 100 differential ATR events (i.e., increase or decrease upon knockdown) to enable association analysis. In addition, we estimated the average numbers of false positives in the association analysis by shuffling the grouping labels of differential ATR events and rerunning the analysis multiple times. This resulted in an average of 3.88 (with an s.d. of 2.38) false positive associations at target FDR level of 0.05 (supplementary Table S3), indicating a conservative overall error control. Fig. 4a reveals a large diversity in global positioning preferences for the 54 RBPs (see a full version of 76 RBPs in supplementary Fig. S17) consistent with the earlier work [39]. Specifically, direct protein-RNA interactions of 14 RBPs significantly associate with differential ATR in all of the eight ATR event types, while some RBPs exhibit a more specialized association, e.g., 24 RBPs govern only one or two ATR event types (Fig. 5a). At the individual RBP level, SURF recapitulated known binding preferences for a number of spliceosome components [40], while revealing additional general trends of RBP interactions with RNA (Fig. 4a). Focusing on SE events, for instance, we observed that AQR and PRPF8 binding at the flanking intronic regions of event body (location features *γ* and *ζ*) associated with differential SE significantly (at FDR level of 0.05, Fig. 4b and supplementary Figs. S18-S20). SF3B4 and SMNDC1 recognize the branching point (location feature *ε* of A3SS, supplementary Fig. S5) of A3SS (supplementary Figs. S21 and S22). In the class of ATI events, binding of GEMIN5 and AQR significantly associated with differential A5U (supplementary Figs. S18 and S23), whereas U2AF1 and U2AF2 exhibited association with differential AFE (supplementary Figs. S24 and S25). The latter further suggested involvement of spliceosomal components in recognizing alternative promoters [41]. In addition to these observations on spliceosomal components, SURF yielded various regulatory rules for non-spliceosomal RBPs. For example, we observed that CPSF6 [42] promotes inclusion of the distal termination sites in regulating TAP (supplementary Fig. S26), while UCHL5 promotes exclusion of the distal termination sites in TAP (supplementary Fig. S27).

**Fig. 4:**
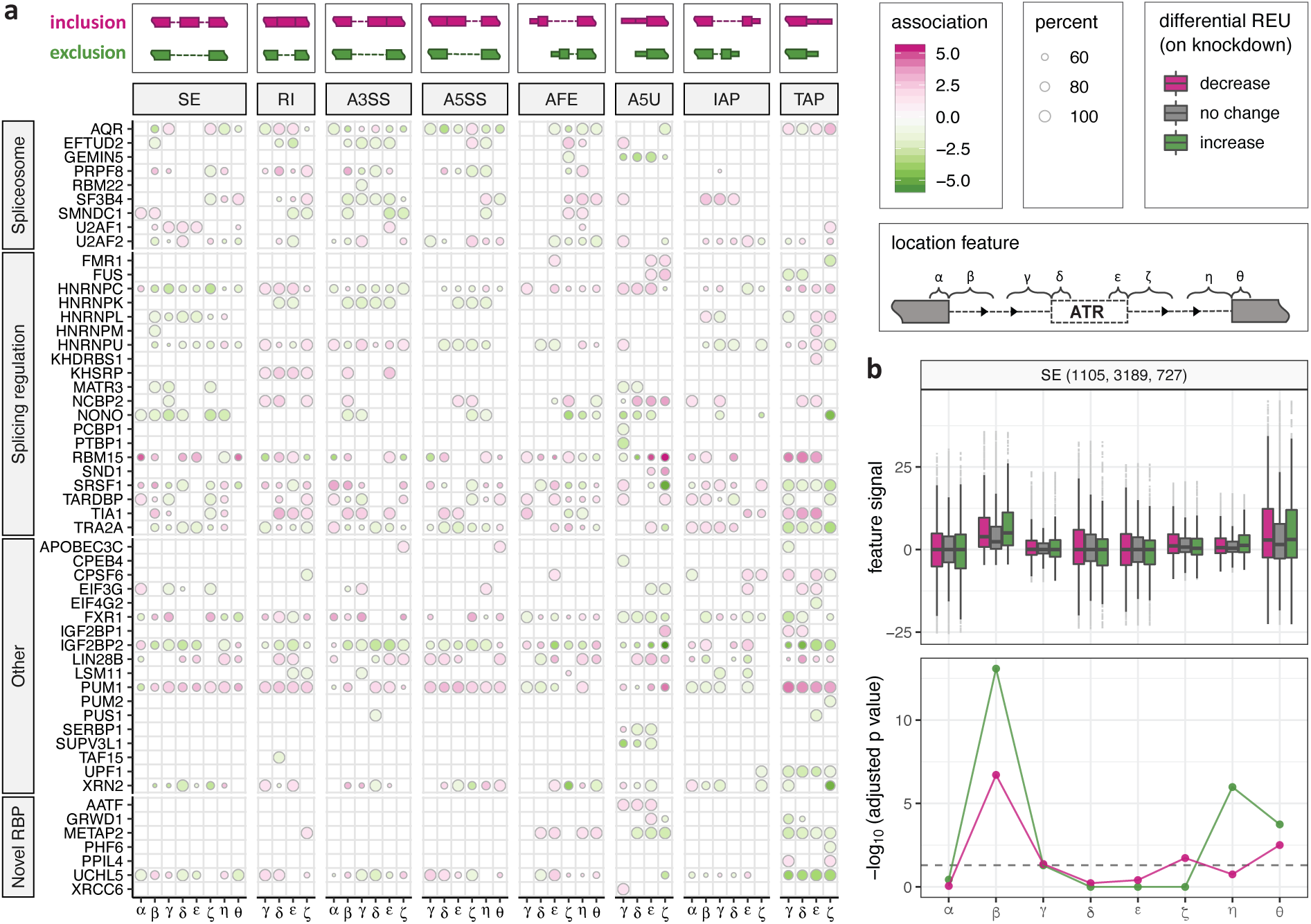
Global positioning associations with differential ATR. (a) SURF-identified associations (analysis module 2) of location features with differential ATR across 54 RBPs. Each row depicts a single RBP and each column represents one location feature, grouped by the corresponding ATR event types (column strips). Each circle symbol in individual cells indicates a significant association at FDR of 0.05. The color of the circles represents inclusion (pink) or exclusion (green) and the fill-in densities are determined by −log_10_ transformed adjusted p-values from the association testing. For features with dual functions (i.e., binding of RBPs at these features associate with both inclusion and exclusion), the circle size indicates the percentage of −log_10_ transformed adjusted p-value for the stronger one relative to the sum of both (i.e., smaller circles indicate similar associations). Illustrations above the column strips of event types depict the notions of exclusion and inclusion status of ATR. (b) Functional association plot of AQR for SE event. Upper panel box plot displays the distributions of feature signals among the three differential ATR groups (decrease, no change, increase). The numbers of SE events in each group are reported in parentheses at the top strip. The lower panel depicts the −log_10_ transformed p-values for each tested association after multiplicity correction. The dashed line indicates the FDR level of 0.05.

**Fig. 5:**
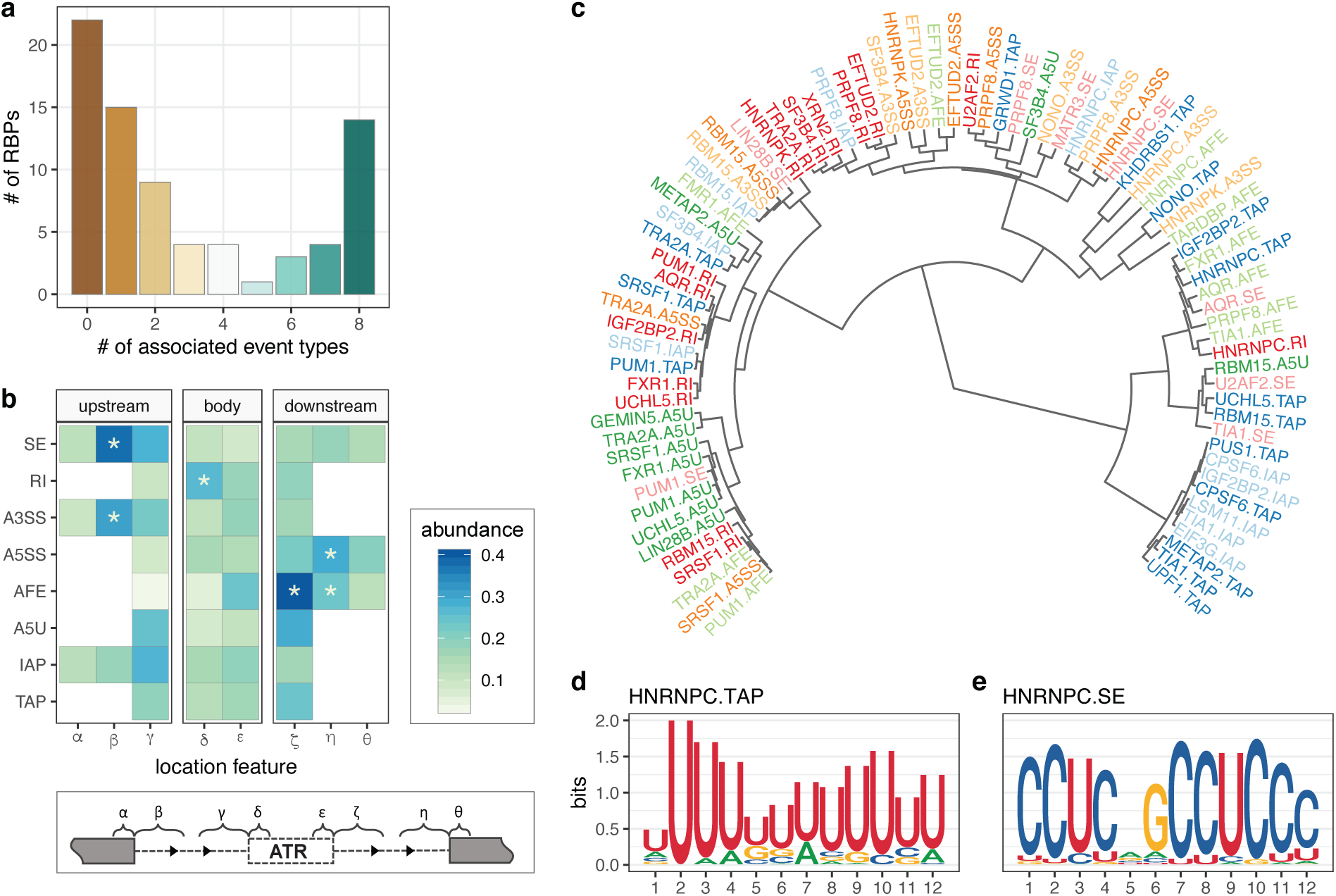
Summary of global positioning preferences across 76 RBPs and event-specific sequence preferences of RBPs. (a) The number of ATR event types that each RBP regulates, summarized from SURF (analysis module 2) results of 76 RBPs. (b) Aggregation of global positioning association results across each ATR event type. Column strips cluster location features into three groups: upstream, body, and downstream. The darker color indicates higher abundance of RBPs on the location feature (i.e., there are more RBPs for which the specific protein binding associates with differential ATR). Location features with a significant relative abundance at the significance level of 0.05 are marked with an asterisk. (c) Hierarchical clustering of motifs from *de novo* sequence analysis of location features for 82 RBP-event type combinations. Each tip point corresponds to the top motif found in SURF-identified location features that associated with a specific combination of RBP and ATR event. Going from the outer tips towards the root, the motifs that merge earlier have higher similarity. (d) Sequence motif identified in 394 SURF-inferred location features for HNRNPC in TAP (E-value 3.0 × 10^−68^). The total height of the letters depicts the information content of the position, in bits. (e) Sequence motif identified in 306 SURF-inferred location features for HNRNPC in SE (E-value 3.2 × 10^−152^).

While most global positioning tendencies of RBPs that associate with AS, ATI, and APA events (either previously reported [41, 43] or as revealed above) reside on or near the event body, our large-scale analysis with SURF instigates that many location features near the upstream (and sometimes downstream) constitutive exons of the ATR events also play fundamental roles in ATR through their interactions with RBPs. For example, SURF identified 21 RBPs as associating with SE by interacting at the intronic region adjacent to the upstream constitutive exon (location feature *β*). We further assessed the overall significance of each feature by aggregating the SURF-identified global positioning associations across all the 76 RBPs for each individual event type (Fig. 5b) as described in Materials and Methods. For SE events, the intronic region adjacent to upstream constitutive exon (*β*) exhibited marked enrichment compared to others with a p-value of 4.88 × 10^−4^ according to Pearson’s *χ*^2^ test. The same observation held for A3SS events (p-value of 1.79 × 10^−4^, Pearson’s *χ*^2^ test test) and highlighted the notable contribution from regions beyond proximal alternative splice sites. Overall, SURF identified 3′ (40.7%) RBPs that showed evidence for interaction with features distal to the event body, suggesting a broader genomic scope of functional protein-RNA interactions than previously revealed.

### 2.4 SURF-identified location features carry ATR-specific sequence motifs

The analysis module 2 of SURF identified 34,795 differential ATR events associated with binding of at least one of 54 RBPs. Aggregation across these associations resulted in 71,667 SURF-inferred location features (at FDR level of 0.05 and feature signal cut-off of 20, Materials and Methods), among which 34.9% (25,006) are associated with two or more RBPs. The numbers of inferred location features exhibited considerable heterogeneity across ATR types, with TAP having the largest numbers of inferred location features (19.2% and constituting 10.2% of all the possible location features of TAP event type), followed by A5U (15.5% and constituting 7.9% of all the possible location features of TAP event type) and SE (13.3% and constituting 5.5% of all the possible location features of A5U event type) classes (supplementary Fig. S28). supplementary Fig. S29 displays the distribution of genomic coverage by inferred features for all the eight event types.

Recent machine learning applications to predict pre-mRNA processing highlighted strong predictive power of sequence variants [44, 45, 46]. While the SURF framework does not emphasize *in silico* prediction of ATR events from sequence input, we expected some location features to harbor sequence motifs for RBPs. Leveraging the ability of SURF to identify distinct location features for different ATRs regulated by the same RBP, we asked whether these location features harbored distinct sequence motifs. Specifically, we performed *de novo* motif analysis with the MEME suite [47] for location features of 222 RBP-ATR event type combinations involving 52 distinct RBPs (Materials and Methods). This analysis captured a number of known, experimentally validated RBP motifs [48], e.g., TARDBP, U2AF2, HNRNPC, and TIA1. We next clustered these RBP-event type combinations, which involved 30 RBPs for which significant motifs were identified, based on the similarities of their identified motifs (E-values ≤ 0.01) (Fig. 5c). The event type-based clustering evident in Fig. 5c is partly driven by the shared location features of RBPs for the same event type (supplementary Fig. S30). However, the extent of overlap between the location features of different RBPs for the same event type is relatively low (supplementary Fig. S31) and indicates marked contribution of sequence similarities of the RBP-event type-specific motifs to the clustering structure. Overall, *de novo* sequence analysis of SURF-inferred features unveiled multiple unique sequence motifs of RBPs for distinct ATR events. Specifically, 19 RBPs exhibited motif multi-modality of varying degrees which we quantified by the maximum and minimum Kullback-Leibler divergence between all the learnt PWMs of individual RBPs (supplementary Fig. S32). For example, we observed that HNRNPC, a known poly(U)-binding regulator for the TAP class (E-value 3.0 × 10^−68^, Fig. 5d) [49, 50, 51], potentially recognizes a 5′-CCUC-3′ motif for the SE class (E-value 3.2 × 10^−152^, Fig. 5e). Furthermore, this clustering reconfirmed that while TIA1 overall prefers uridine-rich sequences [52, 53, 54], it also interacts with an AU-riched sequences [55], 5′-AAUAAA-3′, which resembles the canonical cleavage and polyadenylation site motif, in event types IAP and TAP in addition to two other significant motifs (supplementary Fig. S33).

### 2.5 SURF-identified location features are enriched for somatic mutations from ICGC and TCGA

Recent studies highlighted central roles of RBPs in neurodegeneration, auto-immune defects, and cancer [56, 57, 58, 59, 60, 61, 62, 63, 64]. We hypothesized that the 71,667 location features associated with differential ATR events should harbor more somatic mutations than expected by chance or expected in comparable regions of the genome not necessarily associated with such ATR events. To test this, we leveraged genome-wide mutation data from ICGC (excluding U.S. projects as they only contained coding mutations) and coding mutations from TCGA. Noting that the set of SURF-inferred features is a subset of all possible location features extracted from genome annotation, we designed the following control sets: (i) the full set of location features; (ii) union of span of events where the span is defined as the event body and its location features; (iii) genomic span of genes harboring all the events with an extension by 300bp upstream of transcription start sites and downstream of transcription end sites. These control sets form a nested set of regions from (i) to (iii) (Fig. 6a). Then, we calculated mutation per kilobase (MPK) as a measure of mutation quantification for each set of the genomic sets (Fig. 6b and supplementary Fig. S34). We observed an increasing enrichment of somatic mutations from genes to SURF-inferred features (p-value 3.00 × 10^−4^ from an exact Poisson test with a fold change of 1.50, supplementary Table S4). This enrichment highlights that location features inferred as associated with differential ATR events by SURF harbor markedly more somatic mutations compared to a large class of control sets. In addition, SURF-inferred location features also harbored more somatic mutations than the regions significantly bound in eCLIP-seq data (i.e., peaks with IDR *<* 0.01, covering a total length of 30.77 million bp and with an MPK of 17.08 compared to the 21.97 MPK of SURF-inferred location features, p-value 5.138 × 10^−5^ from an exact Poisson test with a fold change of 1.27). A similar result continued to hold with TCGA mutation data and is available from supplementary Fig. S35 and Table S5.

**Fig. 6:**
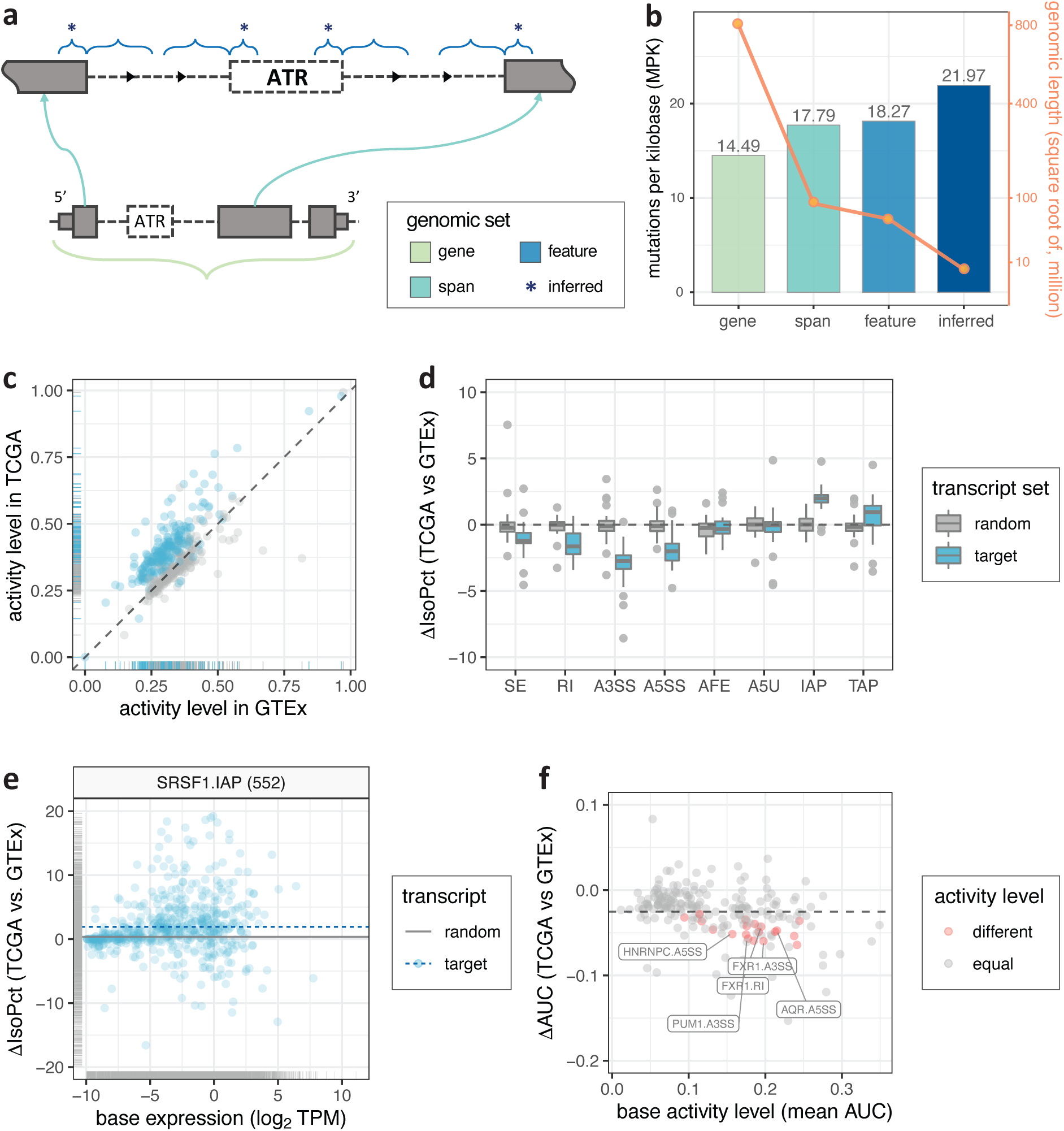
Integration of SURF results with large-scale datasets from ICGC, TCGA, and GTEx. (a) Illustration of RBP location features and three sets of control genomic regions. The set of SURF-inferred features (asterisks) is a subset of all possible location features (blue). The spanning regions (cyan) consist of event body and all possible location features. The set of spanning regions is a subset of the set of all genes (light green). (b) Mutations per kilobase (MPK) of SURF-inferred features and three control genomic sets. The bars depict the MPK quantified with somatic mutations in ICGC (non-US only) projects. The solid line with the right-hand side axis depicts the total length (in bp) of the genomic regions considered. (c) Comparison of gene activity level between normal whole blood and LAML tissues. Each dot depicts the mean AUC (activity level) of a gene set in whole blood (GTEx) and LAML (TCGA) samples. The 222 target gene sets are color in blue. The randomized control gene sets are colored in grey. (d) Changes in isoform percentages across eight ATR event types. Each data point corresponds to either a target transcript set (blue) or a randomized control transcript set (grey). The mean difference in isoform percentages (∆IsoPct) from normal whole blood (GTEx) to LAML (TCGA) samples is averaged across individual transcript sets. Each of the 222 transcript sets is depicted in one of eight associated ATR event types. (e) Change in isoform percentages (∆IsoPct) of 552 SURF-inferred transcript targets of SRSF1 in IAP regulation. A blue dashed line indicates the mean of ∆IsoPct across the full set. The grey solid line indicates the mean of ∆IsoPct across all transcripts in a randomized control set. (f) Detection of differential activity by discover module of SURF for 222 target transcript sets. Each dot depicts, for one RBP-ATR event type combination, the base activity level (x-axis) and the observed change in the activity level (∆AUC) from normal (TCGA) to tumor (GTEx) samples. Transcript sets with significant differential activity (FDR *<* 0.05) are marked in red. The grey dashed line indicates the mean of ∆AUC across all control transcript sets.

### 2.6 SURF-identified ATR targets of RBPs exhibit differential transcriptional activity between TCGA acute myeloid leukemia tumor and GTEx whole blood samples

We next performed a comparative analysis of transcriptional activity of genes and transcripts harboring specific RBP-ATR event type combinations in normal versus tumor samples by leveraging GTEx and TCGA resources. We specifically considered TCGA LAML sample for which the K562 erythroleukemic cell line has the most relevance for. Harnessing of gene and transcript targets based on SURF-inferred location features capitalizes our construction of location features – each location feature is designated to a single ATR event that originates from a unique transcript. Among 222 RBP-ATR event type combinations, 132 (or 131) of them had more than 100 genes (or transcripts) with at least one SURF-inferred location feature (supplementary Fig. S36). On average, protein-coding genes compromised 97.3% of all the gene sets. We first asked whether the activity of an RBP, as measured by expression of its target genes, significantly varied between 173 GTEx whole blood samples and 337 TCGA LAML samples (supplementary Fig. S37). To account for potential within cohort biases of the RNA-seq data, we constructed randomized control sets for each gene set by matching their sizes and average activity levels (as measured by AUC, Materials and Methods). With such control, we observed a general shift of target gene activity from normal to tumor tissues (Fig. 6c and supplementary Fig. S38), suggesting potential importance of SURF-identified RBP targets for LAML.

We next queried whether specific transcript targets of RBPs behaved differentially among all isoforms originating from the same gene, across tissue conditions. To this end, for every transcript, we quantified the changes in isoform percentage (`IsoPct` as reported by RSEM) from GTEx whole blood samples to TCGA LAML samples. Isoform percentage of a transcript depicts the percentage of its abundance over its parent gene’s abundance and its comparison across conditions elucidates how the contribution of a transcript to overall gene abundance is changing across conditions. Comparison of isoform percentages revealed a diverse pattern across ATR event types (Fig. 6d), suggesting a refined transcript level biological relevance of SURF-identified RBP targets. Specifically, transcript sets associated with AS events (AS, RI, A3SS, and A5SS) exhibited reduced isoform percentage (as compared to matched random control sets from other isoforms originating from the same gene) between normal (GTEx) and tumor (TCGA) tissues (supplementary Fig. S39). In contrast, transcript sets associated with APA events (namely IAP and TAP) exhibited increased isoform percentage in tumor samples compared to normal samples (e.g., SRSF1 in IAP, Fig. 6e), indicating that transcripts harboring these event types contributed more to their parent gene’s overall abundance. This observation is further supported by multiple replicates of random control and after de-duplicating transcripts across event types (supplementary Figs. S40 and S41). Since ATR events are defined by variable bins, which originate from shorter transcripts (i.e., those with absent exonic parts), this observation suggests that the transcripts that terminate earlier at 3′ end, as well as those with more internal “spliced-in” exonic parts, tend to be more transcriptionally active in the LAML cancer tissue compared to normal blood tissue. Shortening of 3′ UTRs is widely observed during tumorigenesis and cancer progression [65, 66]. In addition to supporting this widely observed phenomenon, our results suggest that AS and APA events contribute more to transcriptome diversity in cancer compared to ATI events. To test the differential activity for individual transcript sets, we applied the discovery module of SURF, with their paired controls by including transcripts originating from the same corresponding target genes. Of all the RBP-ATR event type combinations, 18 of them (out of 131 with more than 100 transcripts) exhibited significantly different transcriptional activity between the whole blood (GTEx) samples and LAML (TCGA) samples, at FDR cut-off of 0.05 (Fig. 6f). The detected transcript sets originate from 9 RBPs, among which FXR1, SRSF1, and U2AF2 associate with the most diverse ATR event types (SE, A3SS, A5SS for all three RBPs, and RI for FXR1 additionally), followed by AQR and HNRNPC through two event types (supplementary Table S6), many of which have previously reported implications in cancer (e.g., [67, 68]). All but two of these RBPs (FXR1 and PUM1) are splicing factor genes and U2AF1 exhibited significant driver mutation patterns across a wide variety of cancer types [63].

## 3 Discussion and Conclusions

In this study, we presented SURF, a new integrative framework for the analysis of large-scale CLIP-seq and coupled RNA-seq data and applied it to ENCODE consortium data (Table 1). A key advancement in SURF is the extension of differential exon usage method DEXSeq to differential alternative transcriptional regulation at the event level. We presented a formulation, DrSeq, that avoided *ad hoc* downstream analysis stitching DEXSeq results into event level, while controlling FDR and achieving as well or better power than alternatives. In addition, DrSeq uniquely enabled differential ATR detection for alternative transcription initiation and alternative polyadenylation events, a feature currently lacking in the widely adapted tool rMATS. SURF performs all integrated analyses based on the AS, ATI, and APA events as experimental units. Event-centric analyses are preferred when focusing on alternative splicing at the level of individual splicing events, e.g., the inclusion of a skipped exon, or the usage of a particular alternative splicing site. Interpreting the results of event-centric analyses is straightforward, and follow up validation experiments for differential AS, ATI, and APA events are easier to design.

**Table 1:** Overview of the data used by individual modules of SURF. *: RBPs lacking at least 100 differential ATR events are excluded from the analysis module 2. **: The discovery module studies a subset of RBPs in * with at least one inferred location feature.

| Module | # of RBPs | Input datasets (sources) |
| --- | --- | --- |
| 1 – DrSeq | 104 | 104 shRNA-seq with paired control RNA-seq from wild-type condition (ENCODE) |
| 2 – Association | 73* | 73 eCLIP-seq with paired control datasets from size-matched input library (ENCODE) |
| 3 – Discovery | 54** | 173 whole blood RNA-seq (GTEx) and 337 LAML RNA-seq (TCGA) |

Association of DrSeq-identified differential ATR results to global positioning preferences of RBPs with eCLIP-seq data revealed large diversity among RBPs and suggested dual inclusion and exclusion roles for RBPs based on the positioning of protein-RNA interactions. Furthermore, aggregation of the positioning preferences across RBPs within each event type revealed that locations near the upstream and sometimes downstream constitutive exons of the ATR events also play a significant role in ATR. Sequence analysis of SURF-inferred location features (e.g., regions that SURF deemed as significantly related to ATR) revealed known and novel interaction modalities for RBPs in ATR. To further assess the overall relevance and importance of SURF-inferred location features, we compared the somatic mutation densities of these regions with carefully designed controls. This analysis revealed that location features inferred as associated with differential ATR events harbored significantly more somatic mutations both in ICGC and TCGA consortia data. Further integration with reference transcriptomes from GTEx and TCGA highlighted contribution of APA and AS regulation in diverse transcriptome activity in cancer and revealed RBPs with ATR event specific differential transcriptional activities in LAML tumors compared to GTEx normals.

SURF presents the first framework for systematic integrative analysis of large collections of CLIP-seq and RNA-seq data and includes a discovery module for probing public TCGA and GTEx transcriptome data. This comprehensive approach coupled with the ability to identify differential ATR events from settings with multiple biological replicates underlines SURF’s potential as an integrative tool. There are a number of directions that naturally build on the results of SURF. Since we have observed significant enrichment of somatic mutations from ICGC and TCGA in SURF-identified location features, evaluation of impact of individual somatic mutations on RBP binding with *in silico* calculations [69, 70] could elucidate how such mutations might be manifesting their effects. SURF focused on each RBP marginally. Given the complex combinatorial nature of ATR [43, 11], a promising extension to SURF framework should be in the direction of multivariate analysis of large collections of RBPs.

## 4 Materials and Methods

### 4.1 Isoform pre-filtering

In order to improve FDR control, we pre-filtered transcripts in the human genome annotation file (GENCODE version 24) before extracting ATR events, as suggested by [27]. We performed the filtering based on the RSEM [71]) transcriptome quantification of 486 RNA-seq datasets, including 220 RBP knockdown experiments (with two replicates) and 23 wild-type control experiments (with two replicates). Specifically, the transcripts with (i) relative isoform abundance estimate (`IsoPct`, IP) of less than *c*% across all samples or (ii) TPM estimate less than 1 across all samples were removed. We varied the threshold *c* for filtering relative isoform abundance as 5%, 10%, and 25% and considered the following strategies: (1) IP ≥ 5%, (2) IP ≥ 10%, (3) IP ≥ 25%, (4) TPM ≥ 1, and (5) IP ≥ 5% and TPM ≥ 1. supplementary Table S7 summarizes the filtering thresholds, the number of retained genes, transcripts, and the sum of TPM (averaged across all samples). The total read counts aggregated across transcripts behaved similar to the sum of TPM, thus is omitted from the table. This investigation suggested that isoform pre-filtering markedly reduces the transcriptome complexity of the raw genome annotation, while possessing the vast majority of transcriptome signals. In fact, 70% of the genes in the filtered set were protein-coding as opposed to 32% in the original file. This implies that majority of the filtered transcripts are from non-coding genes.

### 4.2 Parsing AS, ATI, and APA events from genome annotation

The parsing procedure consists of two steps. First, we define *variable* bins of exonic regions for each transcript as the set of absent exonic regions of the transcript compared to the full gene model (supplementary Fig. S3). This definition is previously introduced by [33], and it enjoys the property that all variable bins of a transcript are mutually disjoint. We group runs of variable bins (i.e., merge any consecutive ones within the transcript) and categorize each as an ATR *event*. In the second step, we invoke a decision tree to uniquely annotate each event to a specific category among alternative splicing (AS) events including exon skipping (SE), alternative 3′ (A3SS) or 5′ (A5SS) splicing, intron retention (RI), alternative first exon (AFE), alternative 5′UTR (A5U), and intronic (IAP) or tandem (TAP) alternative polyadenylation, according to its relative position within the gene (supplementary Fig. S4). Note that all the ATR events from one transcript are isolated from each other, which essentially forges the uniqueness of event annotations. We interchangeably refer to the event as an *event body*. Finally, we deduplicated the events from different transcripts of the same gene that (i) shared the same genomic information (coordinates, strand, and constitutive upstream/downstream exons) and (ii) were classified into the same event type.

### 4.3 Extracting location-specific features

We generated a set of location features relevant to the ATR events according to their event types (supplementary Fig. S5). These location features were designed to capture RBP binding signals in the upstream/downstream constitutive exons of the event, starting and end sites of the event body, as well as adjacent intronic regions. For example, an SE event has eight location features: two 100bp regions from the upstream and downstream constitutive exons, four 300bp regions from two flanking introns (upstream and downstream) of the event body (i.e., the cassette exon), and two 100bp flanking regions spanning the start and end of the event body (Fig. 2c). For the flanking or spanning regions that were shorter than the desired length of 100bp or 300bp, we used one half of the exonic or intronic length. The number of location features for the other ATR event types ranges between 4 to 8. Each location feature represents a global positioning preference that potentially associates with the DrSeq-inferred differential ATR status. Detailed pictorial illustrations of location features for each event type are depicted in supplementary Fig. S5.

### 4.4 DrSeq for detecting differential alternative transcriptional regulation

Differential ATR event detection is performed in two phases: RNA-seq read counting and statistical inference. The entire procedure takes as input the annotation of the ATR events and aligned reads from RNA-seq samples and outputs for each ATR event the result of differential testing including the test statistics, p-values, estimated log2 fold changes, and additional summary objects. For each event, DrSeq counts RNA-seq reads at two levels, event body and the gene to which the event belongs, by resorting to `featureCounts` [72]. While this counting scheme allows reads to overlap multiple events or genes, diagnostic plots did not reveal any adverse effects (supplementary Fig. S42). Since DrSeq tests for each event separately, each read is used at most once in each test. DrSeq invokes `DEXSeq` as an inference engine to fit the differential ATR model (1). Specifically, the events are grouped by their types and model fitting is performed separately. This allows capturing the mean-dispersion relationships in an event type specific manner and reduces dependency among the tests (supplementary Fig. S43). After the model fitting and testing, all p-values are jointly corrected for multiplicity with the BH procedure.

### 4.5 Simulation design for differential ATR detection

We simulated RNA-seq data following the strategy by [27]. First, RSEM model parameters were estimated from two real RNA-seq assays (ArrayExpress dataset E-MTAB-3766 in [73]). Then, a mean/dispersion relationship, learnt from real data [74], was used to generate sample wise transcript expression estimates, for three replicates of the wild-type and three replicates of differential conditions. To create the differential condition, we sampled 1,000 genes and introduced differential transcript usage by switching the relative abundances for the two most expressed transcripts. These genes were selected randomly among those with sufficiently high expression (expected count greater than 500) and at least two sufficiently highly expressed isoforms (i.e., with relative expression abundance, as calculated by RSEM, above 10%). Since the isoforms have varying lengths, the expected read count for these genes varied between the conditions, while the TPM remained similar. This procedure generated a spectrum of differential ATR events and eliminated potential confounding due to gene-level differential expression. Finally, we used RSEM to simulate paired fastq files with a read length of 101bp for each sample. For DrSeq, we used the default pipeline. For DEXSeq, we used default settings with the exception of invoking the -r no configuration to avoid merging overlapping genes into complexes. For rMATS (version 4.0.2), we used the default cut-off of 0.0001 for splicing difference and employed the same FDR thresholds as DrSeq. For MAJIQ (version 2.1), we administered both the default setting and a liberal setting where the change threshold of MAJIQ (`--threshold`) is set to 0.1.

### 4.6 Read coverage of ATR events from RNA-seq

In the analysis module 2 of SURF, we filtered out (i) low coverage events and (ii) overlapping ATR events of the same type to avoid near-duplication observations in the association analysis. Specifically, we employed a *read coverage* measure that is akin to the Fragments Per Kilobase Million (FPKM) in transcriptome quantification for RNA-seq. For a given event, we first counted the number of reads residing on the event body (in million), where we utilized only the reads with mapping quality score greater than 10 (this threshold also applies to eCLIP-seq reads). To account for sequencing depth differences, the counts were then normalized by the size factor of each sample [75] and averaged across all four samples (two replicates per condition). We refer the resulting quantity as the *base mean*, which is also reported by DrSeq. Next, we adjusted the base mean by dividing it with the pseudo length of each event (in kilobase) and used this adjusted mean as the coverage of ATR events. Here, the *pseudo length* of an ATR event is defined as its actual length (in kilobase) shifted by the read length minus one, that is, 99 bp (all shRNA-seq experiments from the ENCODE consortium have a read length of 100 bp). The adjustment with pseudo length (as opposed to the actual length) regularized those extremely short (in length) ATR events and reduced the effects of overlapping counting. This had negligible effect on long ATR events. Finally, we excluded ATR events with read coverage smaller than 0.05, and among the overlapping ATR events, we only kept the event with the highest read coverage for the downstream analysis.

### 4.7 Generation of feature signals from eCLIP-seq data

Each ATR event has a set of location features, each of which is a genomic region of potential protein-RNA interaction. This step of SURF takes as input the aligned reads from the IP and accompanying control input samples of eCLIP-seq and summarizes the eCLIP-seq signals at these features. To this end, for each feature, it first counts the number of overlapping reads (in million) in each sample by requiring an overlap of at least 12 bp between the read and the location feature. The 12 bp corresponds to the 25% of the read length in the ENCODE eCLIP-seq experiments. Then, the read counts are normalized by the sequencing depth of each sample. Next, the normalized read counts are weighted by the genomic length (in kilobase) of the feature, which is then averaged separately across the IP and control input samples. Finally, the difference between normalized IP and input samples is reported as *feature signal*.

### 4.8 Association analysis for global positioning preferences

We used a logistic regression framework to derive global positioning preferences, e.g., position specific rules for RBPs in regulating ATR. Specifically, for each event *i* with event type τ (*i*) = *t*, let *x*_*ik*_ denote the feature signal at the *k* -th location feature. The random variable *Y*_*i*_ ∼ Bernoulli(*p*_*t*_) corresponds to whether an event is differentially regulated (*Y*_*i*_ = 1) or not (*Y*_*i*_ = 0). We link its mean parameter *p*_*t*_ to the eCLIP-signals of the location features marginally with a logit function as

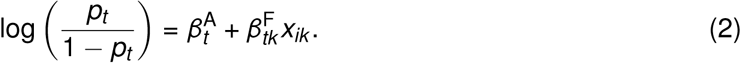

Here, parameter 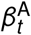 represents the grand effect of ATR event *t* and quantifies a baseline for the probability of observing a differential ATR event. This parameter also accounts for the position independent effects of RBP in regulating this event type. Position-specific main effect 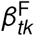 captures the global positioning effect of RBP at the specific location feature *k*.

In fitting the above model for every ATR event type *t*, we also considered increased- and decreased-regulation separately: model for *Y*_*i*_ = 1 corresponds to increased-regulation (p-value *<* 0.05 and log_2_ fold change *>* 0) and model for *Y*_*i*_ = 1 represents decreased-regulation (p-value *<* 0.05 and log_2_ fold change *<* 0). Both models incorporated the same set of equal-regulation (i.e., “no change”) events for *Y*_*i*_ = 0 (p-value *>* 0.4). The reason for separating increased- and decreased-regulation is primarily due to the possibility that interactions of a single RBP at a specific pre-mRNA location feature might result in both inclusion and exclusion of the alternative site, depending on the presence/absent of other collaborative/competitive RBPs. Differential ATR analysis of the ENCODE compendium data of 104 RBPs with module 1 of SURF resulted in, on average and after filtering out overlaps, 2,182.02 (1.01% of all tested events) events with increased REU, 1,914.98 (0.89% of all tested events) events with decreased REU. This enabled fitting of model (2) and testing of the position-specific main effect parameter 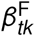 by a standard likelihood ratio test. Tests for all the location features were corrected for multiplicity with the BH procedure at FDR level of 0.05.

### 4.9 Summarizing association tests across RBPs

We quantified the abundance of a location feature by the normalized number of RBPs that significantly associated with it. Specifically, we aggregated the results of global positioning associations across all the 76 RBPs for each individual event type. For each RBP and each event type, we first applied minus logarithm (base 10) transformation on p-values and normalized them to have unit sum. The normalization accounts for the fact that an RBP may interact with its target RNA (pre-mRNA) at multiple location features. Then, for every event type, these values were averaged across RBPs and interpreted as *feature abundance* (Fig. 5c). Next, we evaluated whether any of the features were significantly more abundant in comparison to other features of the same ATR event type by a Pearson’s *χ*^2^ test on two variables denoting (i) whether or not a location feature is significantly associated with the differential ATR and (ii) whether or not the location feature is *x*, where *x* ∈ {*α*, *β*, …, *θ*} are the set of location features for that ATR event type. All the tests employed Yates’s correction for continuity.

### 4.10 Extracting SURF-inferred location features

As a result of association testing in the analysis module 2 of SURF, each location feature is assigned a p-value (adjusted for multiplicity when appropriate). SURF then generates a list of *inferred* location features by filtering as follows. First, it selects statistically significant location features at FDR of 0.05. Note that each selected feature has a corresponding training dataset, which consists of differential ATR events and control events. SURF then extracts the list of genomic regions for each selected location feature from this corresponding set of differential ATR events. Finally, leveraging the quantified feature signals from the analysis module 2, it filters out the genomic regions with feature signals, i.e., normalized background adjusted eCLIP-seq read counts, below 20 and defines the set of remaining genomic regions as the inferred features. Imposing the eCLIP-seq signal condition ensures that the set reflects the consequences of *direct* protein-RNA interaction. Note that, the inferred features are event-specific and are associated with specific regulation status. As a result, each inferred feature has its associated ATR event and inferred RBP regulation function (i.e., “inclusion” or “exclusion”).

### 4.11 Sequence analysis for SURF-inferred location features

For SURF-inferred features, we performed *de novo* motif discovery using the MEME-suite [76], with a zero or one occurrence per sequence (ZOOPS) model in the default mode with background distribution estimated from the input sequences. Specifically, we extracted the genome sequence (human genome version GRCh38) of each SURF-inferred location features, which were then assembled into 326 various collections based on their associated ATR event type, RBP, and feature type (e.g. *α*, *β*,…). The collections of interest corresponded to (i) 52 individual RBPs, (ii) 8 types of ATR events, (iii) 44 event × location feature combinations (e.g., SE-*α*, SE-*β*, etc), and (iv) 222 RBP-ATR event type combinations (e.g., SRSF1-SE). Finally, we searched for the top-3 motifs with the minimum width of 5bp and the maximum of 12bp in each collection of sequences, where we filtered the resulting motifs by a minimum occurrences of 40 times. The SURF-inferred location features and the output of 326 individual MEME runs (version 5.0.5) are deposited (see Availability of supporting data). For clustering of identified motifs, we used a metric based on Kullback-Leibler divergence of position weight matrices (PWMs) as adapted in [77] and the Wald’s agglomeration method [78].

### 4.12 Differential activity of transcript sets

The discovery module of SURF takes in sets of RBP targets (e.g., transcripts or genes) that harbor SURF-inferred location features and perform differential analysis on these sets across conditions, in terms of their activity. This module adapts a rank-based approach, inspired by the single-cell RNA-seq application of [35], to score the *activity* of a target set in a sample by the area under the recovery curve (AUC). The AUC evaluates the ranking of its elements compared to other quantified units in the sample in terms of expression (supplementary Fig. S6). As a result, it transforms the transcriptome expression across samples into a measure of activity across samples and enables testing of differential activity across conditions. To calibrate the activity scores within each condition, this module also allows a paired control set for each target set. To enable subsampling for generating a null distribution, the control set is preferably larger in size than the corresponding target set. The differential analysis then tests for the difference in AUC between the two conditions using the difference in AUC. Our test statistic is the difference in AUC between the two conditions (e.g., GTEx vs. TCGA). An empirical null distribution for this statistic is estimated by randomly subsampling from the control set 5, 000 times. Each subsampling matches the size of the target set. The p-values are adjusted to control for FDR using the BH procedure (see supplementary Fig. S44 for the histogram of raw p-values realized with 222 target transcript sets).

## Declarations

### Supplementary figures and tables

See “Supporting Information” for supplementary Figs. S1-S44 and Tables S1-S7.

### Availability of supporting data

The datasets supporting the conclusions of this article are available in the following repositories. The human genome annotation file (version 24) is available from the GENCODE project [36], https://www.gencodegenes.org. The shRNA-seq and the eCLIP-seq datasets were accessed from the ENCODE project [9], https://www.encodeproject.org, lastly on October 28, 2018. The somatic mutation datasets for both the TCGA and ICGC projects and the transcript expression datasets (RSEM quantification) for both the TCGA and the GTEx projects were downloaded from the UCSC Xena platform [79], http://xena.ucsc.edu. The processed data and the results of SURF analysis are deposited at Zenodo, https://doi.org/10.5281/zenodo.3779037 [80]. The source code for reproducing the results of this paper is released at GitHub, https://github.com/keleslab/surf-paper [81].

## Acknowledgements

We are grateful to Michael Cammilleri for his endless technical support in high throughput computing. We thank Marvin Wickens, Judith Kimble, Heidi Dvinge, Brian Carrick, and members of Keleş Group for many helpful discussions. We also thank the anonymous reviewers for their detailed comments that led to improvements in the manuscript.

