## Supporting Information for "SURF: integrative analysis of a compendium of RNA-seq and CLIP-seq datasets highlights complex governing of alternative transcriptional regulation by RNA-binding proteins"

The document provides supporting information to "SURF: integrative analysis of a compendium of RNA-seq and CLIP-seq datasets highlights complex governing of alternative transcriptional regulation by RNA-binding proteins." This contains supplementary Figs. S1-S44 and Tables S1-S7.

#### Supplementary Figures

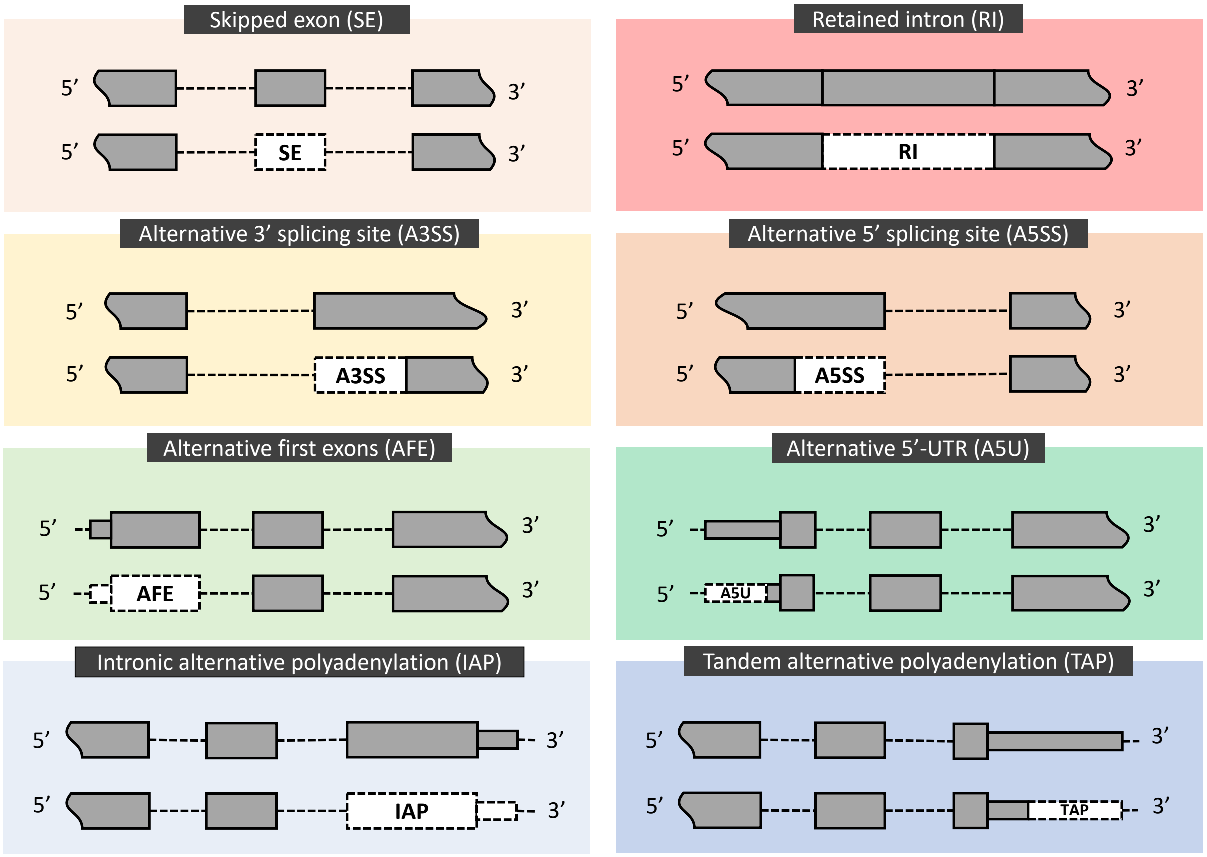


Supplementary Fig. S1: Illustration of eight ATR event types: exon skipping (SE), alternative 3′ (A3SS) and 5′ (A5SS) splicing, and intron retention (RI) within the AS class; alternative first exon (AFE) and alternative 5′UTR (A5U) within the ATI class; intronic (IAP) and tandem (TAP) alternative polyadenylation within the APA class. In each panel, the upper track depicts part of a gene model, and the lower track demarcates a specific ATR event in a transcript (isoform) with a white dashed box.

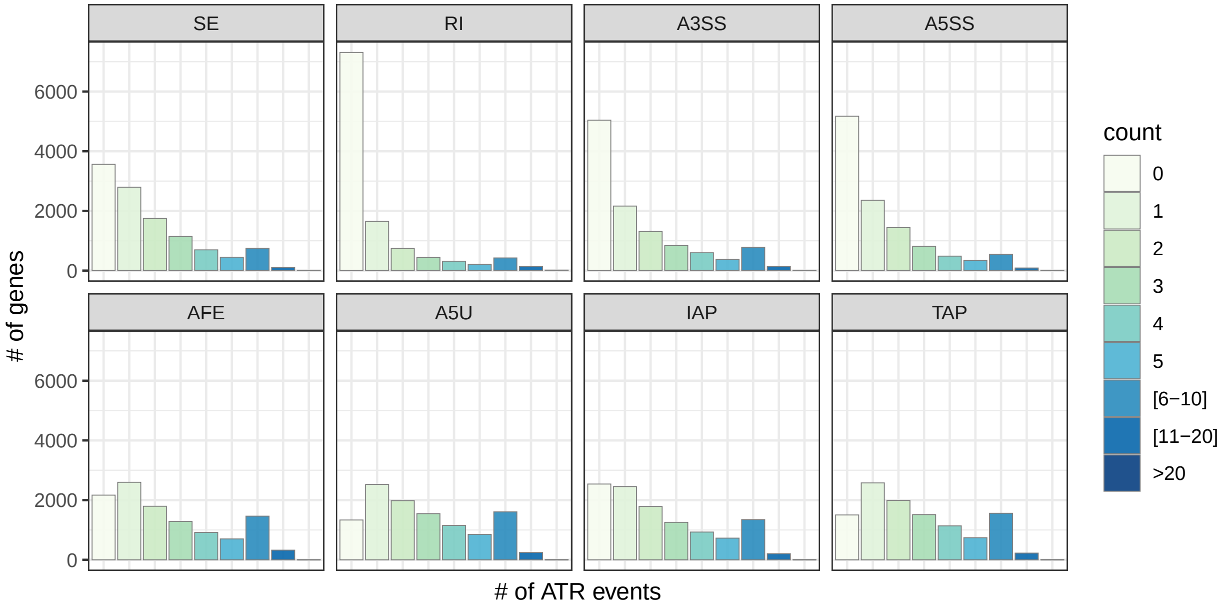


Supplementary Fig. S2: Distribution of numbers of ATR events parsed out from human genome annotation (GENCODE version 24), stratified by the event type.

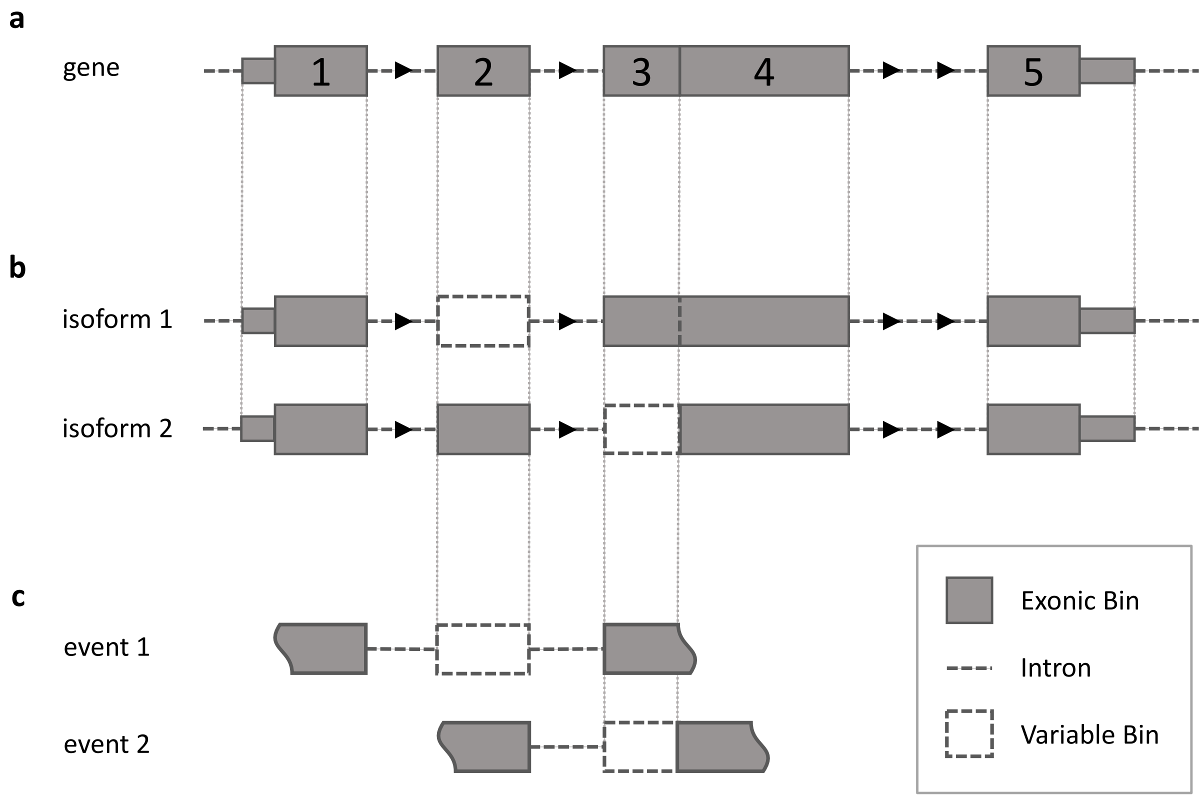


Supplementary Fig. S3: Identification of ATR events from genome annotation. (a) All exons of the same gene are discretized into mutually disjoint exonic bins (numbered grey boxes). (b) Then, for every transcript (isoform) from the same gene, absent exonic bins are labeled as variable bins (white dashed boxes, e.g., exonic bin # 2 is a variable bin for isoform 1 and exonic bin # 3 is a variable bin for isoform 3). (c) Finally, within each transcript, consecutive variable bins are merged together and are labeled as an ATR event. A transcript can harbor multiple ATR events.

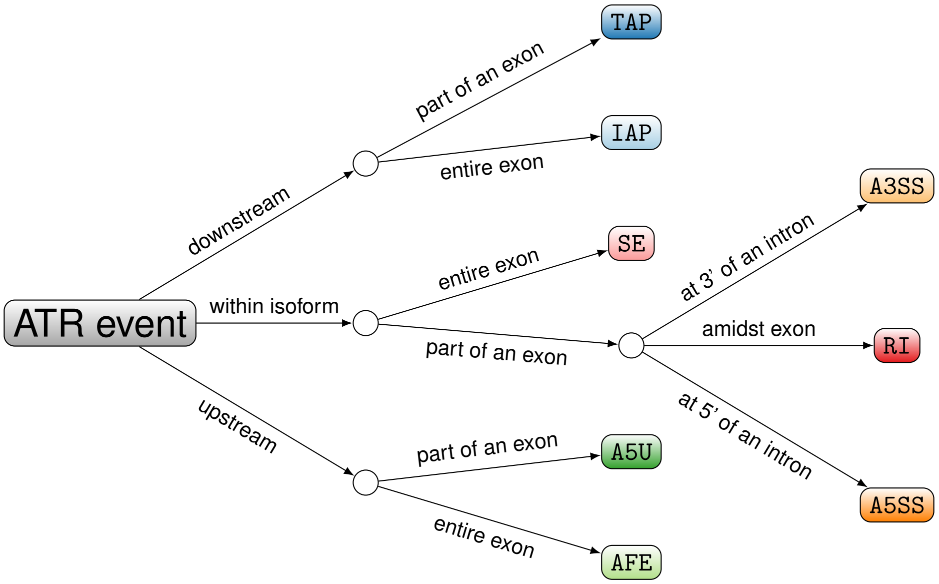


Supplementary Fig. S4: Decision tree for annotating variable bins as ATR events. The decision tree takes a run of variable bins as input and classifies it into one of the eight event types. “downstream” and “upstream” refer to whether the variable bin is at the 3′ or 5′ end, respectively.

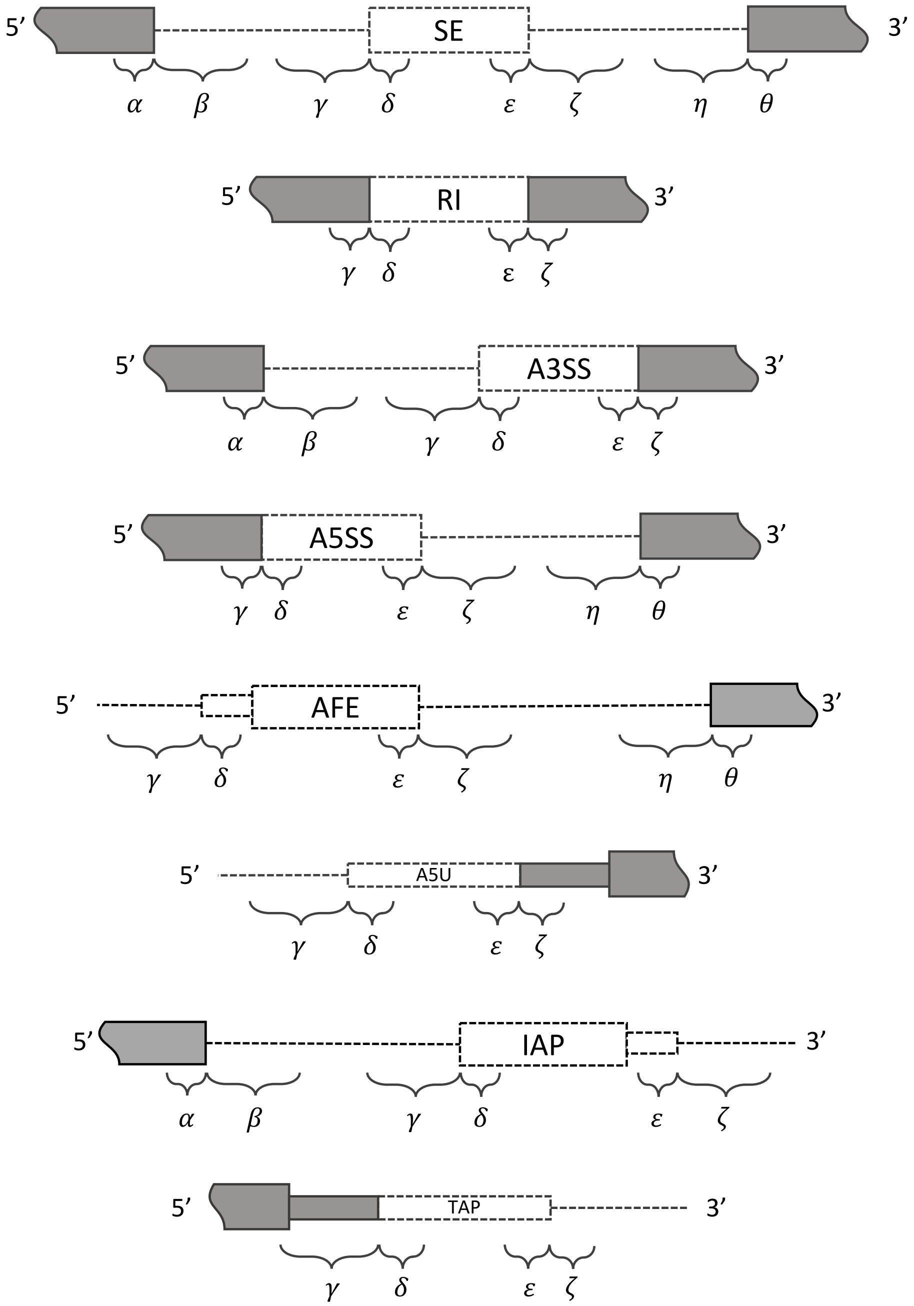


Supplementary Fig. S5: Illustration of location features for eight ATR event types. White boxes depict the ATR events with the event type labeled inside. Short and long curly brackets correspond to genomic regions of length 100bp and 300bp respectively.

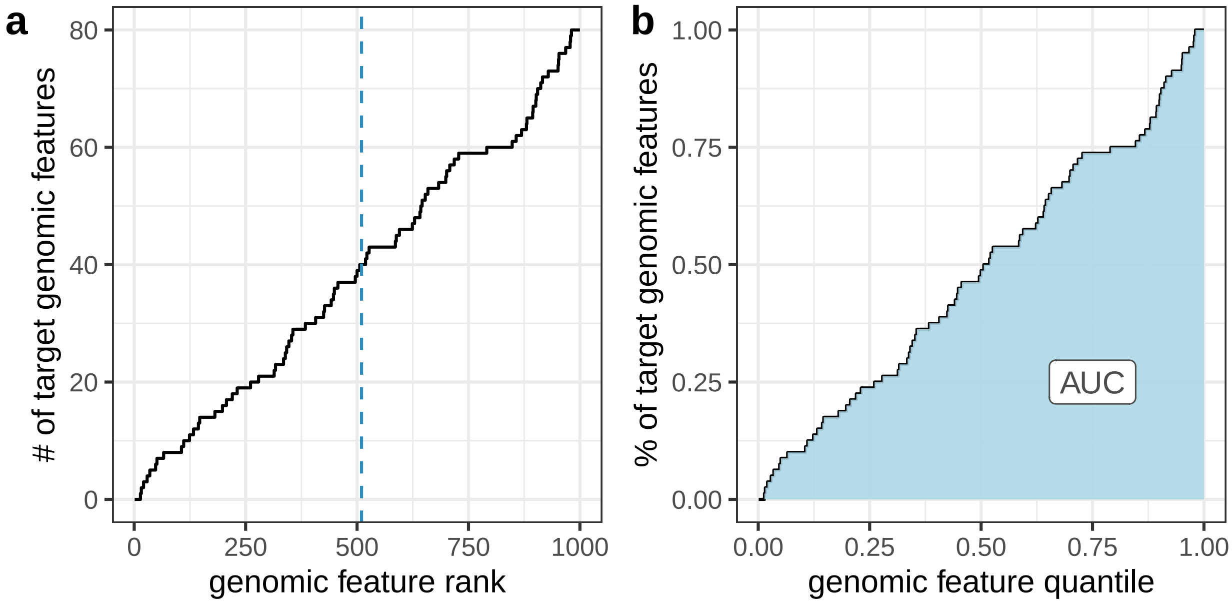


Supplementary Fig. S6: Illustration of AUC calculation in discovery module of SURF. (a) Given a target genomic feature set (e.g., gene set or transcript set) and a descending ordering of all the features (e.g., ranked genes based on gene expression), the recovery curve (solid line) depicts the number of target genomic features among the top-ranked genomic features. For example, the set of top 510 features (marked by the vertical blue dashed line) harbors 40 target features. (b) Recovery curve in percentage scale. Both axes are re-scaled into [0, 1]. The blue area under the recovery curve (AUC) quantifies the activity level of target genomic set.

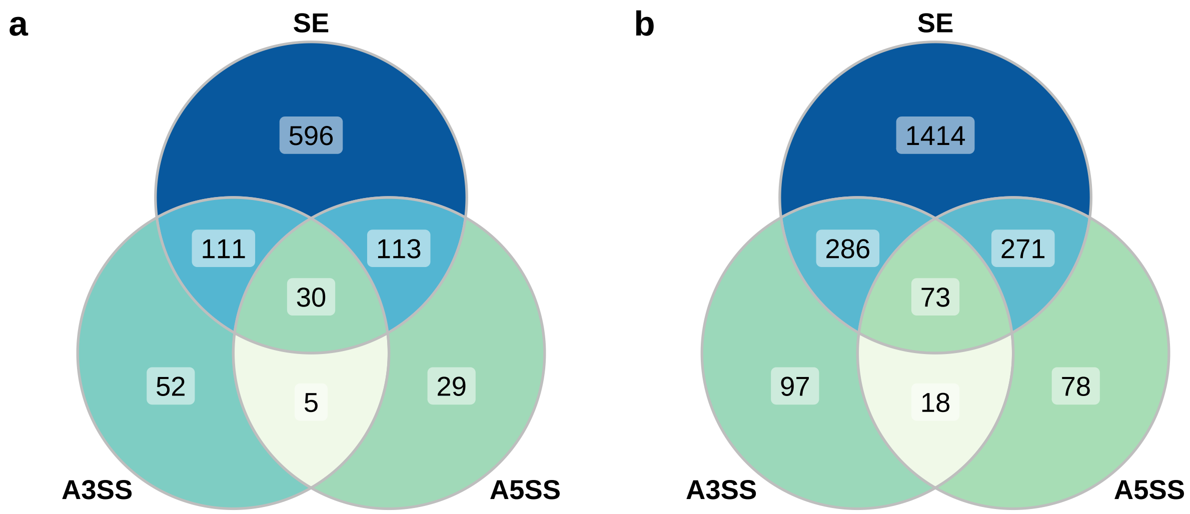


Supplementary Fig. S7: Annotation of local splice variations (LSV) detected by MAJIQ on simulated RNA-seq data (Materials and Methods) into ATR event types. MAJIQ annotates most LSVs into three ATR event types: SE, A3SS, and A5SS; however, mappings from LSVs to ATR event types are not unique. (a) ATR event type annotations of 936 (out of 974) LSVs detected with the default settings of MAJIQ. (b) ATR event type annotations of 2237 (out of 2323) LSVs detected with a liberal configuration of MAJIQ, where the change threshold is set to 0.1 (via option `--threshold`).

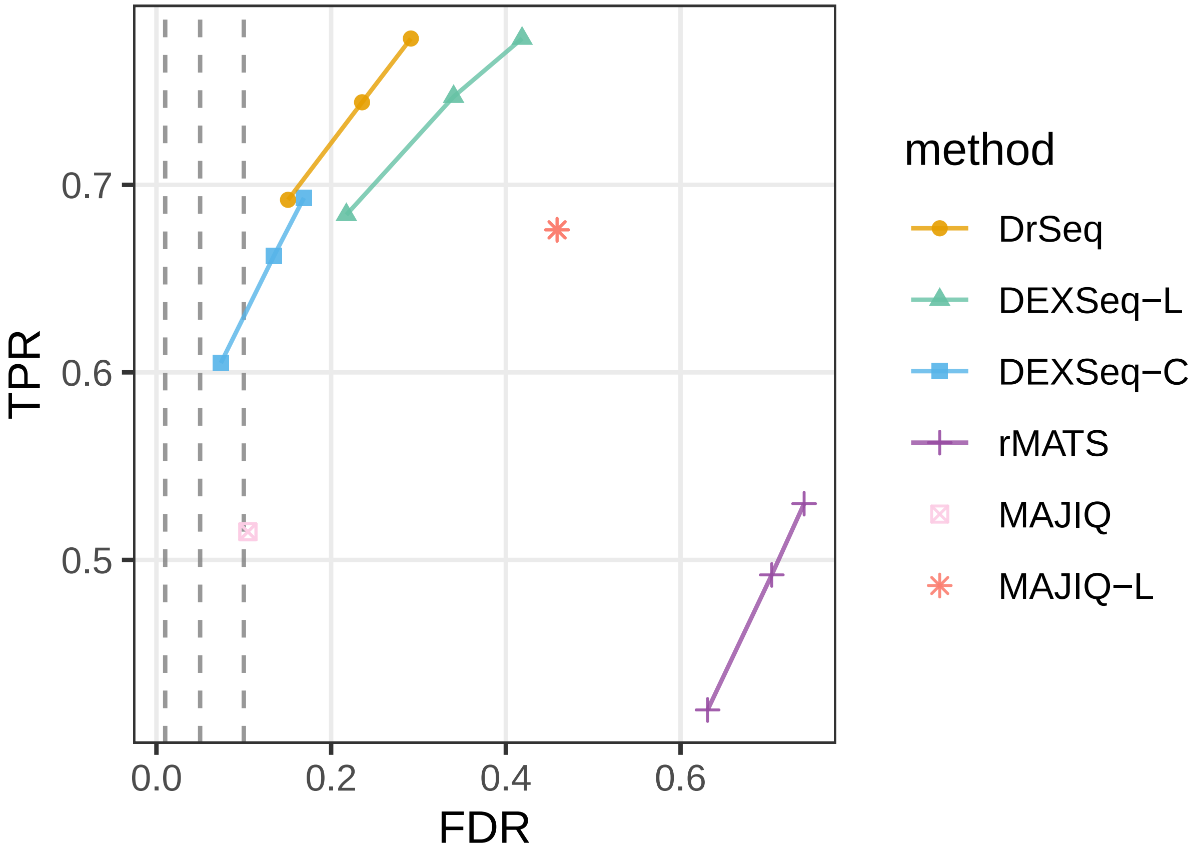


Supplementary Fig. S8: Comparison of the aggregate performances of differential ATR event detection methods DrSeq, rMATS, two simple strategies for stitching DEXSeq inferences into ATR level (DEXSeq-L and DEXSeq-C), MAJIQ (default), and MAJIQ-L (liberal as described in Fig. S7) using simulated RNA-seq data. For the first four methods, the three points display the true positive rate (TPR) and observed false discovery rate (FDR) at target FDR levels of 0.01, 0.05, and 0.1, respectively. The MAJIQ variants do not report (adjusted) p-values for the tested local splice variations, thus their performances are summarized as single points.

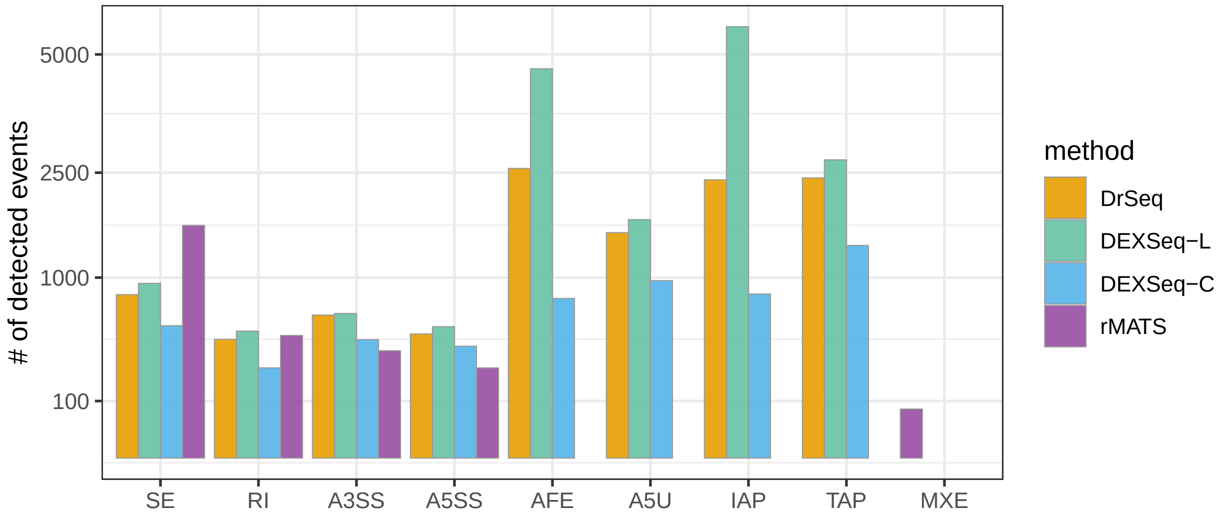


Supplementary Fig. S9: Comparison of the number of detected differential ATR events by various event-based methods in human transcriptome-based simulations. rMATS does not identify ATR event types AFE, A5U, IAP or TAP. In contrast, the only category excluded from DrSeq's ATR repertoire is MXE (mutually exclusive exons).

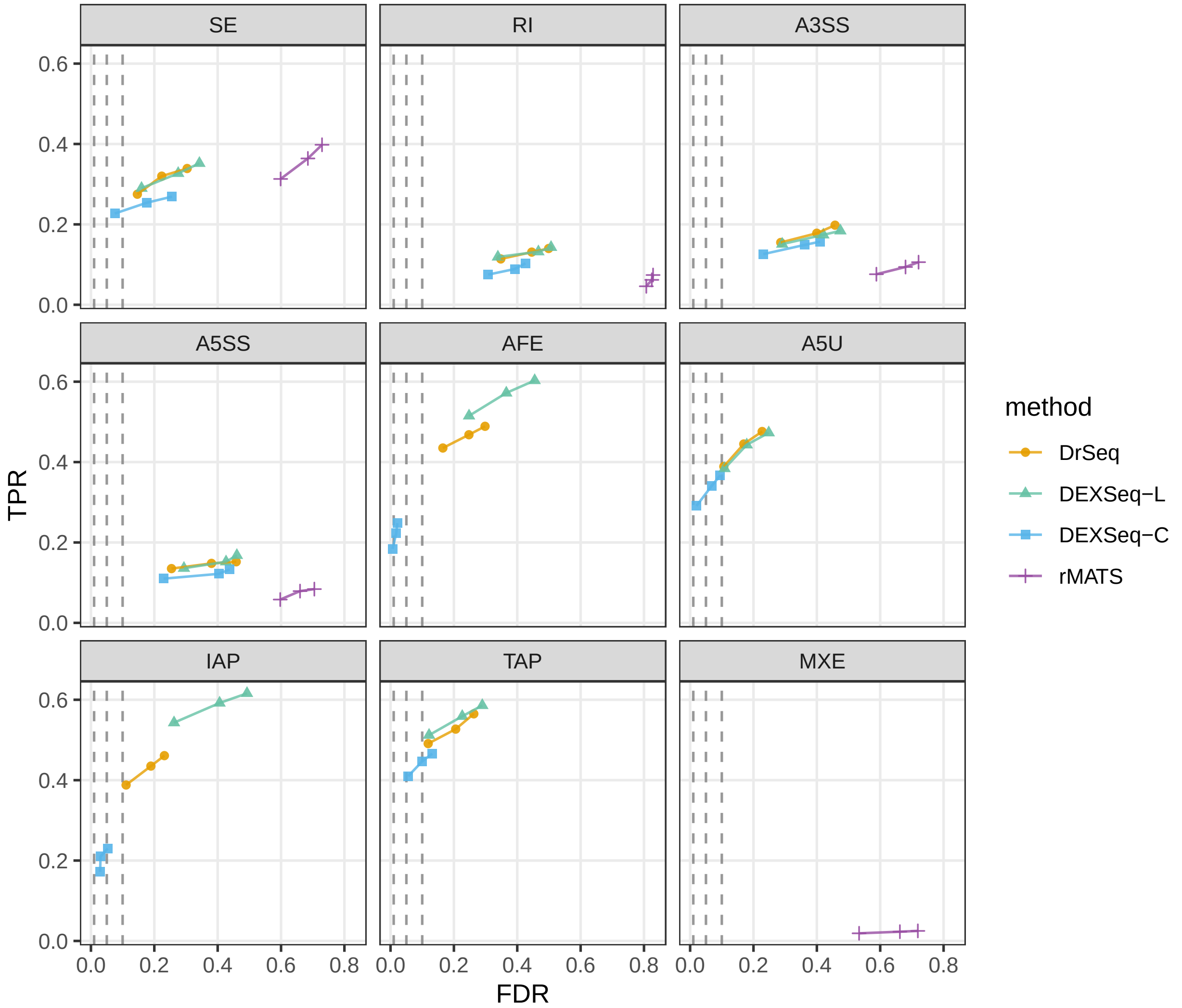


Supplementary Fig. S10: Event type specific comparison of the performances of DrSeq, rMATS, and two simple strategies for stitching DEXSeq inferences into ATR event level (DEXSeq-L and DEXSeq-C) using simulated RNA-seq data. For each method, the three points display the true positive rate (TPR) and observed false discovery rate (FDR) at target FDR levels of 0.01, 0.05, and 0.1, respectively. rMATS does not identify ATR event types AFE, A5U, IAP or TAP. In contrast, the only category excluded from DrSeq's ATR event construction is MXE (mutually exclusive exons).

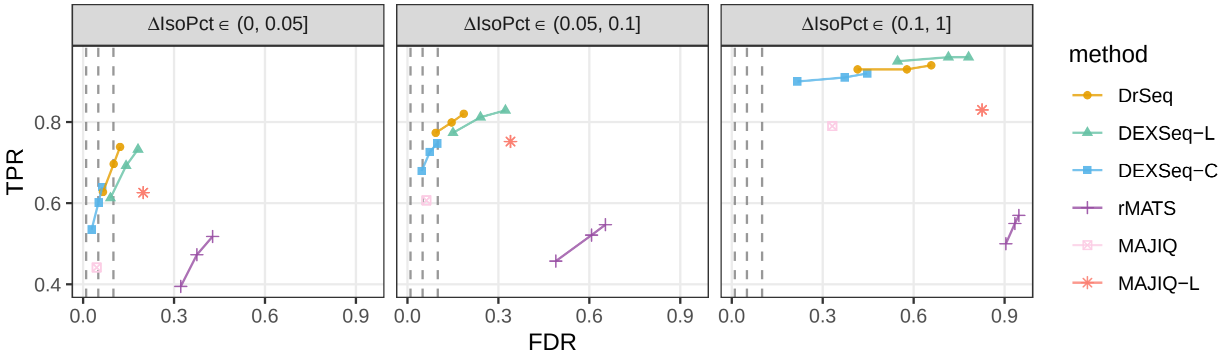


Supplementary Fig. S11: Comparison of the performances of DrSeq, rMATS, MAJIQ, and two simple strategies for stitching DEXSeq inferences into ATR event level (DEXSeq-L and DEXSeq-C) under different signal-to-noise levels. All 16,824 genes with multiple isoforms are divided into three groups by the difference in their isoform percentages (ΔIsoPct) of the most expressed isoforms (in TPM), namely (0, 0.05], (0.05, 0.1], and (0.1, 1]. The number of genes with altered isoform expression (Materials and Methods) in three groups are 666, 234, and 100, respectively. For each method, the three points display the true positive rate (TPR) and observed false discovery rate (FDR) at target FDR levels of 0.01, 0.05, and 0.1, respectively. Two MAJIQ variants do not directly report (adjusted) p-values for the tested local splice variations, thus their performance are each depicted as a single point.

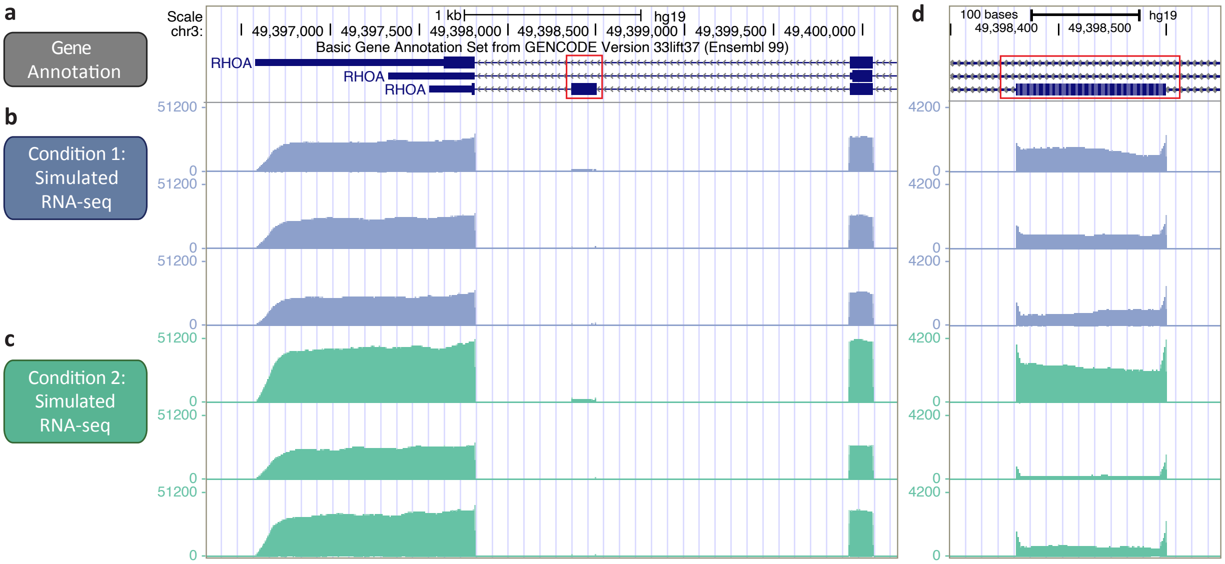


Supplementary Fig. S12: A false positive SE event detected by rMATS in the RHOA gene. (a) The 2nd last exon (red box) of RHOA gene model (Basic Gene Annotation from GENCODE 33 lift over to GRCh37), which resides in the negative strand is displayed. Each row depicts a single isoform of the gene. (b) The normalized read coverage of the gene in three replicates of RSEM-simulated RNA-seq for Condition 1. (c) The normalized read coverage of the gene in three replicates of RSEM-simulated RNA-seq for Condition 2. The expression levels (i.e., TPM values inputted to RSEM simulation) of the isoforms of the RHOA gene in (b) and (c) are set to be identical between the simulated conditions in (b) and (c), i.e., the second last exon is not differentially spliced. However, rMATS detected the SE event as differential (FDR $< 0.05$), while DrSeq did not. The detailed results of DrSeq and rMATS on the SE event are summarized in Tables S1 and S2. (d) A zoomed-in view of (a-c) for the skipped exon event (red box) with re-scaled y-axis.

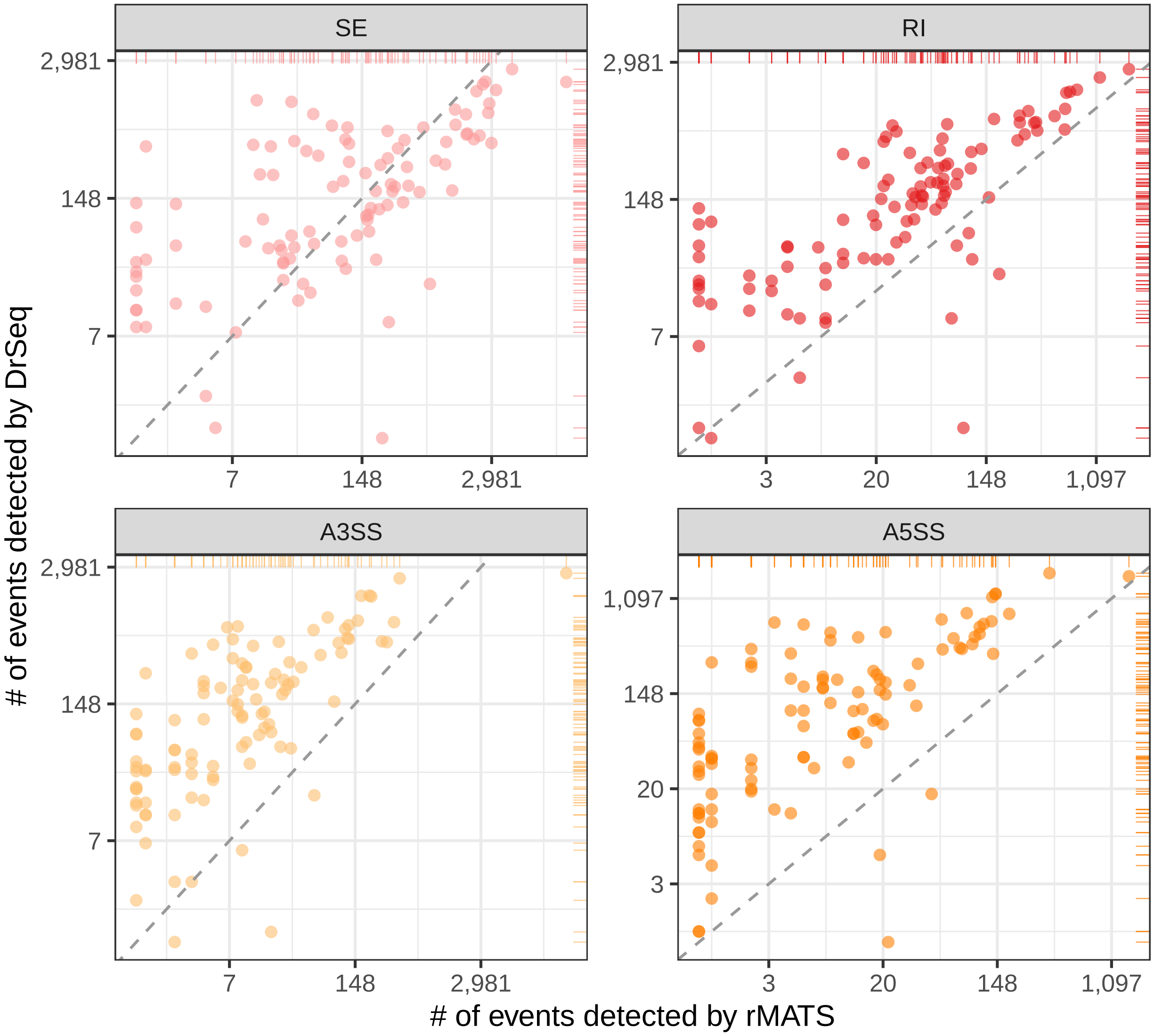


Supplementary Fig. S13: Comparison of differential AS detection by DrSeq and rMATS in 104 short-hairpin RNA knock-down followed by RNA-seq (with paired control) experiments in K562 cells. Each panel corresponds to one of the four common AS event types (SE, RI, A3SS, and A5SS) reported by both methods. Each point indicates the number of detected differential AS events (FDR $<$ 0.05) in one experiment (i.e., one RBP target) by rMATS (x-axis) and DrSeq (y-axis). The rMATS results were obtained from ENCODE portal (as of February 2020 and without surrogate variable analysis), and were based on rMATS software version 3.2.1.beta and human genome assembly version GRCh37.

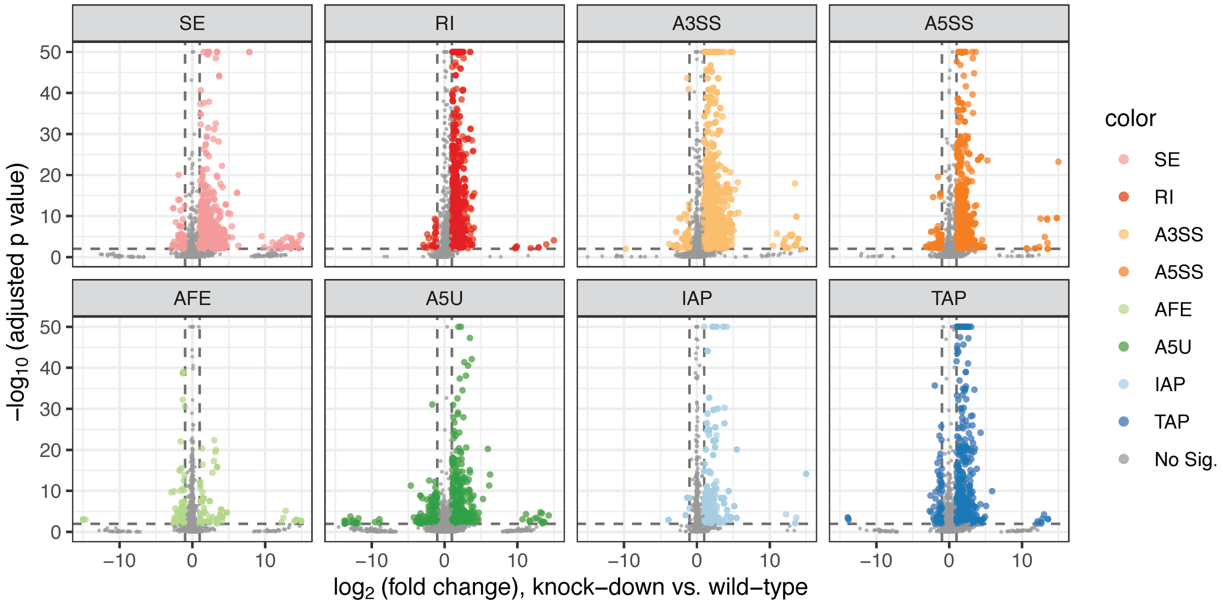


Supplementary Fig. S14: Volcano plot ($-\log_{10}$ transformed adjusted p-value versus $\log_2$ of fold change) of DrSeq results for AQR, stratified by ATR event types. Horizontal dashed line depicts FDR cut-off level of 0.01 and the vertical lines depict $\log_2$ fold change of 1 in absolute value.

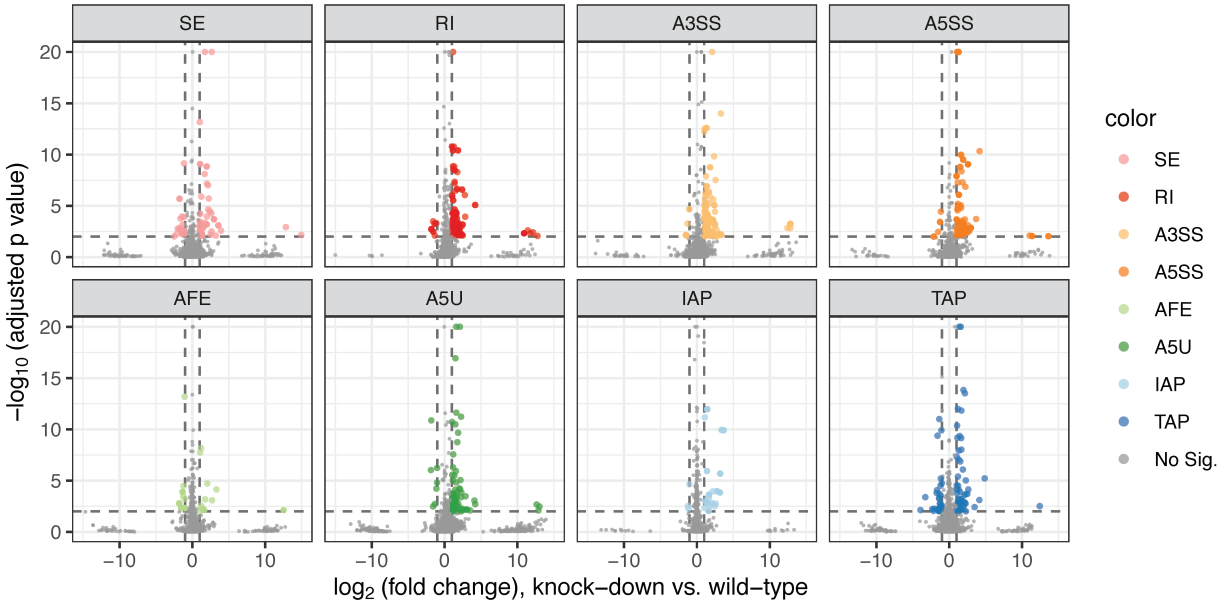


Supplementary Fig. S15: Volcano plot ($-\log_{10}$ transformed adjusted p-value versus $\log_2$ of fold change) of DrSeq results for SF3B4, stratified by ATR event types. Horizontal dashed line depicts FDR cut-off level of 0.01 and the vertical lines depict $\log_2$ fold change of 1 in absolute value.

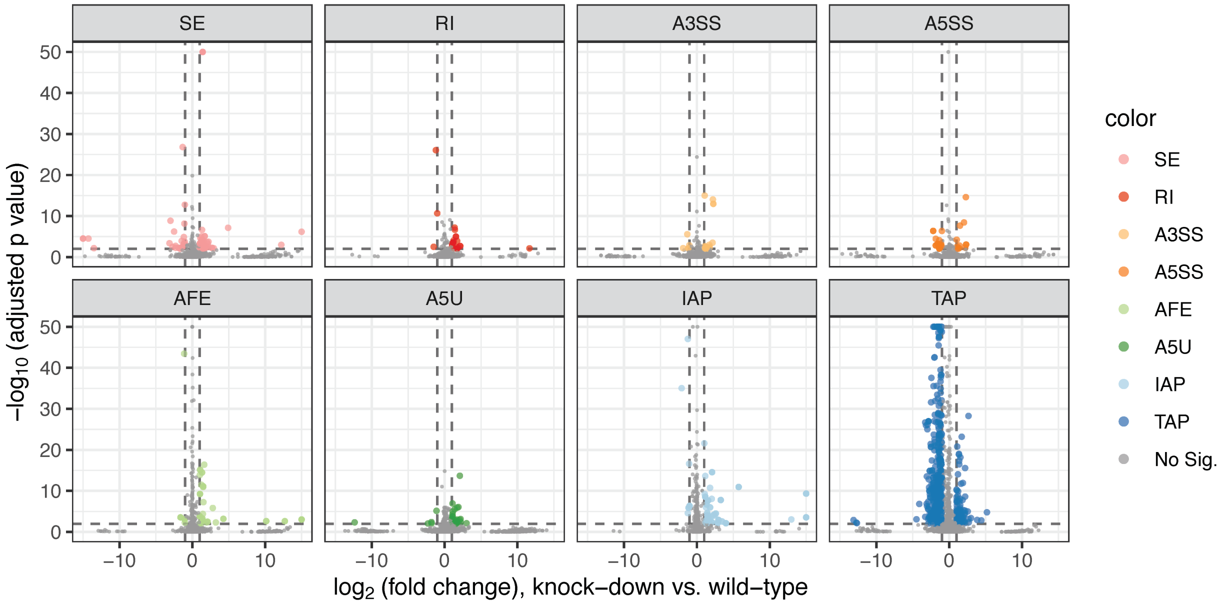


Supplementary Fig. S16: Volcano plot ($-\log_{10}$ transformed adjusted p-value versus $\log_2$ of fold change) of DrSeq results for CPSF6, stratified by ATR event types. Horizontal dashed line depicts FDR cut-off level of 0.01 and the vertical lines depict $\log_2$ fold change of 1 in absolute value.

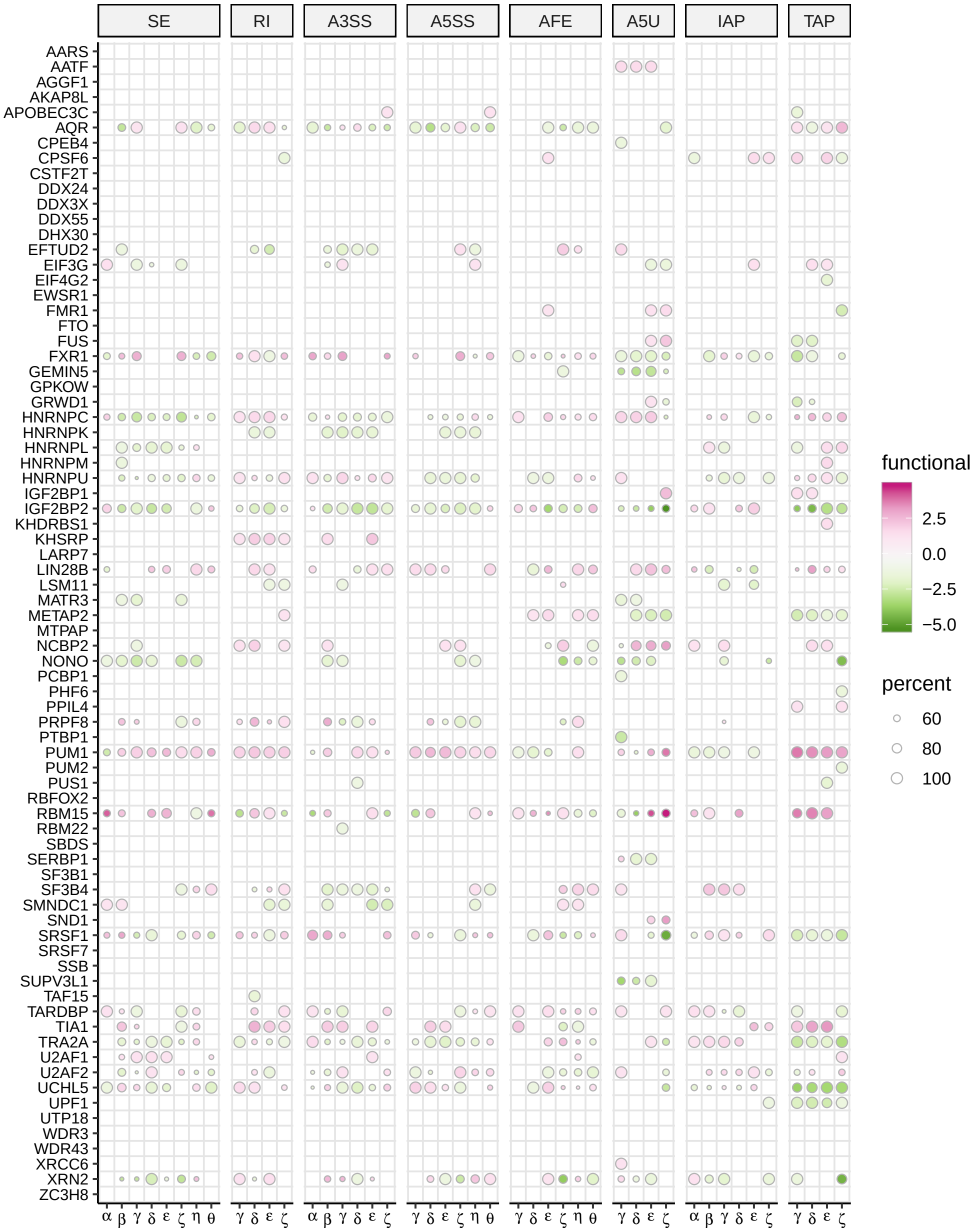


Supplementary Fig. S17: SURF-identified associations of location features with differential ATR across 73 RBPs (including the ones for which none of the feature locations significantly associated with the DrSeq-inferred differential ATR status). Each row depicts a single RBP and each column represents one location feature, grouped by the corresponding ATR event types (column strips). Each circle symbol in individual cells indicates a significant association at FDR of 0.05. The color of the circles represents inclusion (pink) or exclusion (green) and the fill-in densities are determined by $-\log_{10}$ transformed adjusted p-values from the association testing. For features with dual functions (i.e., binding of RBPs at these location features associates with both inclusion and exclusion), the circle size indicates the percentage of $-\log_{10}$ transformed adjusted p-value for the stronger association relative to the sum of both, i.e., smaller circles indicate similar associations for both inclusion and exclusion.

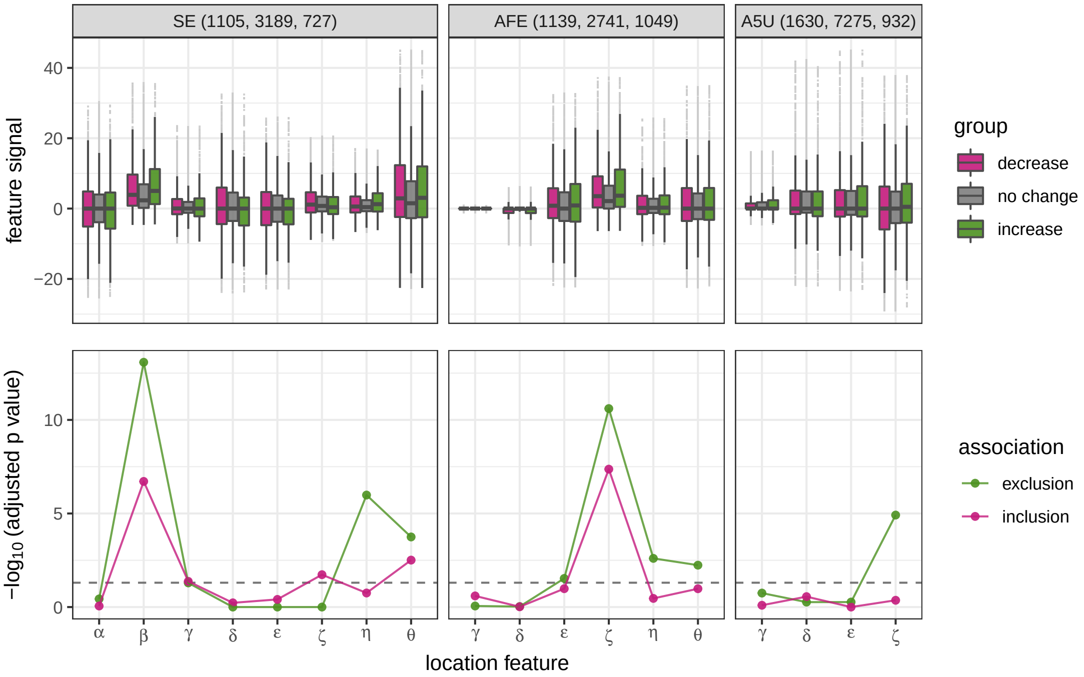


Supplementary Fig. S18: Functional association (FA) plot of AQR for event types SE, AFE, and A5U. Upper panel box plots display the distributions of feature signals among the three differential ATR groups (decrease, no change, increase). The numbers of ATR events in each group are reported in the parentheses at the top plot strip. The lower panels depict the $-\log_{10}$ transformed p-values for each tested association after multiplicity correction. The dashed lines indicate the FDR level of 0.05.

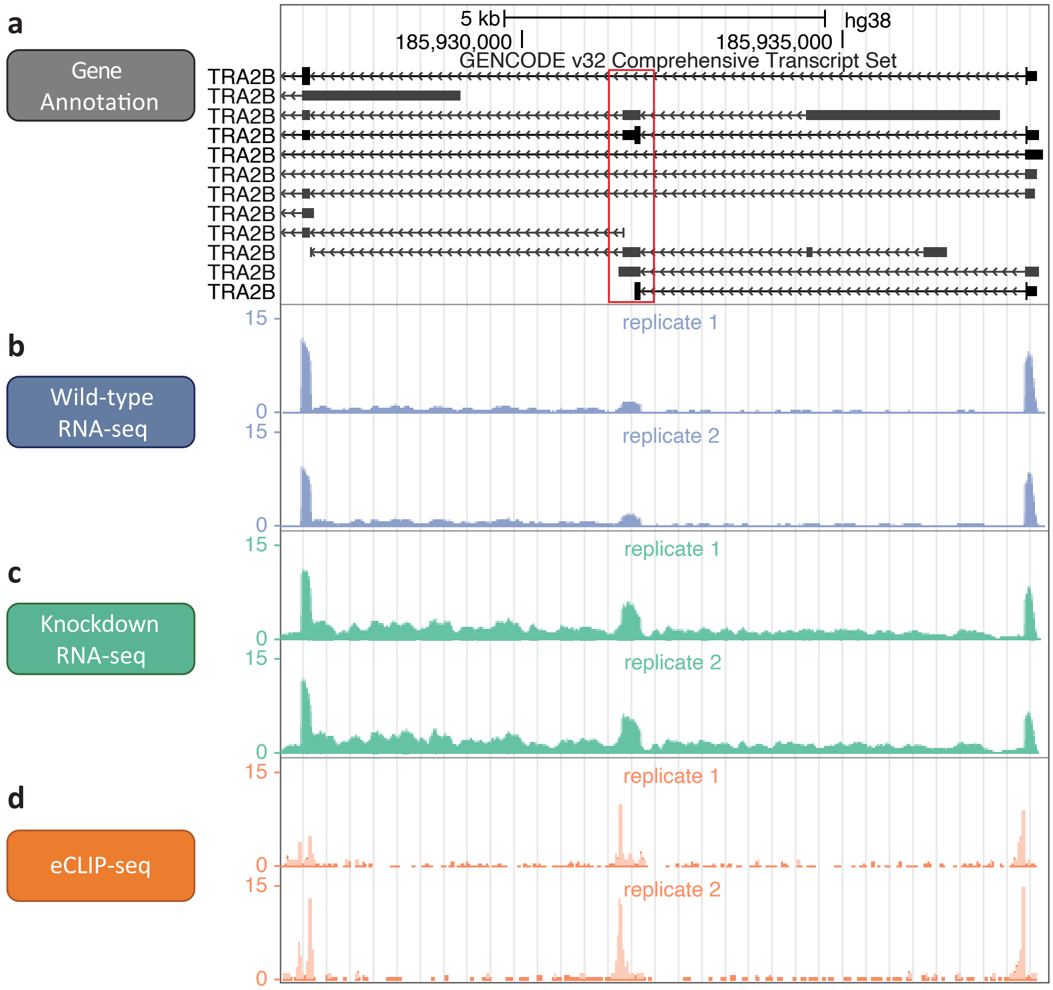


Supplementary Fig. S19: An included exon in the TRA2B gene upon AQR knock-down. (a) The 2nd exon (red box) of TRA2B gene model (GENCODE version 32) which resides in the minus strand. Each row depicts a single isoform of the gene. (b) The normalized read coverage in two replicates of wild-type RNA-seq. The normalized read coverage in two replicates of shRNA AQR knock-down followed by RNA-seq. The 2nd exon exhibits an increased relative exon usage when comparing the knock-down and wide-type conditions. (d) The normalized read coverage in two replicates of AQR eCLIP-seq.

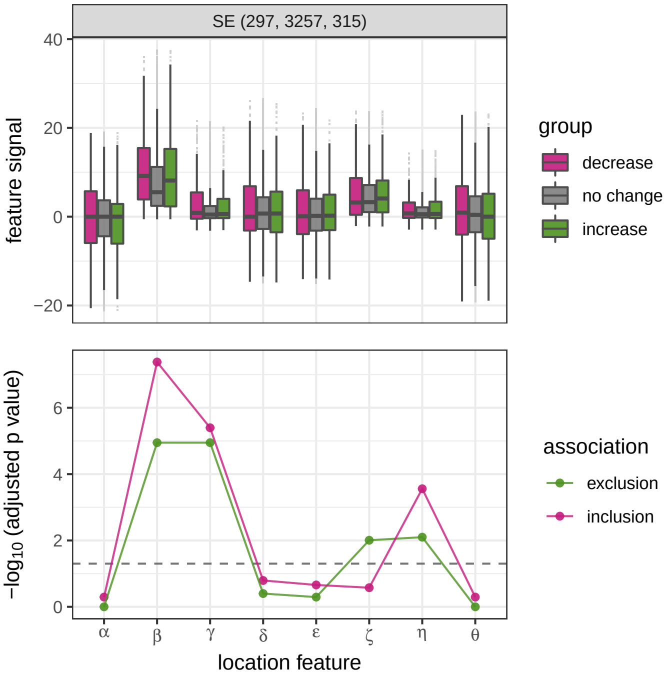


Supplementary Fig. S20: FA plot of PRPF8 for event type SE. See Fig. S18 for more information.

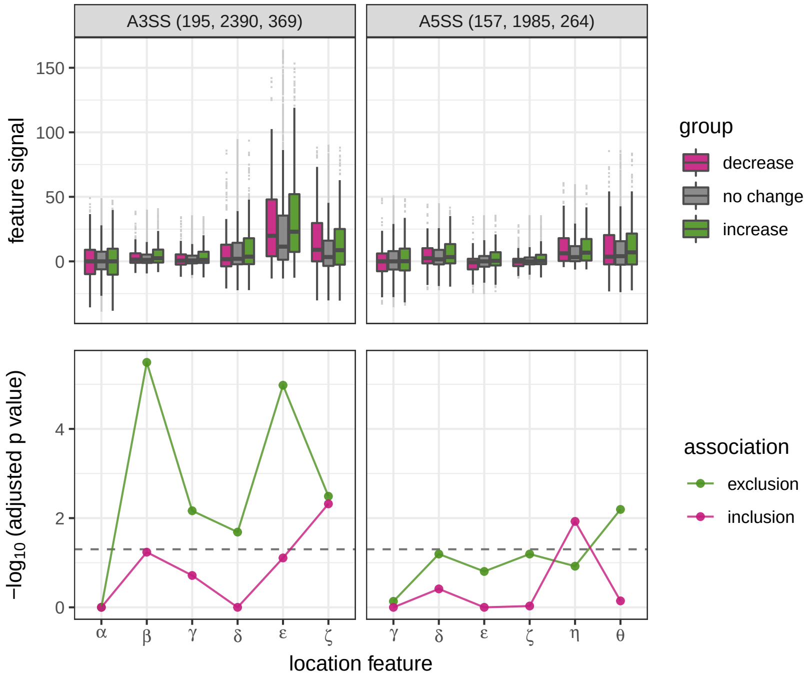


Supplementary Fig. S21: FA plot of SF3B4 for event types A3SS and A5SS. See Fig. S18 for more information.

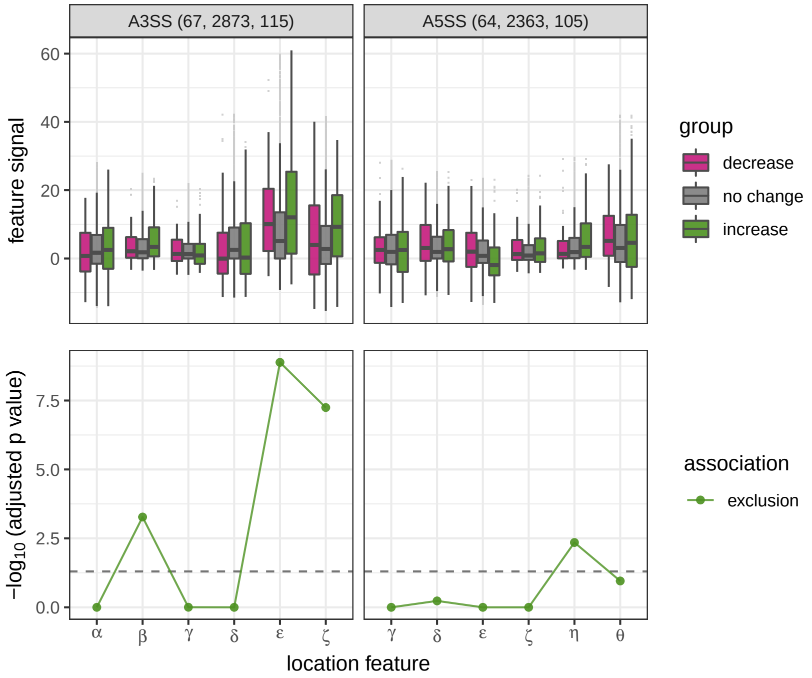


Supplementary Fig. S22: FA plot of SMNDC1 for event types A3SS and A5SS. See Fig. S18 for more information.

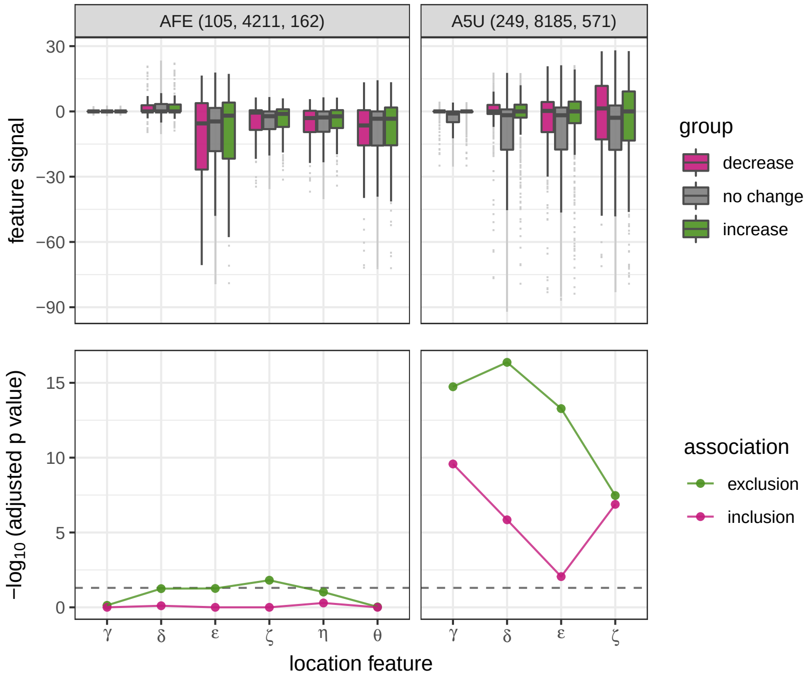


Supplementary Fig. S23: FA plot of GEMIN5 for event types AFE and A5U. See Fig. S18 for more information.

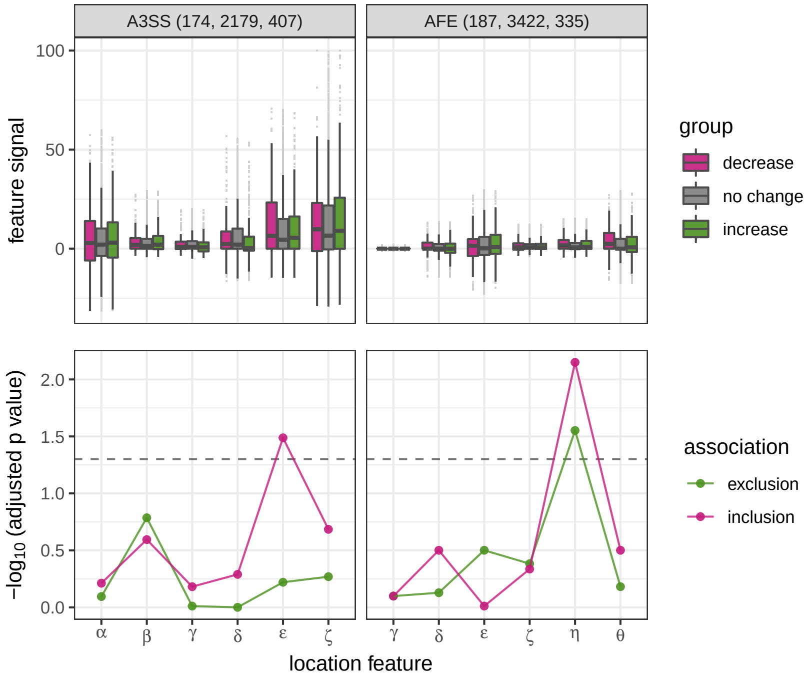


Supplementary Fig. S24: FA plot of U2AF1 for event types A3SS and AFE. See Fig. S18 for more information.

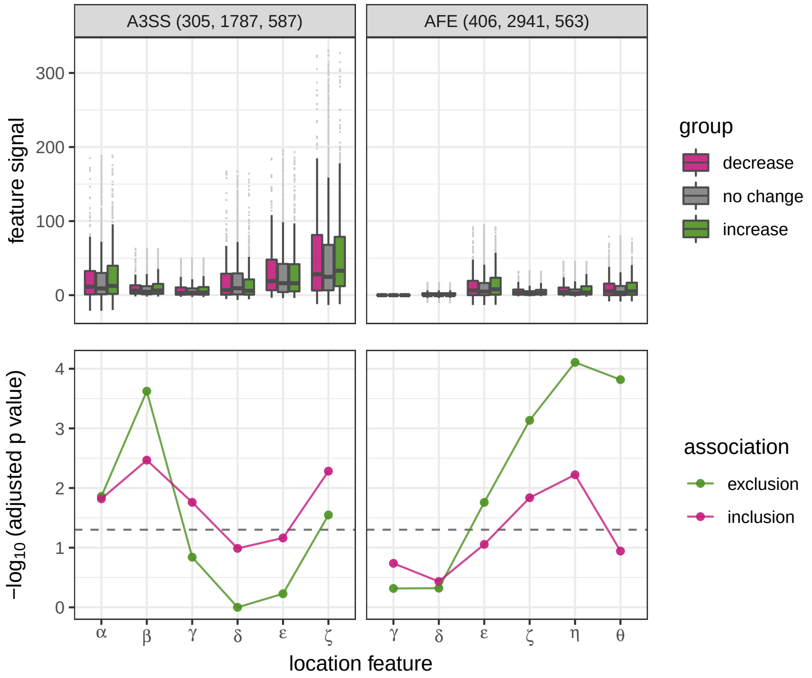


Supplementary Fig. S25: FA plot of U2AF2 for event types A3SS and AFE. See Fig. S18 for more information.

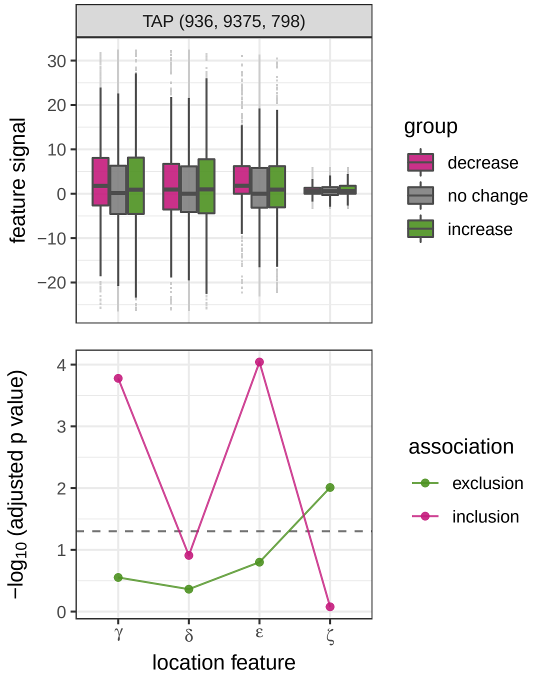


Supplementary Fig. S26: FA plot of CPSF6 for event type TAP. See Fig. S18 for more information.

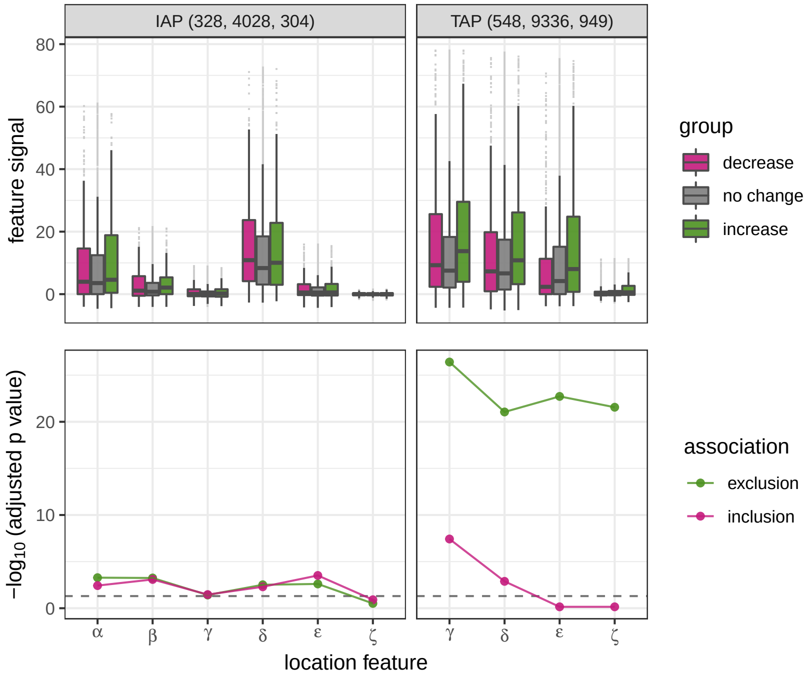


Supplementary Fig. S27: FA plot of UCHL5 for event types IAP and TAP. See Fig. S18 for more information.

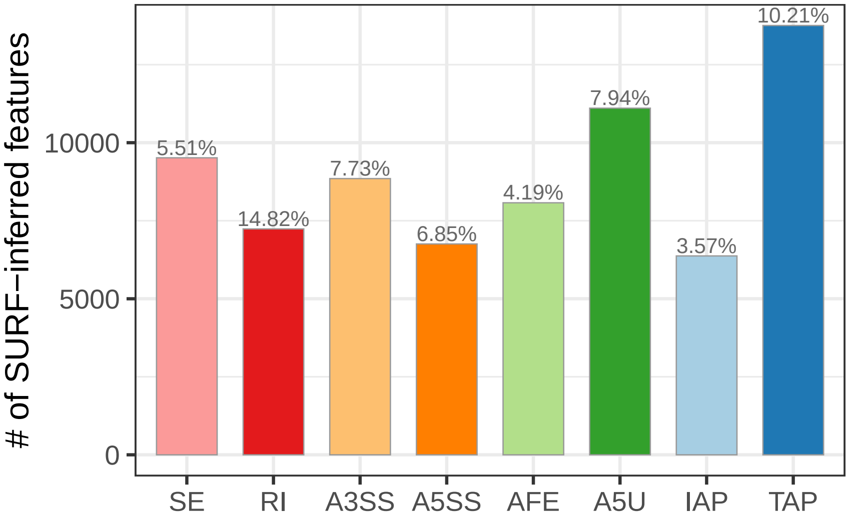


Supplementary Fig. S28: Numbers of SURF-inferred features (i.e., genomic locations for which an RBP binding is associated with differential ATR) for each ATR event category. The percentage (%) of SURF-inferred location features among all possible location features of the ATR event type is listed above each bar.

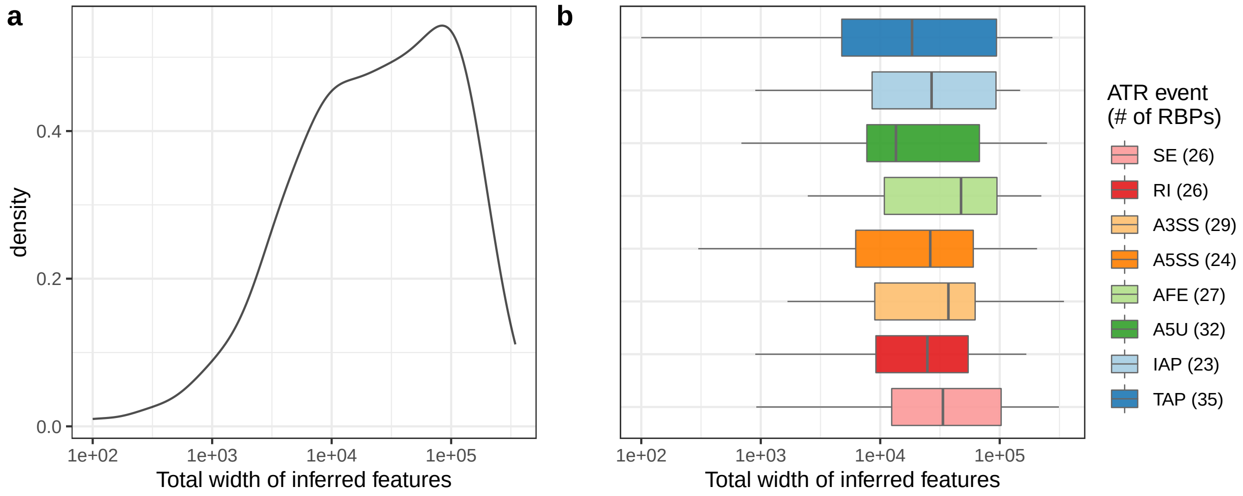


Supplementary Fig. S29: Distribution of the total length of SURF-inferred location features. Left panel density plot depicts the distribution over 53 individual RBPs. Right panel box plots depict the distributions stratified by ATR event types. For each event type, the number of RBPs with SURF-inferred location features is reported in parentheses with color legend.

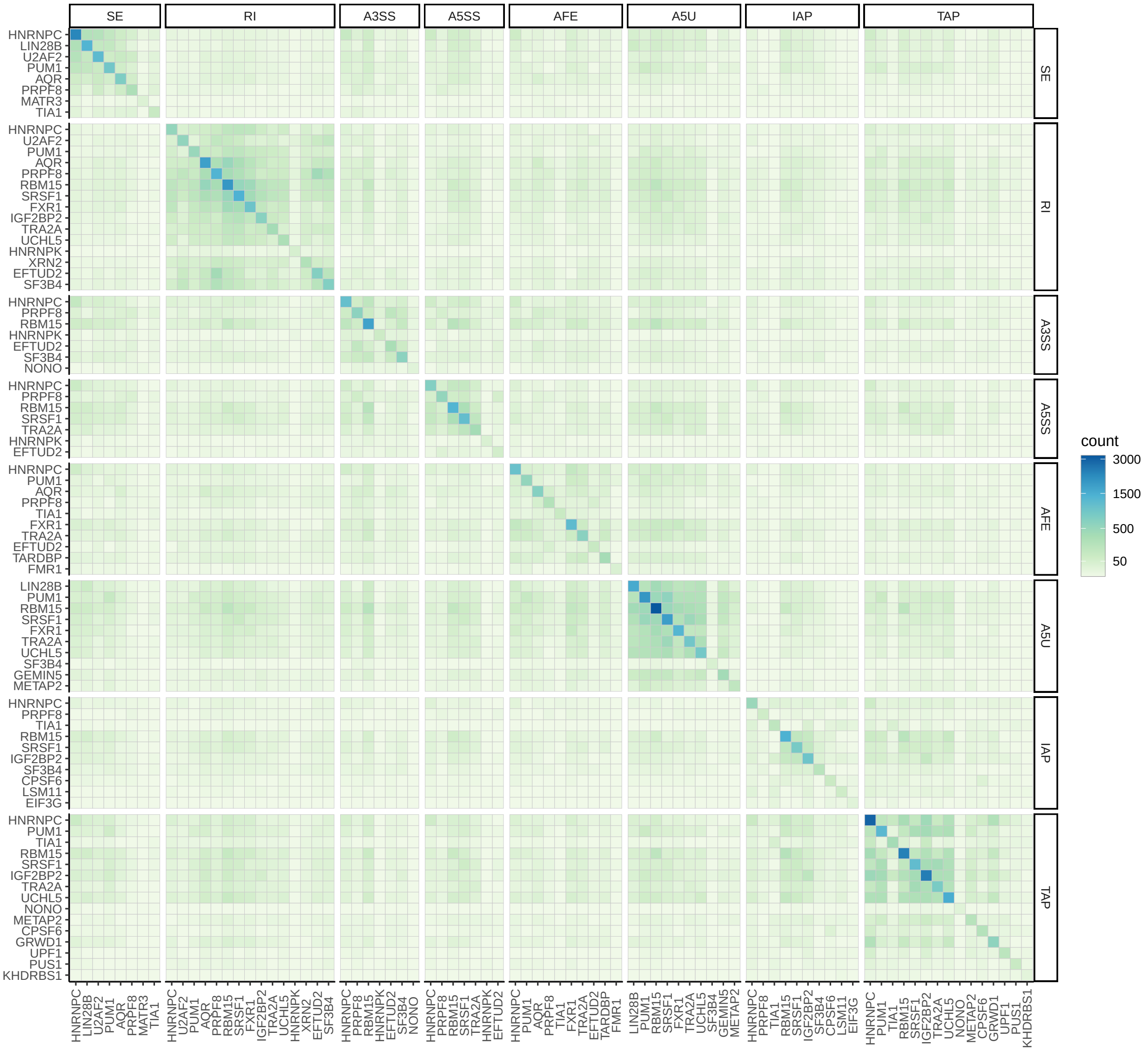


Supplementary Fig. S30: Clustering of RBP-ATR event types based on their numbers of shared SURF-inferred location features. The heatmap depicts a total of 82 RBP-ATR event types. Each entry reports the overlap between the two RBP-event type combinations. The diagonal color shade indicates the total number of SURF-inferred location features associated with each RBP-ATR event type combination. The off-diagonal color shades depict the numbers of shared location features between pairs of RBP-ATR event type combinations.

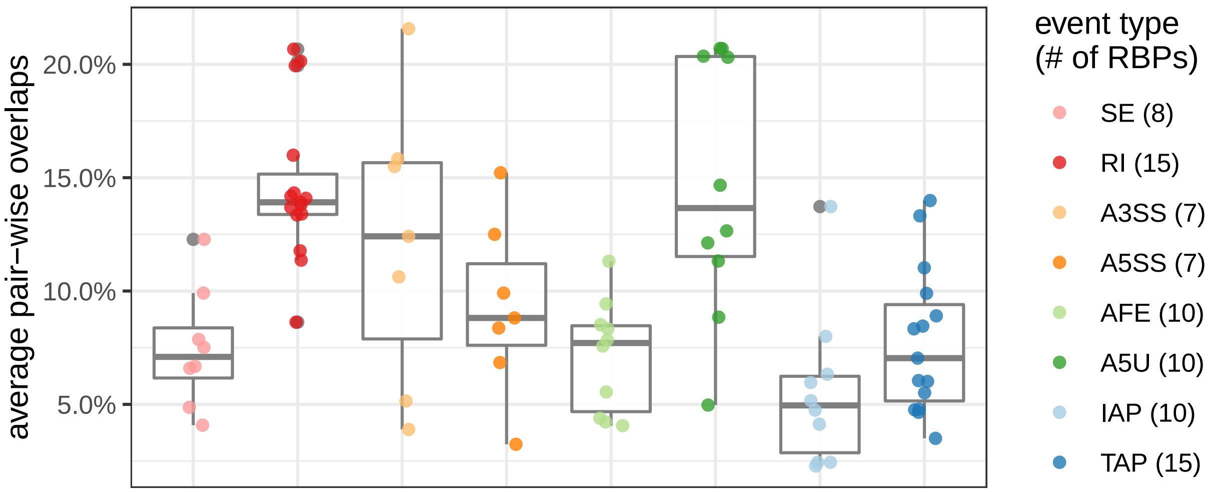


Supplementary Fig. S31: Summary of the extent of overlap between the SURF-inferred location feature sets of 82 RBP-ATR event type combinations. For each RBP-ATR event type combination, the corresponding SURF-inferred location feature set is compared with location feature sets of other RBPs within the same ATR event type. The average overlap is quantified by the average of percentages of its location features that are shared by other RBPs within the same event type. For each event type, the number of location feature sets (i.e., the number of RBPs) is reported in parentheses in the color legend.

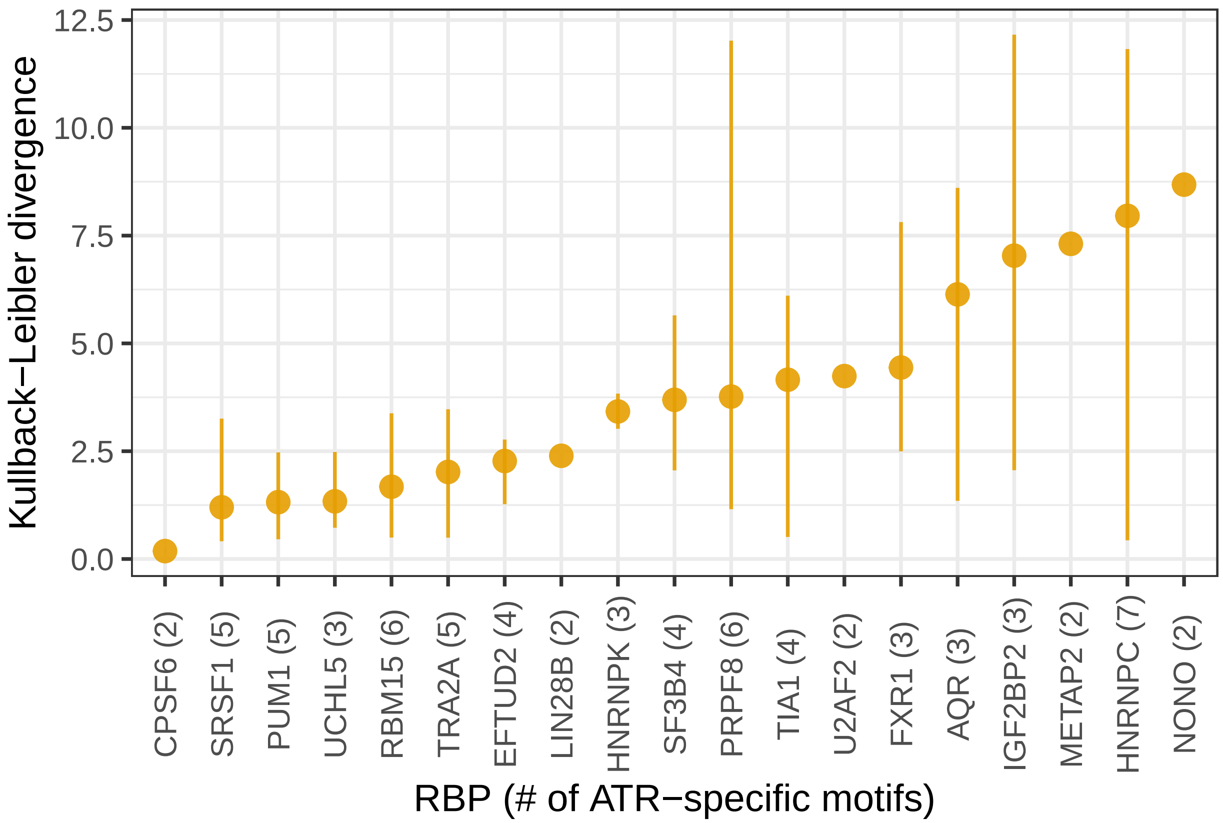


Supplementary Fig. S32: Diversity of multiple motifs identified for individual RBPs. Position weights matrices of the motifs (E-value $< 0.01$) identified by *de novo* sequence analysis of SURF-identified location features of RBPs across event types are compared with Kullback-Leibler (KL) divergence (y-axis). For each RBP, the number of significant motifs is reported in parenthesis in the x-axis label For each RBP, average diversity of all pairs of learnt motifs is reported along with the range (depicted by the vertical lines) indicating the minimum and maximum deviations observed across all pairs of motifs for the RBP.

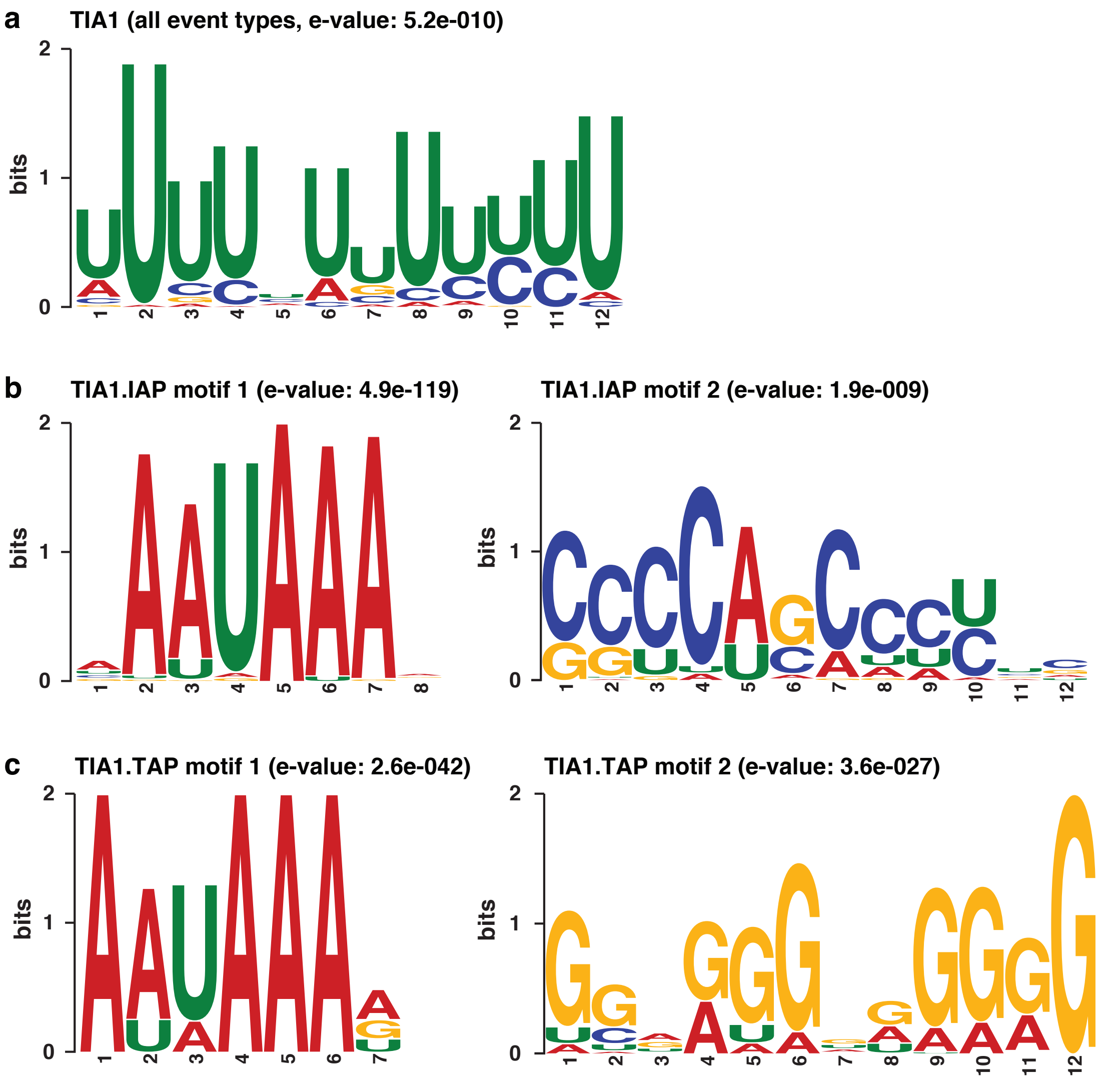


Supplementary Fig. S33: (a) Sequence logo of motif identified in 348 SURF-inferred location features of TIA for ATR. (b) Sequence logos of the two motifs identified in 161 SURF-inferred location features for TIA1 in regulation of IAP events. (c) Sequence logos of the two motifs identified in 144 SURF-inferred location features for TIA1 in regulation of TAP events. The total height of the letters depicts the information content of the position in bits.

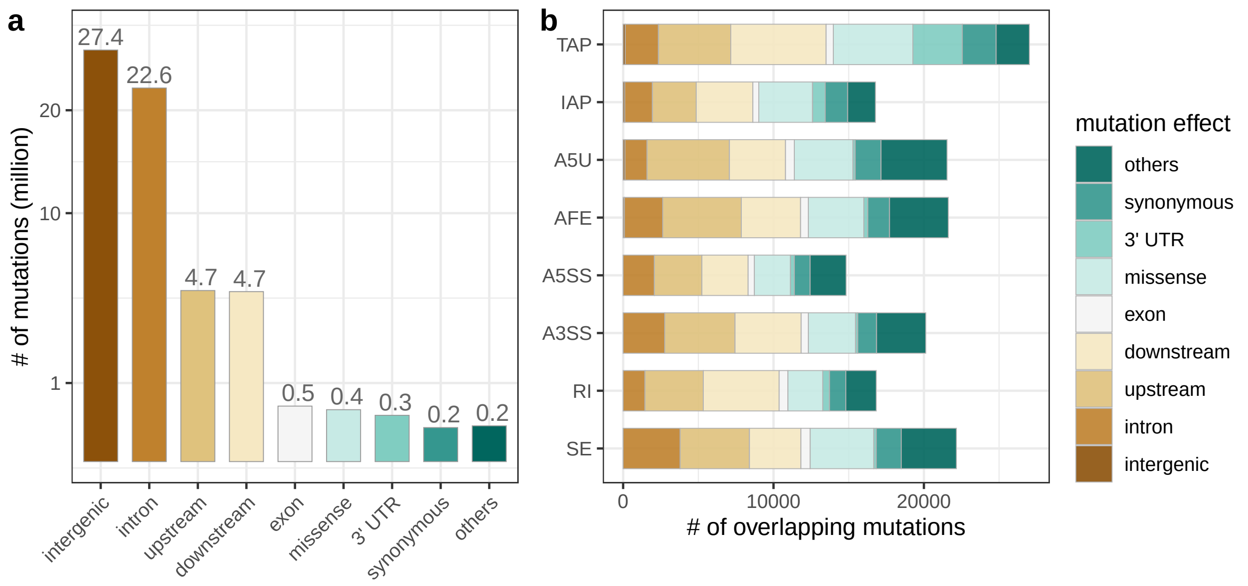


Supplementary Fig. S34: (a) Genomic location annotations of somatic mutations from ICGC (non-US) projects. The percentage of mutations in each category is listed above each bar. (b) The composition of ICGC mutation effects overlapping with SURF-inferred location features based on 54 RBPs. The class of intergenic mutations has fewest overlaps.

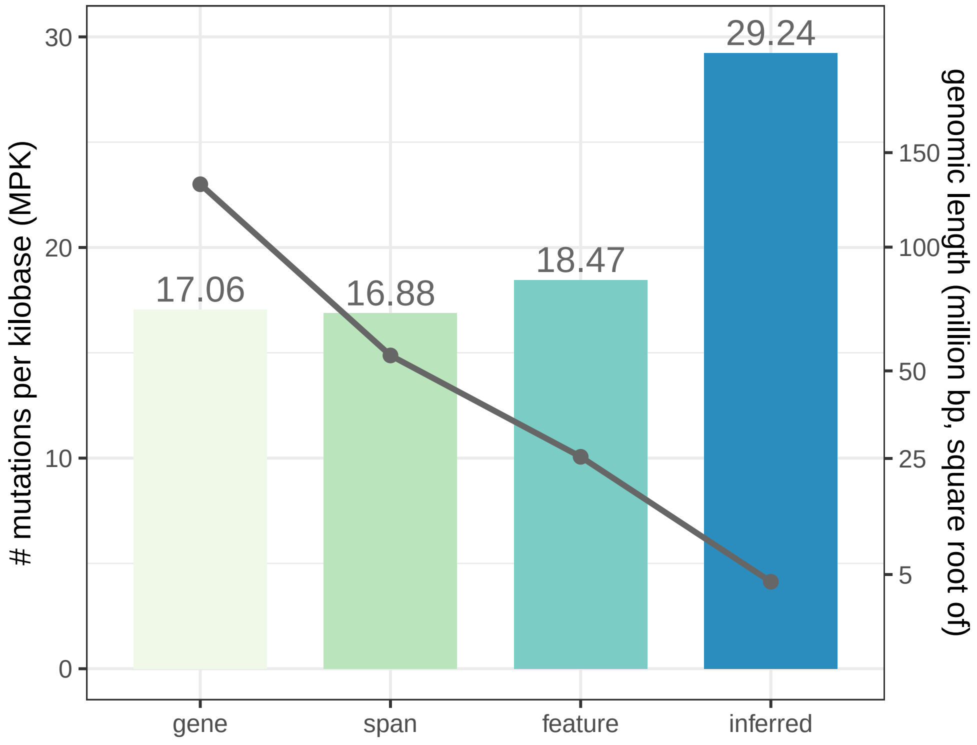


Supplementary Fig. S35: Mutations per kilobase (MPK) of SURF-inferred features (inferred) and three control genomic sets (gene, span, and feature). The bars depict the MPK quantified with somatic mutations from the TCGA project. The solid line with the right-hand side axis depicts the total length of the genomic regions considered.

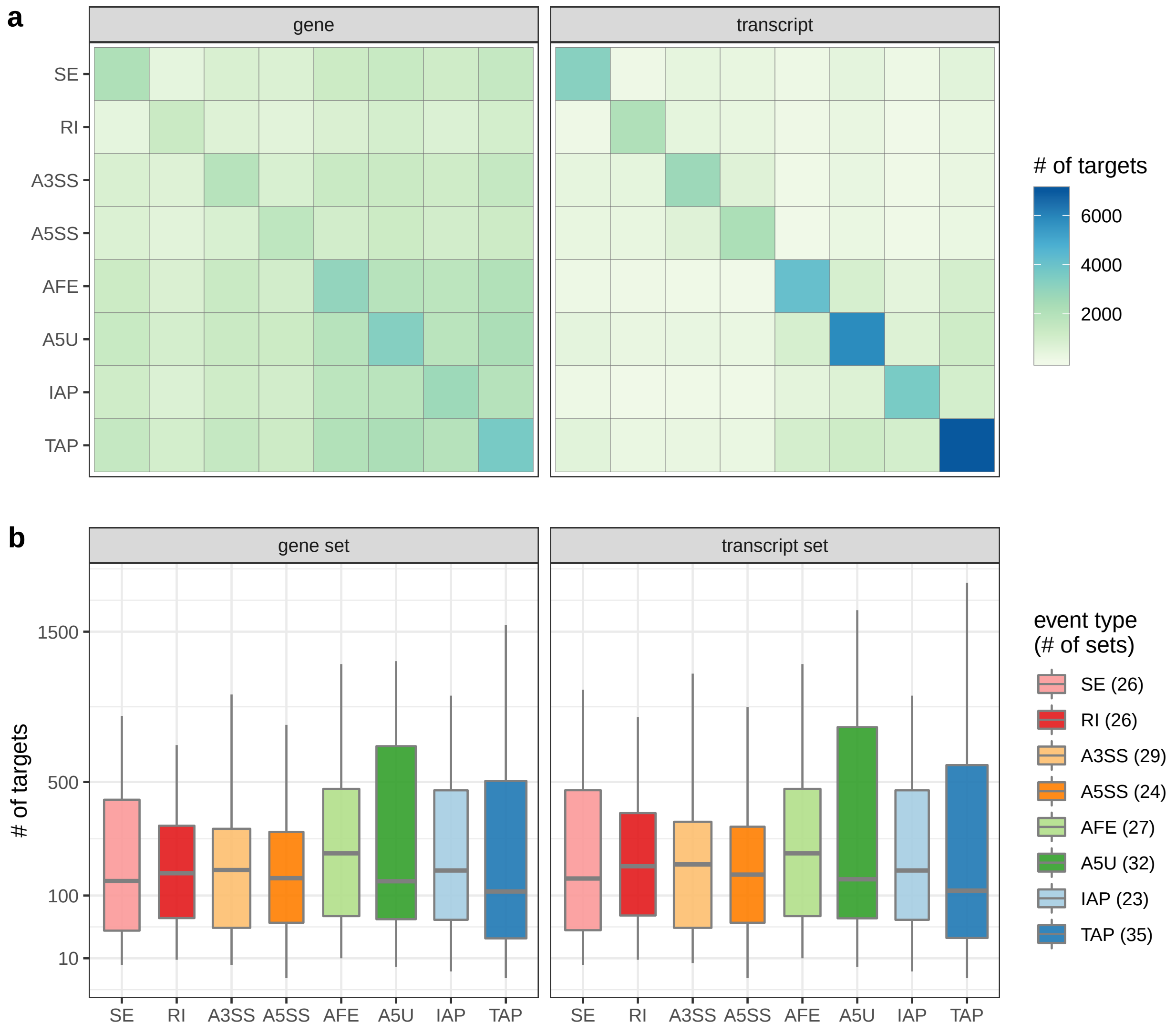


Supplementary Fig. S36: The number of SURF-inferred target genes and target transcripts harboring SURF-inferred location features of 52 RBPs. (a) The number of target genes (left) and transcripts (right) shared between pairs of ATR event types. The number of targets belonging to each ATR event types is depicted in diagonal. (b) The size of target gene sets (left) and transcript sets (right) grouped by associated ATR event type. For each event type, the number of target sets is reported in parentheses in the color legend.

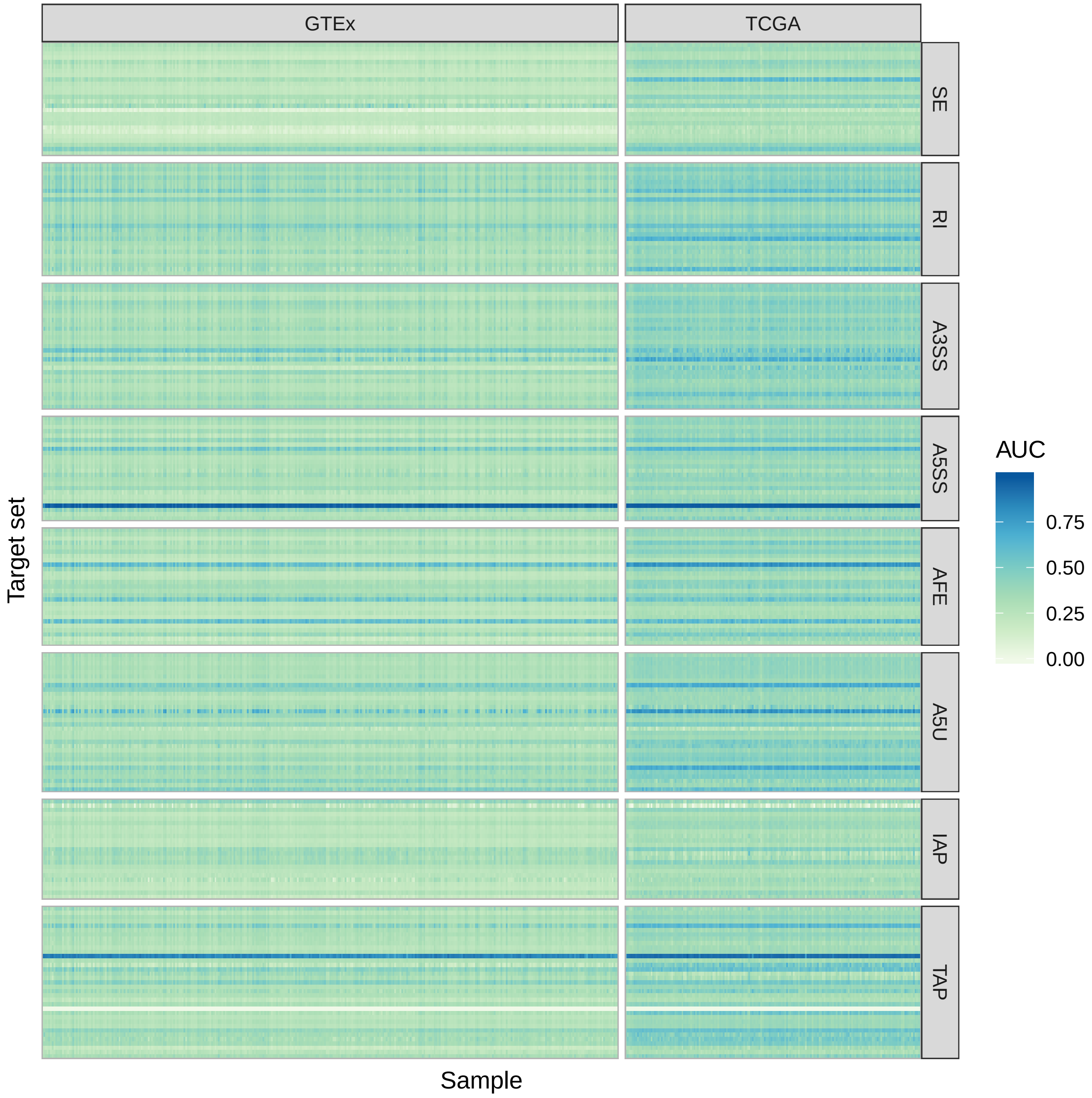


Supplementary Fig. S37: Heatmap of the observed AUC for 222 gene sets (row) in 337 GTEx whole blood samples and 173 TCGA LAML samples (column). Row strips indicate the eight corresponding ATR event types of the gene sets.

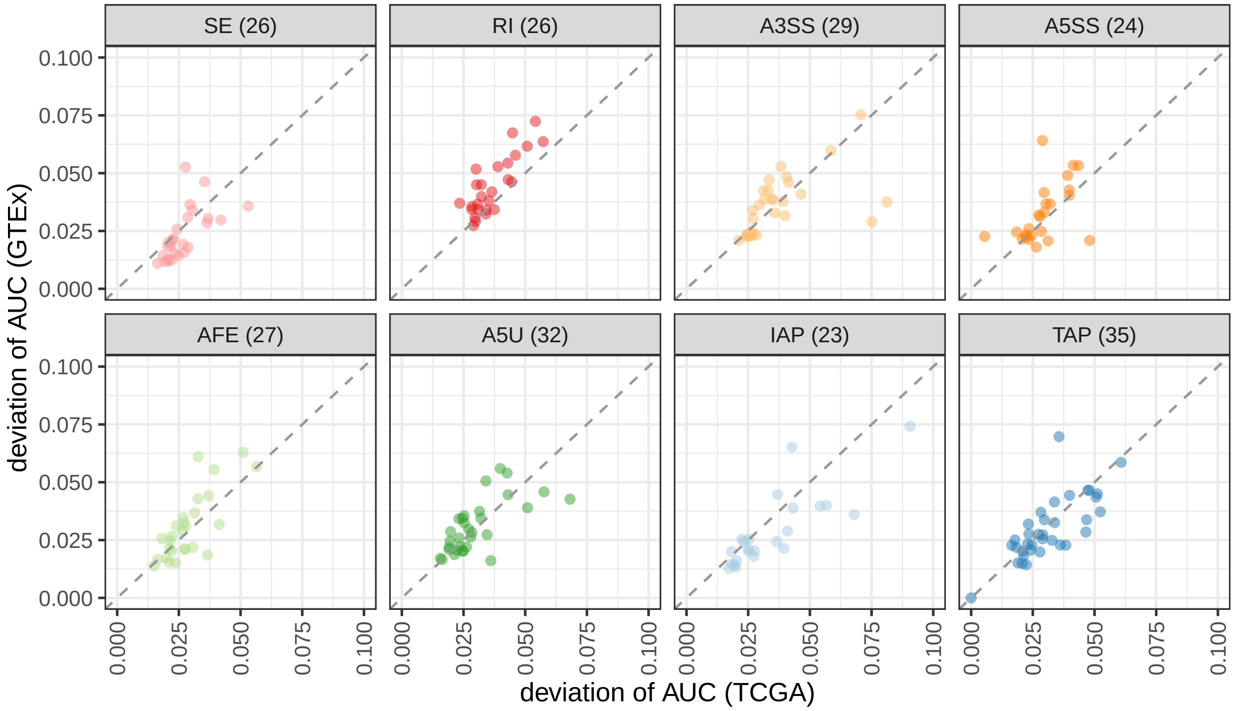


Supplementary Fig. S38: The variation of empirical AUC for 222 target gene sets harboring SURF-identified location features, stratified by associated ATR event type. Each point indicates, for one gene set, the standard deviation of AUC in 173 LAML (TCGA) samples and in 337 whole blood (GTEx) samples. The diagonal dashed line in each panel indicates equal variance between two projects. For each event type, the number of target gene sets is reported in parentheses within the top panel strips.

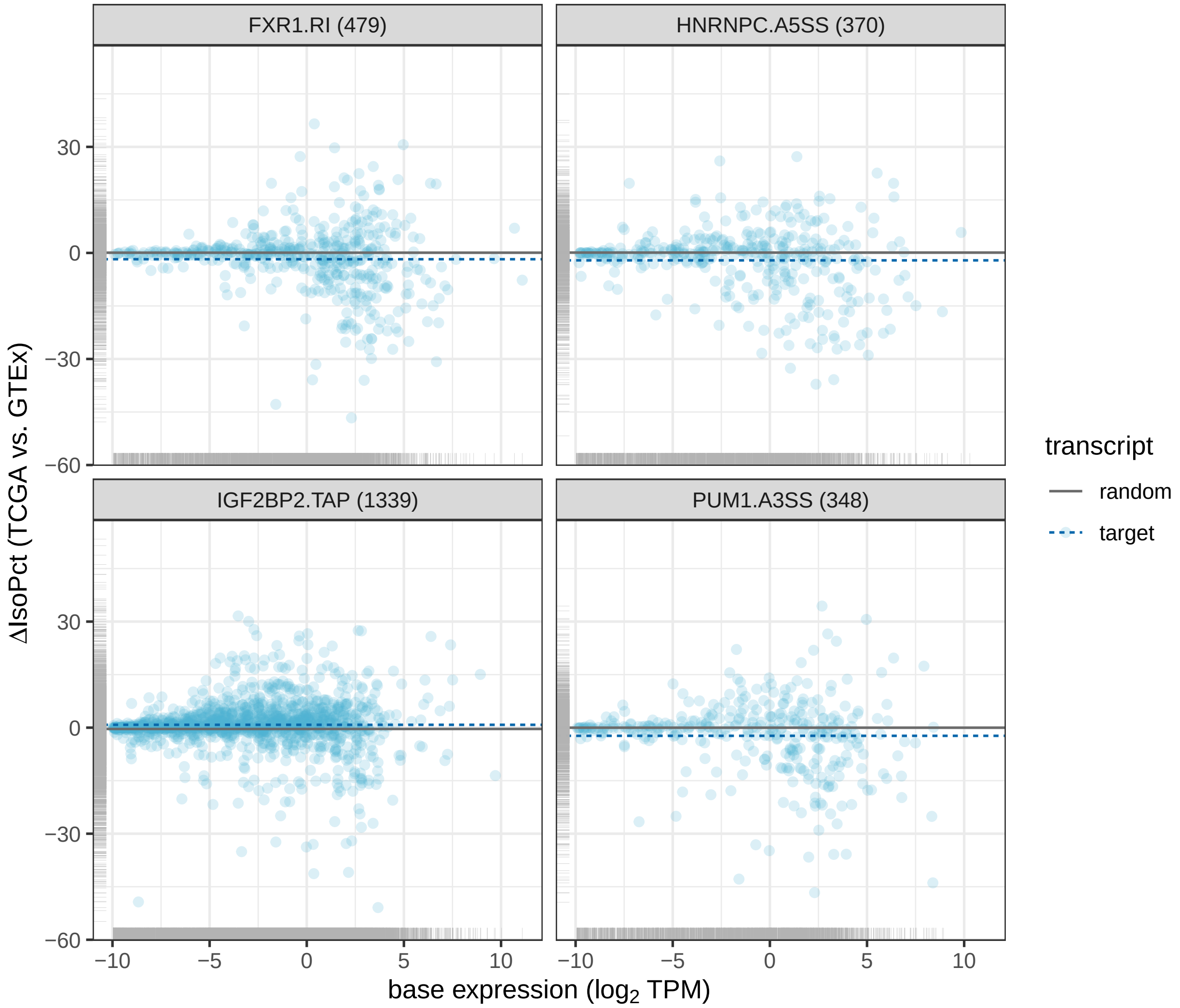


Supplementary Fig. S39: Comparison of target transcript expression changes in GTEx whole blood and TCGA LAML RNA-seq samples. The four panels depict SURF-inferred transcript targets of (i) FXR1 in RI, (ii) HNRNPC in A5SS, (iii) IGF2BP2 in TAP, and (iv) PUM1 in A3SS. Each point displays the base expression (in log 2 transformed TPM) and the difference in isoform percent (ΔIsoPct) from normal (GTEx) to tumor (TCGA) samples. The number of transcripts in each target set is reported in parentheses in the top panel strips. A dashed blue line indicates the average ΔIsoPct across the target transcript set, and a solid grey line indicates the average ΔIsoPct across the randomized control transcript sets.

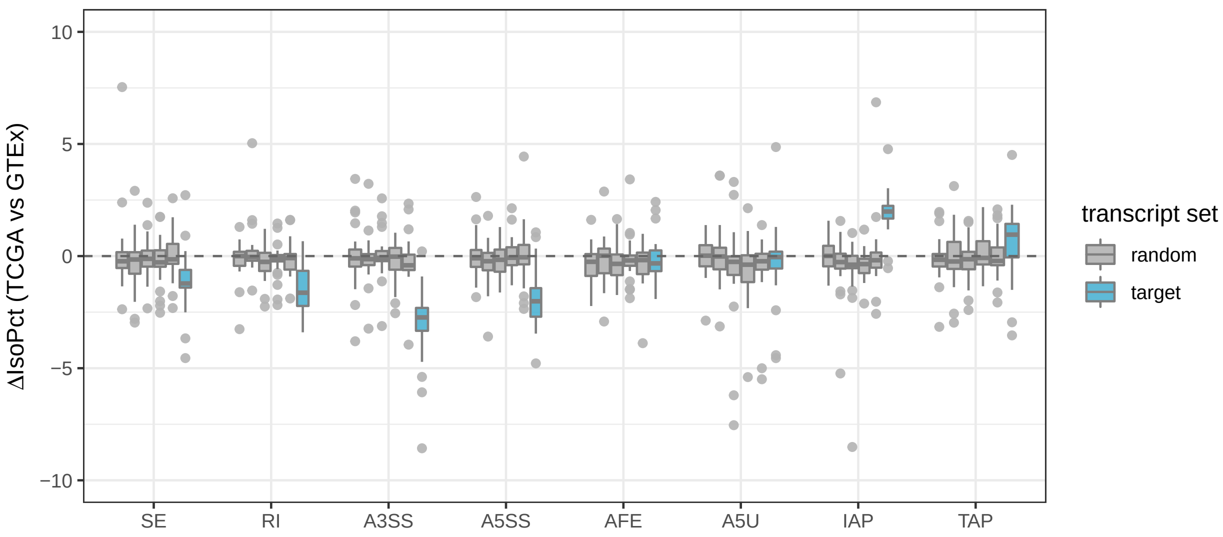


Supplementary Fig. S40: Comparison of the average changes in isoform percentages of transcript sets between TCGA LAML and GTEx whole blood samples, with multiple replicates of control transcript sets, across ATR event types. Boxplots are over either target transcript sets (blue) or randomized control transcript sets (grey, 5 sets of controls for each event type). The differences in isoform percentages (ΔIsoPct) between TCGA LAML and GTEx whole blood samples are averaged within individual transcript sets. 222 transcript sets are depicted in one of eight associated ATR event types.

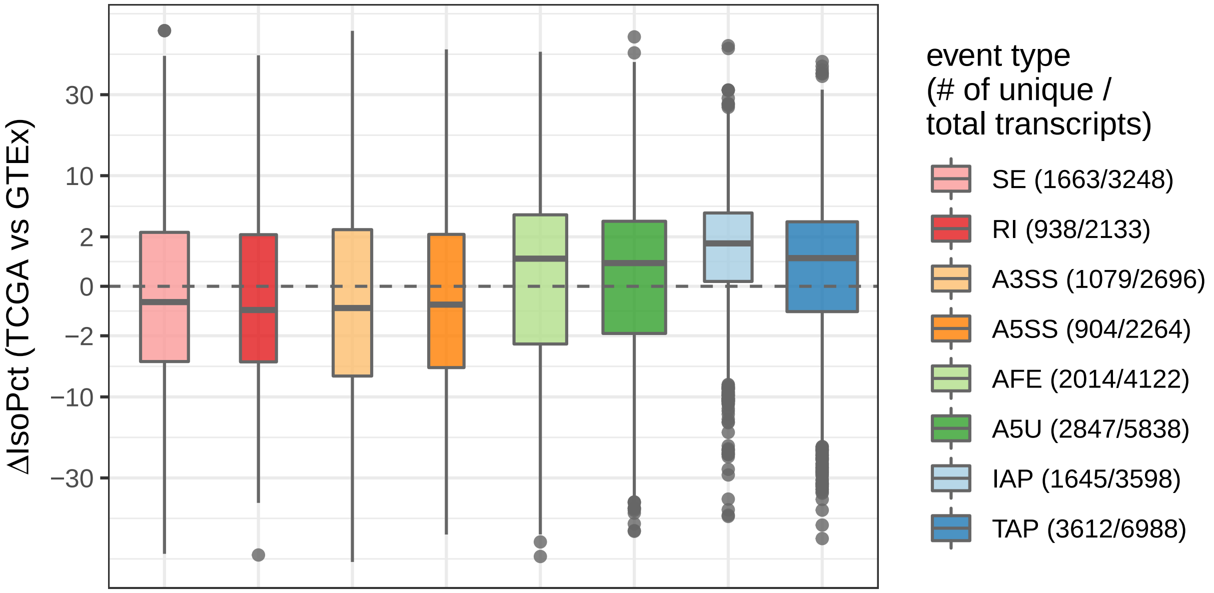


Supplementary Fig. S41: Comparison of ΔIsoPct (see Fig. S40) for a subset of SURF-inferred target transcripts that uniquely associated with only one ATR event type. The numbers of transcripts uniquely associating with each event type are reported in parentheses in the color legend along with the total numbers of transcripts of the event types and are also depicted by the width of the box plots. The differences in the ΔIsoPct of the four classes of AS events (SE, RI, A3SS, and A5SS) are not statistically significant (adjusted p-values $> 0.05$ in pairwise t-tests). Similarly, the differences between the two ATI event types (AFE and A5U) are not statistically significant (adjusted p-value $> 0.05$). Remaining pairwise comparisons of ΔIsoPct values between ATR event types (e.g., between AS and APA event types) are significantly different (adjusted p-values $< 0.01$)

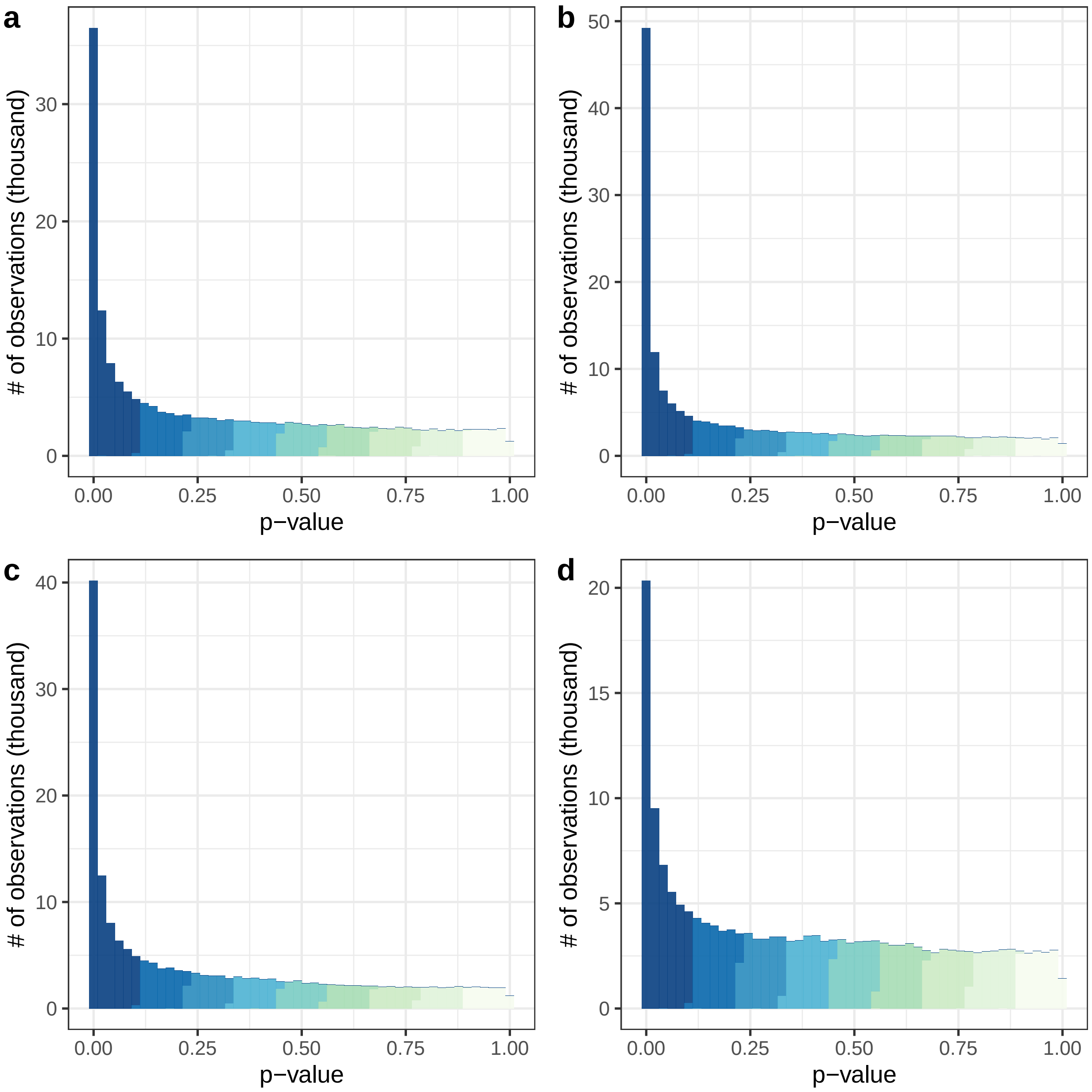


Supplementary Fig. S42: Histogram of raw p-values from the differential ATR analysis (DrSeq) of ENCODE datasets (a) AQR, (b) SRSF1, (c) CPSF6, and (d) SF3B4. Histograms display a mixture of uniform distribution on $[0, 1]$ and a point mass at zero, suggesting well-calibrated p-values.

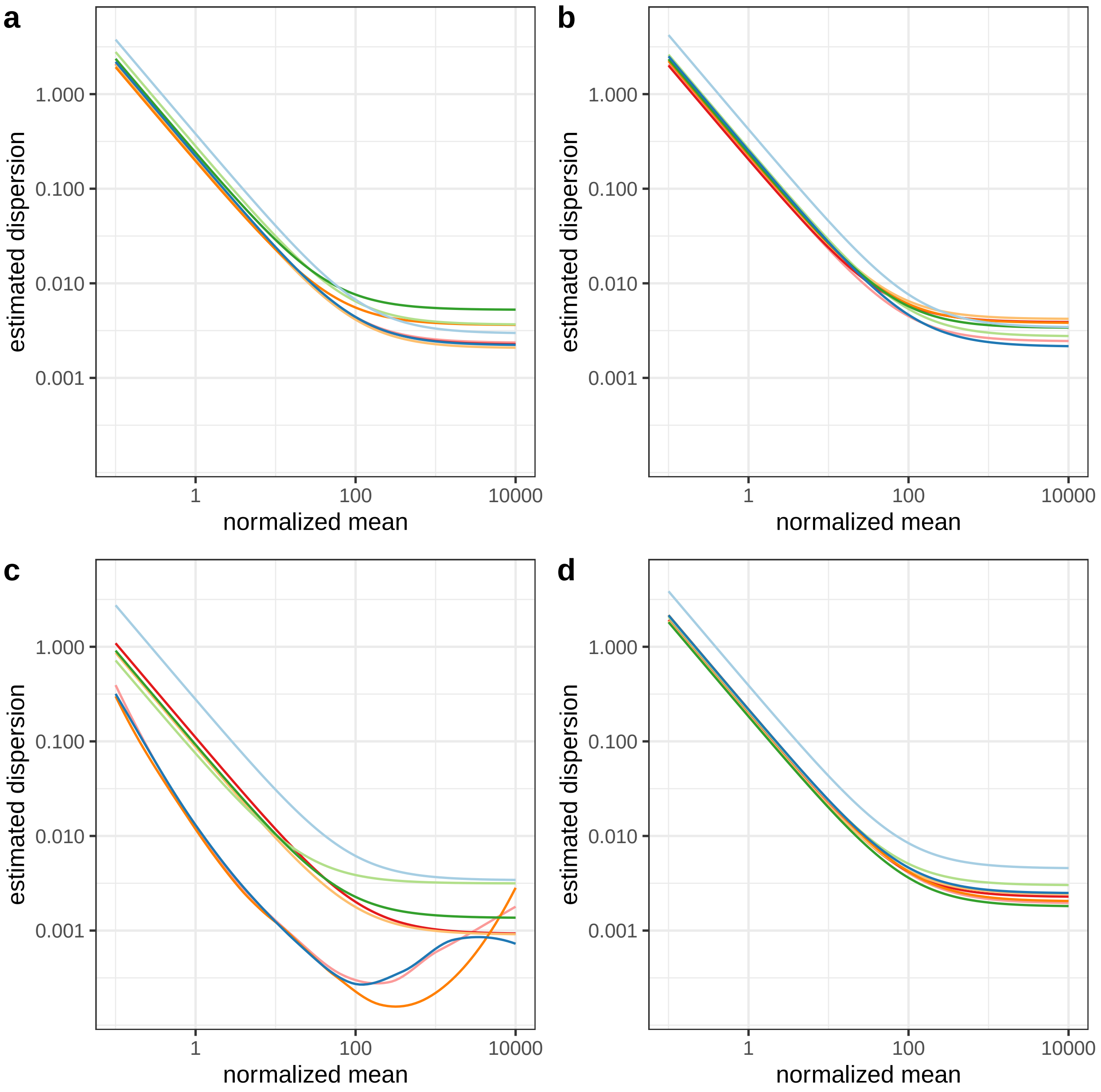


Supplementary Fig. S43: Estimated mean-dispersion functions from DrSeq analysis of RNA-seq datasets (ENCODE) with RBP targets (a) AQR, (b) SRSF1, (c) CPSF6, and (d) SF3B4. Each line corresponds to an ATR event type from Fig. S1.

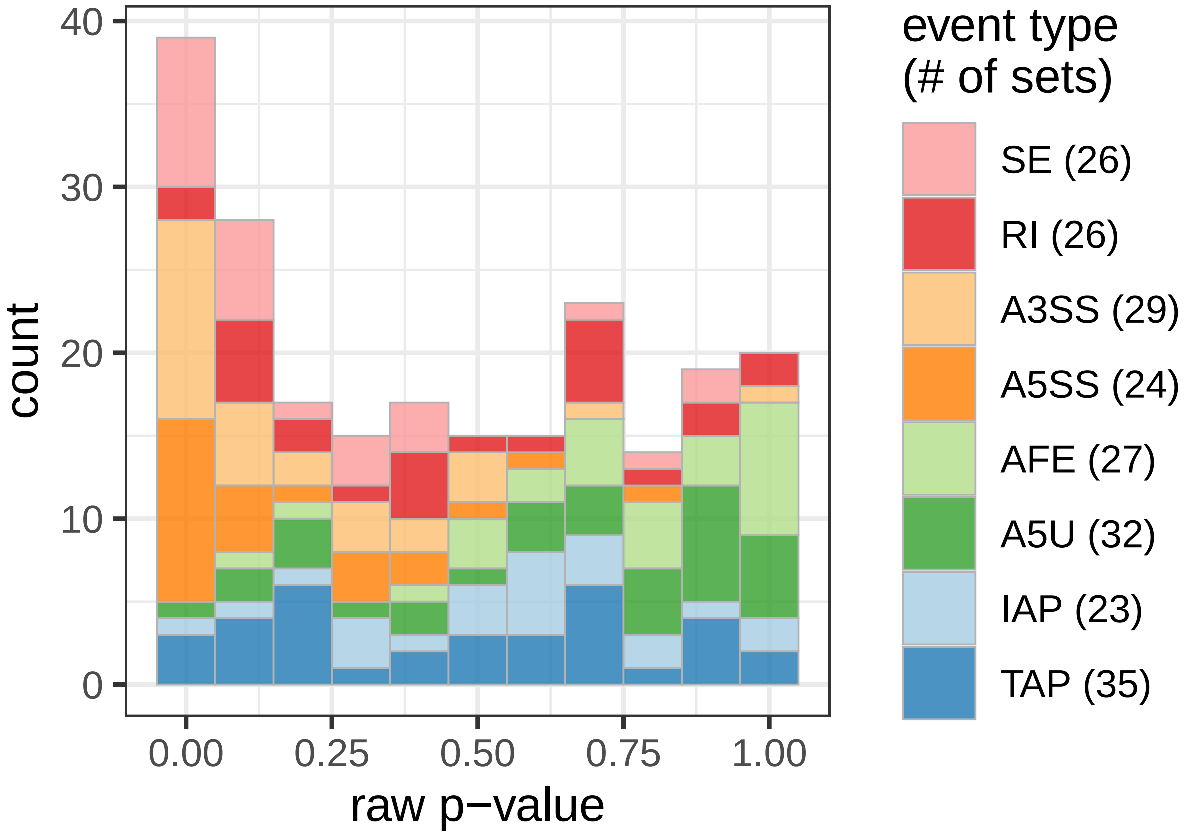


Supplementary Fig. S44: Histogram of raw p-values from the application of SURF discovery module to 222 target transcript sets. The transcript sets are colored with the associated ATR event types. For each event type, the number of transcript sets is reported in parentheses in the color legend.

### Supplementary Tables

Supplementary Table S1: Summary of DrSeq results for the differential skipping of the 2nd last exon in the RHOA gene. Counts across replicates of each condition are separated by semicolons. Fig. S12 displays a genome browser view of the normalized RNA-seq read coverage.

| | DrSeq results |
|---:|---|
| gene identifier | ENSG00000067560 |
| gene name | RHOA |
| event defining transcript identifier | ENST00000418115 |
| chromosome name | chr3 |
| strand | - |
| skipping exon (start-end) | 49,398,361-49,398,499 |
| event body read counts (condition 1) | 142; 111; 98 |
| event body read counts (condition 2) | 265; 27; 65 |
| gene read counts (condition 1) | 17,260; 17,203; 16,399 |
| gene read counts (condition 2) | 30,915; 17,744; 23,840 |
| base normalized read counts | 115.222 |
| estimated dispersion parameter | 0.150 |
| estimated REU coefficient (condition 1) | 17.336 |
| estimated REU coefficient (condition 2) | 14.765 |
| $\log_2$ REU fold change (condition 2 vs 1) | -0.487 |
| significance of differential REU (p-value) | 0.129 |
| adjusted p-value | 0.773 |

Supplementary Table S2: Summary of rMATS results for the differential skipping of the 2nd last exon in the RHOA gene. Counts across replicates of each condition are separated by semicolons. Fig. S12 displays a genome browser view of the normalized RNA-seq read coverage.

| | rMATS results |
|---:|---|
| gene identifier | ENSG00000067560 |
| gene name | RHOA |
| chromosome name | chr3 |
| strand | - |
| skipping exon (start-end) | 49,398,360-49,398,499 |
| inclusion junction counts (condition 1) | 210; 146; 139 |
| skipping junction counts (condition 1) | 1594; 1501; 1425 |
| inclusion junction counts (condition 2) | 369; 39; 93 |
| skipping junction counts (condition 2) | 2711; 1613; 2156 |
| length of inclusion form | 239 |
| length of skipping form | 100 |
| significance of splicing difference (p-value) | 2.81e-06 |
| FDR (adjusted p-value) | 1.05e-04 |
| inclusion level (condition 1) | 0.052; 0.039; 0.039 |
| inclusion level (condition 2) | 0.054; 0.01; 0.018 |
| inclusion level difference (condition 1 vs. 2) | 0.016 |

Supplementary Table S3: Number of false positives of the SURF analysis module 2 (functional association testing). The analysis module 2 was repeated for each of 100 replicates with permuted differential ATR labels. The average numbers of false positives, with target FDR threshold of 0.01, 0.05, and 0.1 after multiplicity correction across all RBPs and ATR event types, are reported. For each FDR threshold, the standard deviation (std. dev.) of the 100 observed numbers is listed besides.

| FDR threshold | average # of false positives | std. dev. |
|---|---|---|
| 0.01 | 0.73 | 0.886 |
| 0.05 | 3.88 | 2.38 |
| 0.10 | 8.52 | 3.80 |

Supplementary Table S4: The fold change in MPK among four genomic sets – genes, spanning regions of ATR events, location features, and SURF-inferred features – from the ICGC (non-US) projects. For each pair of genomic sets, the observed fold change for the genomic set in column over the genomic set in row is listed in the table. The corresponding p-value of exact Poisson test for each fold change is reported in parentheses.

| | span | feature | inferred |
|---|---|---|---|
| gene | 1.21 ($1.05 \times 10^{-2}$) | 1.24 ($4.20 \times 10^{-2}$) | 1.50 ($3.00 \times 10^{-4}$) |
| span | / | 1.02 ($3.19 \times 10^{-4}$) | 1.23 ($6.12 \times 10^{-3}$) |
| feature | / | / | 1.20 ($3.68 \times 10^{-3}$) |

Supplementary Table S5: The fold change in MPK among four genomic sets – genes, spanning regions of ATR events, location features, and SURF-inferred features – from the TCGA project. For each pair of genomic sets, the observed fold change for the genomic set in column over the genomic set in row is listed in the table. The corresponding p-value of exact Poisson test for each fold change is reported in parentheses.

| | span | feature | inferred |
|---|---|---|---|
| gene | 0.99 (0.629) | 1.075 ($1.22 \times 10^{-5}$) | 1.65 ($2.23 \times 10^{-3}$) |
| span | / | 1.08 ($1.05 \times 10^{-12}$) | 1.66 ($8.52 \times 10^{-6}$) |
| feature | / | / | 1.151 ($3.88 \times 10^{-9}$) |

Supplementary Table S6: The activity levels of SURF-identified transcript sets in TCGA LAML and GTEx whole blood samples. SURF-inferred transcription sets corresponding to 222 RBP $\times$ event combinations, from which 15 sets with the number of transcripts (size) fewer than 10 are excluded. For each transcript set, the p-value of differential activity between normal (GTEx) and tumor (TCGA) samples, after multiplicity correction using BH procedure, are reported (adj. p-value). The significance codes – $***$: adjusted p-value$<$0.001, $**$: adjusted p-value $<$0.01, $*$: adjusted p-value $<$0.05, $\cdot$: adjusted p-value $<$0.1 – are also marked.

| | factor | event | size | AUC(GTEx) | AUC(TCGA) | adj. p-value | significance |
|---|---|---|---|---|---|---|---|
| 1 | AQR | SE | 481 | 0.1500 | 0.1716 | 0.4050 | |
| 2 | EFTUD2 | SE | 20 | 0.2579 | 0.2476 | 1.0000 | |
| 3 | EIF3G | SE | 92 | 0.1676 | 0.2204 | 0.3863 | |
| 4 | FXR1 | SE | 513 | 0.1085 | 0.1548 | 0.0000 | *** |
| 5 | HNRNPC | SE | 931 | 0.0783 | 0.1103 | 0.0028 | ** |
| 6 | HNRNPL | SE | 25 | 0.0867 | 0.1855 | 0.0727 | . |
| 7 | HNRNPM | SE | 11 | 0.0000 | 0.0915 | 0.2556 | |
| 8 | HNRNPU | SE | 146 | 0.1513 | 0.1918 | 0.8046 | |
| 9 | IGF2BP2 | SE | 403 | 0.0947 | 0.0957 | 1.0000 | |
| 10 | LIN28B | SE | 674 | 0.0747 | 0.0959 | 0.5352 | |
| 11 | MATR3 | SE | 21 | 0.0116 | 0.0343 | 0.6899 | |
| 12 | NONO | SE | 29 | 0.0824 | 0.0858 | 1.0000 | |
| 13 | PRPF8 | SE | 187 | 0.1812 | 0.2334 | 0.3280 | |
| 14 | PUM1 | SE | 378 | 0.0960 | 0.1146 | 0.2789 | |
| 15 | RBM15 | SE | 1050 | 0.0989 | 0.1279 | 0.1083 | |
| 16 | SF3B4 | SE | 119 | 0.0948 | 0.1281 | 0.3870 | |
| 17 | SMNDC1 | SE | 17 | 0.0936 | 0.2172 | 0.3609 | |
| 18 | SRSF1 | SE | 671 | 0.0987 | 0.1352 | 0.0000 | *** |
| 19 | TARDBP | SE | 114 | 0.1117 | 0.1335 | 0.4724 | |
| 20 | TIA1 | SE | 76 | 0.0900 | 0.1219 | 0.9612 | |
| 21 | TRA2A | SE | 118 | 0.1002 | 0.1223 | 0.8093 | |
| 22 | U2AF1 | SE | 186 | 0.0990 | 0.1442 | 0.2442 | |
| 23 | U2AF2 | SE | 625 | 0.1005 | 0.1285 | 0.0795 | . |
| 24 | UCHL5 | SE | 244 | 0.0955 | 0.1167 | 0.7666 | |
| 25 | XRN2 | SE | 135 | 0.0914 | 0.1078 | 0.8046 | |
| 26 | AQR | RI | 865 | 0.2032 | 0.2011 | 0.9678 | |
| 27 | CPSF6 | RI | 31 | 0.1585 | 0.1795 | 1.0000 | |
| 28 | EFTUD2 | RI | 378 | 0.2356 | 0.2437 | 0.8578 | |
| 29 | FXR1 | RI | 479 | 0.1730 | 0.2159 | 0.0210 | * |
| 30 | HNRNPC | RI | 314 | 0.1272 | 0.1609 | 0.4680 | |
| 31 | HNRNPK | RI | 33 | 0.1636 | 0.1741 | 0.9612 | |
| 32 | HNRNPU | RI | 59 | 0.1400 | 0.1544 | 1.0000 | |
| 33 | IGF2BP2 | RI | 321 | 0.1668 | 0.1888 | 0.1623 | |
| 34 | KHSRP | RI | 38 | 0.1813 | 0.2356 | 0.9612 | |
| 35 | LIN28B | RI | 53 | 0.1300 | 0.1712 | 0.8046 | |

Supplementary Table S6: The active level of SURF-identified transcript sets in TCGA LAML and GTEx whole blood samples (continued).

| | factor | event | size | AUC(GTEx) | AUC(TCGA) | adj. p-value | significance |
|---|---|---|---|---|---|---|---|
| 36 | LSM11 | RI | 75 | 0.2315 | 0.2434 | 0.9400 | |
| 37 | METAP2 | RI | 38 | 0.3286 | 0.3696 | 0.7725 | |
| 38 | NCBP2 | RI | 111 | 0.1727 | 0.2010 | 0.4749 | |
| 39 | PRPF8 | RI | 500 | 0.2050 | 0.2226 | 0.3788 | |
| 40 | PUM1 | RI | 218 | 0.1578 | 0.1749 | 0.5352 | |
| 41 | RBM15 | RI | 754 | 0.1986 | 0.2077 | 0.7666 | |
| 42 | SF3B4 | RI | 362 | 0.2051 | 0.2322 | 0.4618 | |
| 43 | SMNDC1 | RI | 68 | 0.2721 | 0.2790 | 1.0000 | |
| 44 | SRSF1 | RI | 596 | 0.1648 | 0.1837 | 0.3956 | |
| 45 | TARDBP | RI | 137 | 0.1696 | 0.2124 | 0.5935 | |
| 46 | TIA1 | RI | 122 | 0.1524 | 0.1816 | 0.9612 | |
| 47 | TRA2A | RI | 180 | 0.2059 | 0.2101 | 1.0000 | |
| 48 | U2AF2 | RI | 354 | 0.2026 | 0.2193 | 0.8994 | |
| 49 | UCHL5 | RI | 190 | 0.1880 | 0.2174 | 0.7725 | |
| 50 | XRN2 | RI | 168 | 0.1696 | 0.1704 | 1.0000 | |
| 51 | APOBEC3C | A3SS | 42 | 0.2495 | 0.2693 | 0.8994 | |
| 52 | AQR | A3SS | 1167 | 0.2265 | 0.2625 | 0.0000 | *** |
| 53 | EFTUD2 | A3SS | 179 | 0.2383 | 0.2685 | 0.3870 | |
| 54 | EIF3G | A3SS | 38 | 0.1975 | 0.2643 | 0.8994 | |
| 55 | FXR1 | A3SS | 563 | 0.1678 | 0.2276 | 0.0000 | *** |
| 56 | HNRNPC | A3SS | 482 | 0.1513 | 0.1799 | 0.3788 | |
| 57 | HNRNPK | A3SS | 40 | 0.2216 | 0.1848 | 0.5054 | |
| 58 | HNRNPU | A3SS | 97 | 0.1862 | 0.2707 | 0.2442 | |
| 59 | IGF2BP2 | A3SS | 261 | 0.1906 | 0.2154 | 0.2442 | |
| 60 | KHSRP | A3SS | 40 | 0.1532 | 0.3063 | 0.0266 | * |
| 61 | LIN28B | A3SS | 325 | 0.1470 | 0.1678 | 0.7725 | |
| 62 | NCBP2 | A3SS | 23 | 0.1926 | 0.2266 | 0.9612 | |
| 63 | NONO | A3SS | 11 | 0.2721 | 0.0989 | 0.4680 | |
| 64 | PRPF8 | A3SS | 297 | 0.2315 | 0.2574 | 0.4791 | |
| 65 | PUM1 | A3SS | 348 | 0.1491 | 0.2060 | 0.0000 | *** |
| 66 | PUS1 | A3SS | 80 | 0.2267 | 0.2569 | 0.6655 | |
| 67 | RBM15 | A3SS | 888 | 0.1885 | 0.2373 | 0.0000 | *** |
| 68 | RBM22 | A3SS | 12 | 0.2311 | 0.3195 | 0.5352 | |
| 69 | SF3B4 | A3SS | 294 | 0.2108 | 0.2646 | 0.0000 | *** |
| 70 | SMNDC1 | A3SS | 60 | 0.2375 | 0.3135 | 0.3008 | |
| 71 | SRSF1 | A3SS | 587 | 0.1663 | 0.2060 | 0.0000 | *** |
| 72 | TARDBP | A3SS | 191 | 0.2384 | 0.2809 | 0.2408 | |
| 73 | TIA1 | A3SS | 70 | 0.2025 | 0.2338 | 0.7321 | |
| 74 | TRA2A | A3SS | 181 | 0.1869 | 0.2003 | 0.7725 | |
| 75 | U2AF1 | A3SS | 52 | 0.2194 | 0.2598 | 0.6877 | |

Supplementary Table S6: The active level of SURF-identified transcript sets in TCGA LAML and GTEx whole blood samples (continued).

| | factor | event | size | AUC(GTEx) | AUC(TCGA) | adj. p-value | significance |
|---|---|---|---|---|---|---|---|
| 76 | U2AF2 | A3SS | 621 | 0.1568 | 0.1915 | 0.0173 | * |
| 77 | UCHL5 | A3SS | 231 | 0.1610 | 0.1910 | 0.1835 | |
| 78 | XRN2 | A3SS | 167 | 0.2072 | 0.1952 | 0.8994 | |
| 79 | APOBEC3C | A5SS | 30 | 0.2198 | 0.3393 | 0.0795 | . |
| 80 | AQR | A5SS | 930 | 0.1914 | 0.2389 | 0.0000 | *** |
| 81 | EFTUD2 | A5SS | 59 | 0.2082 | 0.2213 | 0.9612 | |
| 82 | FXR1 | A5SS | 425 | 0.1552 | 0.2146 | 0.0000 | *** |
| 83 | HNRNPC | A5SS | 370 | 0.1312 | 0.1827 | 0.0131 | * |
| 84 | HNRNPK | A5SS | 21 | 0.1444 | 0.1744 | 0.8104 | |
| 85 | HNRNPU | A5SS | 61 | 0.1576 | 0.1661 | 1.0000 | |
| 86 | IGF2BP2 | A5SS | 286 | 0.1543 | 0.1769 | 0.3076 | |
| 87 | LIN28B | A5SS | 216 | 0.1819 | 0.2068 | 0.7666 | |
| 88 | NCBP2 | A5SS | 30 | 0.1739 | 0.2399 | 0.2373 | |
| 89 | NONO | A5SS | 12 | 0.1208 | 0.2069 | 0.4071 | |
| 90 | PRPF8 | A5SS | 280 | 0.2093 | 0.2736 | 0.0000 | *** |
| 91 | PUM1 | A5SS | 239 | 0.1525 | 0.1870 | 0.1212 | |
| 92 | RBM15 | A5SS | 688 | 0.1669 | 0.2158 | 0.0000 | *** |
| 93 | SF3B4 | A5SS | 120 | 0.2075 | 0.2385 | 0.3076 | |
| 94 | SMNDC1 | A5SS | 19 | 0.3087 | 0.3689 | 0.9283 | |
| 95 | SRSF1 | A5SS | 531 | 0.1547 | 0.1969 | 0.0000 | *** |
| 96 | TARDBP | A5SS | 81 | 0.2150 | 0.2487 | 0.8233 | |
| 97 | TIA1 | A5SS | 57 | 0.1302 | 0.2118 | 0.1475 | |
| 98 | TRA2A | A5SS | 166 | 0.1576 | 0.1940 | 0.3552 | |
| 99 | U2AF2 | A5SS | 481 | 0.1488 | 0.2010 | 0.0028 | ** |
| 100 | UCHL5 | A5SS | 172 | 0.1226 | 0.1480 | 0.7666 | |
| 101 | XRN2 | A5SS | 136 | 0.1655 | 0.1860 | 0.6306 | |
| 102 | AQR | AFE | 481 | 0.0671 | 0.0548 | 0.9612 | |
| 103 | CPSF6 | AFE | 52 | 0.0392 | 0.0720 | 1.0000 | |
| 104 | EFTUD2 | AFE | 93 | 0.1118 | 0.1156 | 1.0000 | |
| 105 | FMR1 | AFE | 36 | 0.1047 | 0.1221 | 0.9678 | |
| 106 | FXR1 | AFE | 625 | 0.0584 | 0.0638 | 1.0000 | |
| 107 | HNRNPC | AFE | 669 | 0.0649 | 0.0802 | 1.0000 | |
| 108 | HNRNPU | AFE | 81 | 0.1017 | 0.0924 | 0.9612 | |
| 109 | IGF2BP2 | AFE | 683 | 0.0662 | 0.0672 | 1.0000 | |
| 110 | LIN28B | AFE | 692 | 0.0612 | 0.0693 | 1.0000 | |
| 111 | LSM11 | AFE | 16 | 0.1784 | 0.1505 | 0.9678 | |
| 112 | METAP2 | AFE | 48 | 0.1423 | 0.1173 | 0.9023 | |
| 113 | NCBP2 | AFE | 96 | 0.0872 | 0.1228 | 0.4050 | |
| 114 | NONO | AFE | 69 | 0.1176 | 0.1686 | 0.5352 | |
| 115 | PRPF8 | AFE | 213 | 0.0949 | 0.1029 | 1.0000 | |

Supplementary Table S6: The active level of SURF-identified transcript sets in TCGA LAML and GTEx whole blood samples (continued).

| | factor | event | size | AUC(GTEx) | AUC(TCGA) | adj. p-value | significance |
|---|---|---|---|---|---|---|---|
| 116 | PUM1 | AFE | 453 | 0.0532 | 0.0656 | 1.0000 | |
| 117 | RBM15 | AFE | 1239 | 0.0584 | 0.0616 | 1.0000 | |
| 118 | SF3B4 | AFE | 117 | 0.0710 | 0.0832 | 1.0000 | |
| 119 | SMNDC1 | AFE | 26 | 0.2601 | 0.3352 | 0.9799 | |
| 120 | SRSF1 | AFE | 689 | 0.0581 | 0.0679 | 1.0000 | |
| 121 | TARDBP | AFE | 278 | 0.0520 | 0.0690 | 1.0000 | |
| 122 | TIA1 | AFE | 85 | 0.1104 | 0.1541 | 0.8994 | |
| 123 | TRA2A | AFE | 380 | 0.0827 | 0.0988 | 1.0000 | |
| 124 | U2AF1 | AFE | 28 | 0.1222 | 0.1658 | 0.9530 | |
| 125 | U2AF2 | AFE | 374 | 0.0662 | 0.0735 | 1.0000 | |
| 126 | UCHL5 | AFE | 382 | 0.0813 | 0.0970 | 0.9612 | |
| 127 | XRN2 | AFE | 244 | 0.1002 | 0.1193 | 0.8147 | |
| 128 | AQR | A5U | 144 | 0.0827 | 0.0600 | 0.3956 | |
| 129 | EFTUD2 | A5U | 24 | 0.1208 | 0.1263 | 1.0000 | |
| 130 | EIF3G | A5U | 133 | 0.0944 | 0.1002 | 1.0000 | |
| 131 | FMR1 | A5U | 66 | 0.1315 | 0.1239 | 1.0000 | |
| 132 | FUS | A5U | 61 | 0.1611 | 0.1997 | 1.0000 | |
| 133 | FXR1 | A5U | 823 | 0.0869 | 0.1056 | 1.0000 | |
| 134 | GEMIN5 | A5U | 335 | 0.1080 | 0.1219 | 0.9799 | |
| 135 | GRWD1 | A5U | 243 | 0.0980 | 0.1078 | 1.0000 | |
| 136 | HNRNPC | A5U | 918 | 0.0659 | 0.0864 | 1.0000 | |
| 137 | IGF2BP1 | A5U | 44 | 0.1583 | 0.2600 | 0.0507 | . |
| 138 | IGF2BP2 | A5U | 1318 | 0.0768 | 0.0827 | 1.0000 | |
| 139 | LIN28B | A5U | 1113 | 0.0683 | 0.0786 | 1.0000 | |
| 140 | METAP2 | A5U | 76 | 0.1074 | 0.1074 | 1.0000 | |
| 141 | NCBP2 | A5U | 502 | 0.0882 | 0.0827 | 0.9612 | |
| 142 | NONO | A5U | 63 | 0.1192 | 0.1182 | 1.0000 | |
| 143 | PUM1 | A5U | 1026 | 0.0818 | 0.0836 | 1.0000 | |
| 144 | RBM15 | A5U | 1687 | 0.0541 | 0.0686 | 1.0000 | |
| 145 | SERBP1 | A5U | 100 | 0.0869 | 0.0903 | 1.0000 | |
| 146 | SF3B4 | A5U | 38 | 0.1020 | 0.0718 | 0.8233 | |
| 147 | SND1 | A5U | 113 | 0.1456 | 0.2063 | 0.9612 | |
| 148 | SRSF1 | A5U | 1295 | 0.0699 | 0.0850 | 0.5988 | |
| 149 | SUPV3L1 | A5U | 132 | 0.0828 | 0.0774 | 1.0000 | |
| 150 | TARDBP | A5U | 184 | 0.0740 | 0.0848 | 0.9612 | |
| 151 | TRA2A | A5U | 796 | 0.0954 | 0.1123 | 0.8046 | |
| 152 | U2AF2 | A5U | 401 | 0.0781 | 0.1005 | 0.9612 | |
| 153 | UCHL5 | A5U | 885 | 0.0774 | 0.1032 | 0.5352 | |
| 154 | XRN2 | A5U | 284 | 0.0726 | 0.0747 | 1.0000 | |
| 155 | CPSF6 | IAP | 94 | 0.0592 | 0.0804 | 1.0000 | |

Supplementary Table S6: The active level of SURF-identified transcript sets in TCGA LAML and GTEx whole blood samples (continued).

| | factor | event | size | AUC(GTEx) | AUC(TCGA) | adj. p-value | significance |
|---|---|---|---|---|---|---|---|
| 156 | EIF3G | IAP | 17 | 0.0000 | 0.0232 | 0.9612 | |
| 157 | FXR1 | IAP | 587 | 0.0241 | 0.0451 | 0.8994 | |
| 158 | HNRNPC | IAP | 377 | 0.0280 | 0.0631 | 0.6306 | |
| 159 | HNRNPU | IAP | 38 | 0.0640 | 0.0574 | 1.0000 | |
| 160 | IGF2BP2 | IAP | 642 | 0.0319 | 0.0350 | 0.9799 | |
| 161 | LIN28B | IAP | 803 | 0.0423 | 0.0524 | 1.0000 | |
| 162 | LSM11 | IAP | 73 | 0.0748 | 0.0895 | 0.9612 | |
| 163 | NCBP2 | IAP | 28 | 0.0595 | 0.0577 | 1.0000 | |
| 164 | NONO | IAP | 20 | 0.1810 | 0.1652 | 1.0000 | |
| 165 | PRPF8 | IAP | 73 | 0.0676 | 0.1208 | 0.8994 | |
| 166 | PUM1 | IAP | 182 | 0.0318 | 0.0433 | 0.8994 | |
| 167 | RBM15 | IAP | 1009 | 0.0282 | 0.0506 | 0.9612 | |
| 168 | SF3B4 | IAP | 129 | 0.0379 | 0.0565 | 0.6535 | |
| 169 | SRSF1 | IAP | 552 | 0.0295 | 0.0515 | 0.2373 | |
| 170 | TARDBP | IAP | 196 | 0.0213 | 0.0533 | 0.9612 | |
| 171 | TIA1 | IAP | 162 | 0.0376 | 0.0535 | 0.9612 | |
| 172 | TRA2A | IAP | 326 | 0.0281 | 0.0568 | 0.7441 | |
| 173 | U2AF2 | IAP | 565 | 0.0377 | 0.0577 | 0.8093 | |
| 174 | UCHL5 | IAP | 277 | 0.0554 | 0.0906 | 0.3870 | |
| 175 | XRN2 | IAP | 82 | 0.0627 | 0.0815 | 0.9612 | |
| 176 | APOBEC3C | TAP | 39 | 0.1083 | 0.1875 | 0.4680 | |
| 177 | AQR | TAP | 589 | 0.0350 | 0.0516 | 0.1475 | |
| 178 | CPSF6 | TAP | 194 | 0.0665 | 0.0877 | 1.0000 | |
| 179 | EIF3G | TAP | 110 | 0.0844 | 0.1276 | 0.8899 | |
| 180 | FMR1 | TAP | 26 | 0.0975 | 0.1037 | 1.0000 | |
| 181 | FUS | TAP | 67 | 0.1635 | 0.2139 | 0.9400 | |
| 182 | FXR1 | TAP | 709 | 0.0596 | 0.0771 | 1.0000 | |
| 183 | GRWD1 | TAP | 365 | 0.0664 | 0.0968 | 0.3552 | |
| 184 | HNRNPC | TAP | 1941 | 0.0505 | 0.0680 | 0.9612 | |
| 185 | HNRNPL | TAP | 18 | 0.1735 | 0.1623 | 1.0000 | |
| 186 | HNRNPU | TAP | 184 | 0.1019 | 0.1360 | 0.9678 | |
| 187 | IGF2BP1 | TAP | 53 | 0.0870 | 0.0647 | 0.8994 | |
| 188 | IGF2BP2 | TAP | 1339 | 0.0644 | 0.0735 | 0.3280 | |
| 189 | KHDRBS1 | TAP | 12 | 0.1244 | 0.1669 | 0.8994 | |
| 190 | LIN28B | TAP | 1792 | 0.0512 | 0.0683 | 0.9612 | |
| 191 | METAP2 | TAP | 104 | 0.1598 | 0.1805 | 0.9307 | |
| 192 | NCBP2 | TAP | 91 | 0.0563 | 0.0564 | 1.0000 | |
| 193 | NONO | TAP | 18 | 0.0945 | 0.0112 | 0.4050 | |
| 194 | PHF6 | TAP | 14 | 0.1096 | 0.1047 | 1.0000 | |
| 195 | PPIL4 | TAP | 32 | 0.0099 | 0.1065 | 0.8093 | |

Supplementary Table S6: The active level of SURF-identified transcript sets in TCGA LAML and GTEx whole blood samples (continued).

| | factor | event | size | AUC(GTEx) | AUC(TCGA) | adj. p-value | significance |
|---|---|---|---|---|---|---|---|
| 196 | PUM1 | TAP | 611 | 0.0514 | 0.0595 | 0.6877 | |
| 197 | PUS1 | TAP | 102 | 0.0378 | 0.0544 | 0.9612 | |
| 198 | RBM15 | TAP | 1563 | 0.0425 | 0.0650 | 0.6243 | |
| 199 | SRSF1 | TAP | 582 | 0.0532 | 0.0566 | 0.9612 | |
| 200 | TARDBP | TAP | 160 | 0.0628 | 0.0908 | 0.6090 | |
| 201 | TIA1 | TAP | 255 | 0.1114 | 0.1407 | 0.9678 | |
| 202 | TRA2A | TAP | 436 | 0.0580 | 0.0696 | 0.5935 | |
| 203 | U2AF1 | TAP | 14 | 0.0634 | 0.1077 | 1.0000 | |
| 204 | U2AF2 | TAP | 886 | 0.0528 | 0.0742 | 0.2746 | |
| 205 | UCHL5 | TAP | 728 | 0.0574 | 0.0788 | 0.5352 | |
| 206 | UPF1 | TAP | 111 | 0.1202 | 0.1000 | 0.8578 | |
| 207 | XRN2 | TAP | 170 | 0.0609 | 0.0804 | 0.5935 | |

Supplementary Table S7: Summary of the numbers (and the percentages) of retained genes, transcripts, and RNA-seq reads (in TPM), averaged across all samples. The filtering strategies (first column) are defined with isoform relative abundance (in percent, IP) or TPM. The first row presents the statistics for the complete set of genome annotation from GENCODE version 24. Rows 2-6 represent pre-filtering strategies with different thresholding settings. The percentages of retained genes, transcripts, and RNA-seq reads (in TPM) are listed in parentheses.

| | #gene | #transcripts | Sum of TPM |
|---|---|---|---|
| Complete set | 60,554 | 200,094 | 1,000,000 |
| IP$\geq$ 5% | 40,972 (67.7%) | 147,963 (73.9%) | 991,810.4 (99.2%) |
| IP$\geq$ 10% | 40,972 (67.7%) | 134,063 (67.0%) | 980,396.0 (98.0%) |
| IP$\geq$ 25% | 40,972 (67.7%) | 108,601 (54.3%) | 937,847.9 (93.8%) |
| TPM$\geq$ 1% | 17,767 (29.3%) | 82,535 (41.2%) | 998,544.9 (99.9%) |
| IP$\geq$ 5% & TPM$\geq$ 1% | 17,767 (29.3%) | 74,733 (37.3%) | 990,518.9 (99.1%) |
